# Overlapping representations of food and social stimuli in VTA dopamine neurons

**DOI:** 10.1101/2023.05.17.541104

**Authors:** Lindsay Willmore, Adelaide R. Minerva, Ben Engelhard, Malavika Murugan, Brenna McMannon, Nirja Oak, Stephan Y. Thiberge, Catherine J. Peña, Ilana B. Witten

## Abstract

Dopamine neurons of the ventral tegmental area (VTA^DA^) respond to food and social stimuli and contribute to both forms of motivation. However, it is unclear if the same or different VTA^DA^ neurons encode these different stimuli. To address this question, we performed 2-photon calcium imaging in mice presented with food and conspecifics, and found statistically significant overlap in the populations responsive to both stimuli. Both hunger and opposite-sex social experience further increased the proportion of neurons that respond to both stimuli, implying that modifying motivation for one stimulus affects responses to both stimuli. In addition, single-nucleus RNA sequencing revealed significant co-expression of feeding- and social-hormone related genes in individual VTA^DA^ neurons. Taken together, our functional and transcriptional data suggest overlapping VTA^DA^ populations underlie food and social motivation.

## Introduction

Dopaminergic neurons in the midbrain are essential for motivated behaviors mediated by a range of unconditioned stimuli, including both palatable rewards (Schultz, Dayan, and Montague 1997; Cohen et al. 2012; Fernandes et al. 2020; Grove et al. 2022; Martel and Fantino 1996; Mazzone et al. 2020) and social stimuli (Gunaydin et al. 2014; Willmore et al. 2022; Solié et al. 2021; Louilot et al. 1991; Bariselli et al. 2018). However, whether the same or different dopamine subpopulations encode such different unconditioned stimuli is unclear. If separate subpopulations of dopamine neurons encode social and food stimuli, that would suggest that at the level of the midbrain, specialized and parallel circuits support these different motivations (similar to findings in cortex; (Jennings et al. 2019; Isaac et al. 2023). Overlapping populations would instead suggest these different motivations are encoded in a “common currency” in the midbrain dopamine system (Harrison, von Neumann, and Morgenstern 1945; McNamara and Houston 1986; Landreth and Bickle 2008; Levy and Glimcher 2012), which could support the ultimate role of any decision system in comparing different outcomes against one another in order to choose which decision to make.

While classic work on dopamine neurons in the ventral tegmental area (VTA^DA^) highlighted homogenous responses to reward (and to errors in the prediction of reward; (Houk, Davis, and Beiser 1995; Cohen et al. 2012; Schultz, Dayan, and Montague 1997; Steinberg et al. 2013), more recent work has identified heterogeneity at the single cell level. For example, VTA^DA^ neurons can have heterogeneous and specialized responses to task variables during complex behavior (Parker et al. 2016; R. S. Lee et al. 2019; Choi et al. 2020; Lerner et al. 2015; Collins and Saunders 2020; Verharen, Zhu, and Lammel 2020; Hassan and Benarroch 2015; Marinelli and McCutcheon 2014; Kremer et al. 2020; Howe and Dombeck 2016; Anderegg, Poulin, and Awatramani 2015; Barter et al. 2015; Cai et al. 2020; Hamid, Frank, and Moore 2021; Wei, Mohebi, and Berke 2021; Zolin et al. 2021; Dabney et al. 2020; Engelhard et al. 2019), while at the same time having relatively homogenous responses to food or water (Eshel et al. 2016; Cohen et al. 2012; Kremer et al. 2020; Engelhard et al. 2019; Schultz, Dayan, and Montague 1997).

The fact that VTA^DA^ neurons are increasingly appreciated to be functionally, anatomically, and transcriptionally heterogeneous (Lammel et al. 2012; Watabe-Uchida et al. 2012; Phillips et al. 2022; Tiklová et al. 2019; Lammel et al. 2011; Lerner et al. 2015; Beier et al. 2015; Saunders et al. 2018; La Manno et al. 2016; Poulin et al. 2014) may suggest that separate populations encode food and social stimuli. On the other hand, the fact that most VTA^DA^ neurons respond to palatable rewards while some also encode social stimuli (Gunaydin et al. 2014; Solié et al. 2021) may suggest that overlapping populations of neurons might encode these two sets of stimuli.

To answer whether the same or different VTA^DA^ neurons respond to food and social stimuli, we employed 2-photon calcium imaging of VTA^DA^ neurons in mice presented with conspecifics or food, while varying hunger state and social experience. In addition, we performed single-nucleus RNA sequencing (snRNA-seq) to examine the extent of overlap in expression of feeding and social hormone-related genes in VTA^DA^ neurons. We observed statistically significant overlap between food and social neural representations, as well as between feeding- and social-related gene expression, in individual VTA^DA^ neurons. Together, our neural activity and gene expression findings suggest overlapping VTA^DA^ populations underlie these different behavioral motivations.

## Results

### 2-photon calcium imaging of VTA^DA^ responses to food and social stimuli

To gain optical access to dopamine neurons, we implanted a GRIN lens over the VTA in transgenic mice (N=11 male, 9 females) expressing GCaMP6f in DAT+ neurons (Figure 1A). In order to assess individual neuron responses to food or social stimuli, we devised a paradigm in which animals were presented with stimuli via a software-triggered slider while undergoing 2-photon calcium imaging (Figure 1B-C). On each trial, the slider was loaded with one of five different stimulus types: 3 social stimuli (male, estrus female, non-estrus female), palatable food (sweetened condensed milk), or a control empty box (Figure 1D). This set-up allowed us to examine responses to conspecifics and food in a controlled setting while recording from large, stable populations of dopamine neurons longitudinally.

**Figure 1.**
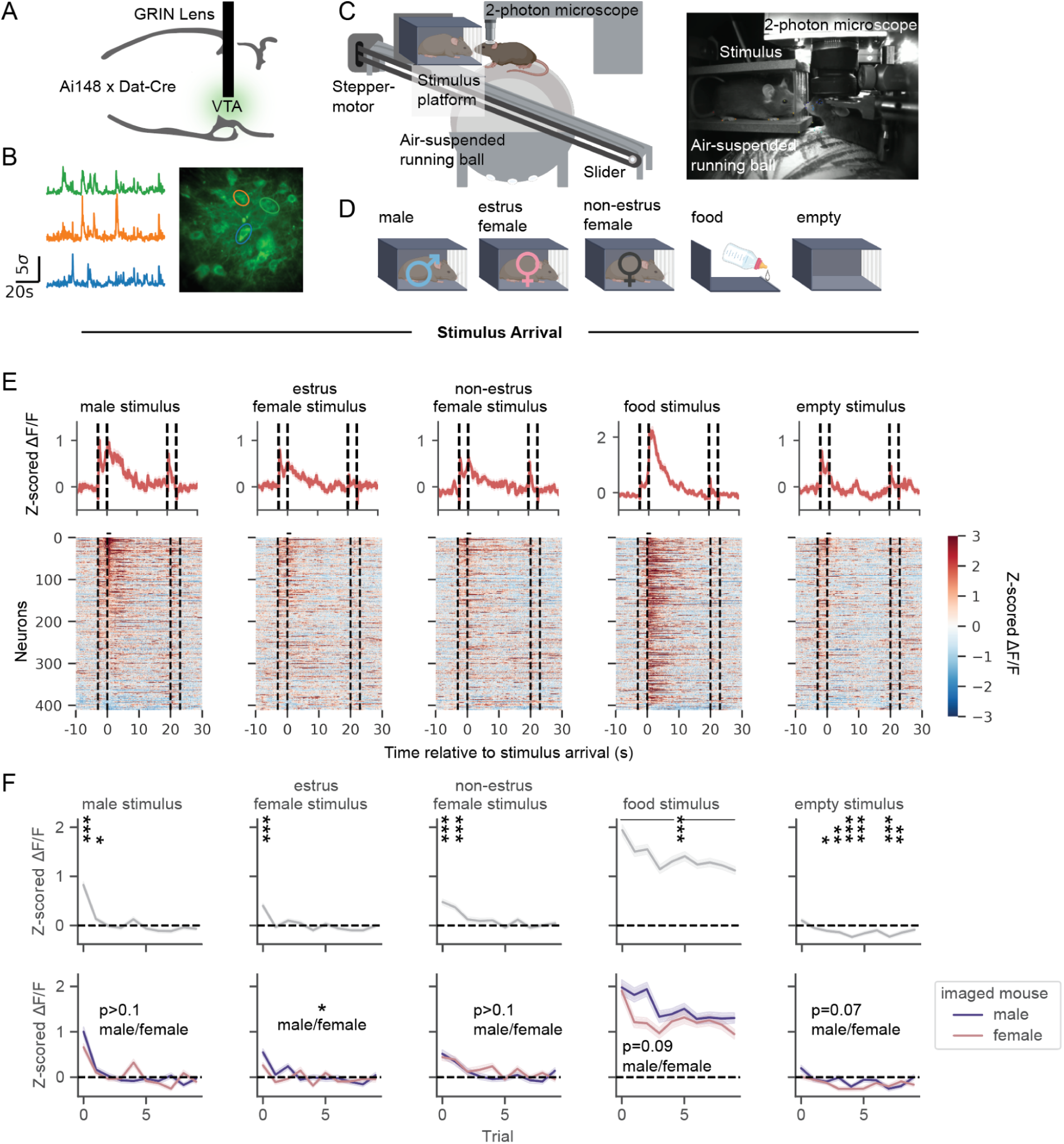
VTA^DA^ neurons respond to the arrival of a conspecific, with strongest responses to males and attenuating responses across trials. (A) Schematic of surgical strategy for GRIN lens insertion over the ventral tegmental area (VTA). (B) Example GCaMP traces from neurons in an example field of view. (C) Schematic of social stimulus and food delivery mechanism to head-fixed mice undergoing two-photon calcium imaging. (D) Stimuli presented during imaging paradigm, including novel conspecifics, food (sweetened condensed milk), and an empty cage. (E) Neural activity to the first trial of each stimulus type on a single baseline imaging session (dotted lines indicate slider movement onset and offset, with arrival at t=0s and departure at t=20s). Neurons plotted in the same order across columns (N=411 neurons, 19 mice). (F) Top: Average activity in the first second after stimulus arrival across all neurons within each trial (1-sample t-test for activity different from 0, after Bonferroni correction for 50 tests). Bottom: Average activity across neurons recorded from male and female imaged mice (generalized estimating equation (GEE) (a linear model that accounts for correlated repeated measurements from each mouse over time) for average activity by trial, imaged mouse sex, and the interaction between trial and imaged mouse sex, N=199 neurons from males, 212 neurons from females). **See Table S1-6 for statistics**. ∗p < 0.05, ∗∗p < 0.01, ∗∗∗p < 0.001. Unless specified, data plotted as mean ± SEM.

We assessed VTA^DA^ responses upon the arrival of the stimulus in front of the imaged animal (Movie S1). VTA^DA^ neurons increased their activity to the first arrival of social and food stimuli but not the empty control stimulus (Figure 1E, N=411 neurons). Food responses remained elevated across subsequent presentations, while social responses were specifically driven by the first few encounters with a novel conspecific (as previously reported in VTA^DA^ neurons: (Gunaydin et al. 2014; Solié et al. 2021; Dai et al. 2022; Robinson, Heien, and Wightman 2002; Damsma et al. 1992), and responses to the empty box became negative across trials (Figure 1F, top). Comparing activity between imaged males and females, we found that activity in response to estrus female arrival was more elevated in imaged males than females (Figure 1F, bottom).

To further characterize social responses across neurons, we clustered responses to social stimulus arrival into profiles with varying magnitudes and durations, all with peak activity immediately following stimulus arrival (Figure S1A-C). After social stimulus arrival, neural responses could also be observed aligned to close contact with the social stimulus, who was not head-fixed and therefore could closely inspect or retreat from the imaged mouse (Figure S2A; contact defined by video recording and automated tracking of key points on the imaged and stimulus mice, see Methods). Responses to social contact were sexually dimorphic. In particular, neurons from imaged females positively responded to males on average, while average responses across male neurons were negative for contact with males and positive for contact with females in estrus (Figure S2B-C).

### Overlap in VTA^DA^ populations responsive to food and social stimuli

We next compared how single VTA^DA^ neurons responded to food versus social stimuli. Within a field of view, we observed examples of neurons responding to the arrival of food, social stimuli, or both stimulus types (Figure 2A-B). Of all imaged neurons, 32% (N=132/411) responded to food but not male arrival, 7% (N=30/411) responded to male but not food arrival, and 14% (N=59/411) responded to both food and male arrival. The fraction of neurons responding to both food and male arrival overlapped significantly, in that it was greater than expected assuming independent food and social populations (Figure 2C, p<0.001 compared to a null distribution constructed based on the recorded fraction of neurons responding to each alone and assuming independence, see Methods). Similarly, when comparing neurons responsive to food and estrus female stimulus arrival, we also observed a significant degree of overlap (Figure 2D, p<0.001). Thus, most socially-responsive DA neurons are also food responsive.

**Figure 2.**
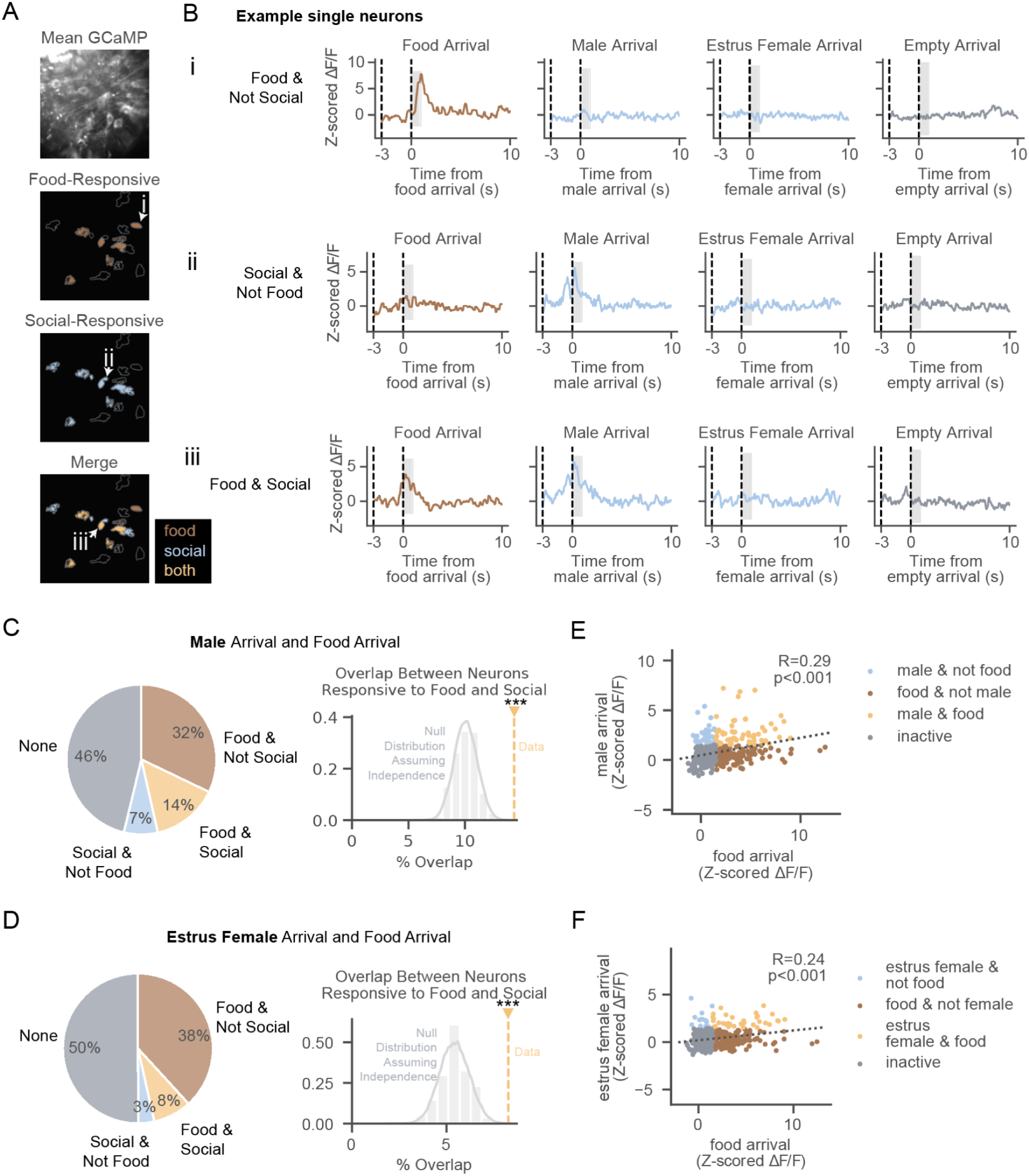
Overlap in VTA^DA^ populations responsive to food and social stimuli. (A) Example field of view with neurons colored by which stimulus type they are responsive to. (N=16 neurons). (B) Examples of neural activity in the 1s after the first presentation of food or social stimuli from a neuron responsive to food, but not social stimuli, arrival (i), a neuron responsive to male, but not food, arrival (ii), and a neuron responsive to both food and male arrival (iii). Slider motor turns on at time -3s and arrives at 0s. (C) Left: Percentage of recorded neurons responsive to food arrival, male arrival, both, or neither. Tuning here and in subsequent panels is based on the mean activity in the first second after arrival (shaded region in B). Right: Overlap compared to null distribution assuming food- and male-responsive neurons are independent samples. (In C-F, N=411 neurons) (D) Same as (C) for estrus female stimulus mice. (E) Correlation between neural activity in response to food arrival and male arrival, colored by selectivity to food arrival, male arrival, or both. (F) Same as (E) for estrus female stimulus mice. **See Table S11 for statistics.** ∗p < 0.05, ∗∗p < 0.01, ∗∗∗p < 0.001. Unless specified, data plotted as mean ± SEM.

If food and social representations overlap, one would expect the magnitudes of food and social responses across the population to be positively correlated. Indeed, the correlation between food and social responses across neurons was significant (Figure 2E for food versus male, R=0.29, p<0.001, Figure 2F for food versus estrus female, R=0.24, p<0.001). To rule out that the relationship between these responses was due to shared cues associated with the arrival of either stimulus (e.g. the sound of the motor), we controlled for the arrival of the empty cage by performing a partial correlation analysis in which, across neurons, the magnitudes of both empty cage and male responses were used to predict the food response. The partial correlation between responses to food arrival and social stimuli were still significant (male, Figure S3A, R=0.23, p<0.001; estrus female, Figure S3B, R=0.17, p<0.001), implying that indeed the overlap in food and social responses is due to encoding of features other than simply motor arrivals.

We investigated the possibility that shared salience or arousal responses could explain the overlapping food and social responses (Figure S4-5). There was not significant overlap in neurons responsive to a mild but salient (flinch-inducing) air puff with food or social stimuli (Figure S4A-F). Furthermore, regressing out each neuron’s response to the salient air puff could not explain the significant relationship between the magnitude of food and social responses (Figure S4G-H). Regressing out the change in pupil diameter upon stimulus presentation also did not explain the relationship between food and social responses (Figure S5A-B).

Significant overlap in food- and social- responsive populations was observed regardless of whether the first stimulus was food or social, and neurons responsive to the final food presentation were significantly overlapping with those responsive to the first male presentation (Figure S6). We did not observe a significant relationship between the anatomical location of recorded neurons and the magnitude of responses to either stimulus (Figure S7).

### Hunger increases the fraction of VTA^DA^ neurons responsive to both food and social stimuli

If food and social representations overlap in VTA^DA^ neurons, one might predict that a change in internal state that affects food motivation would change social responses as well as food responses. An alternative possibility, which would be expected if food and social representations are separate within VTA^DA^ neurons, is that hunger increases only food (and not social) responses in the VTA.

Our experimental paradigm enabled longitudinal recordings from the same population while changing internal state (e.g. food motivation) across days. To change motivation for food, we induced a “sated” state with pre-feeding of sweetened condensed milk and a “hungry” state with 15-20 hours of fasting prior to imaging (Figure 3A). An example field of view tracked across conditions is shown in Figure 3B. To control for time-dependent effects, the hungry and sated sessions were counterbalanced and all mice received two imaging sessions in each state (Figure S8A; Figure 3A-F shows data from the first of the two sated/hungry session pairs).

**Figure 3.**
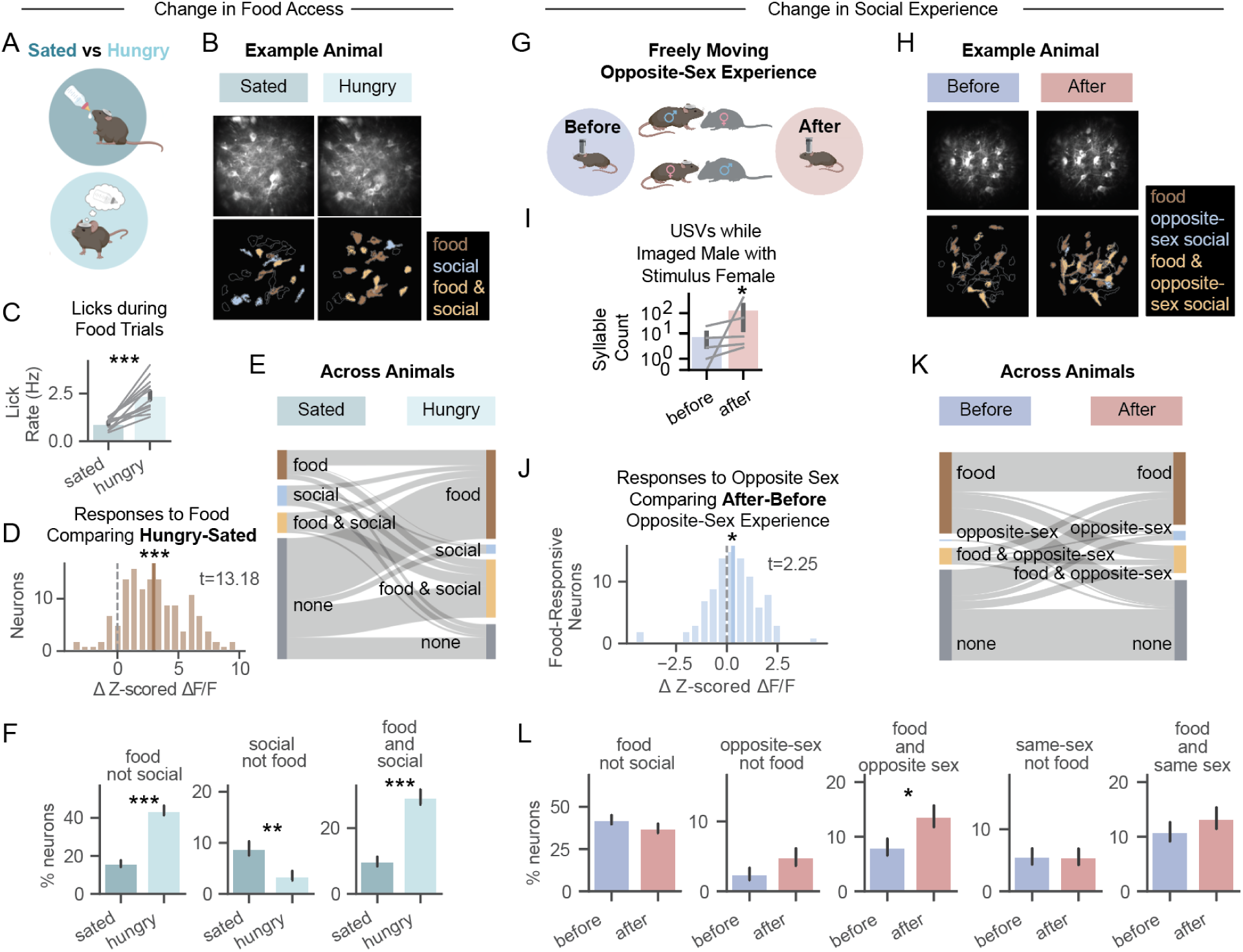
Hunger and opposite-sex experience increase fraction of VTA^DA^ neurons responsive to both food and social stimuli. (A) Cartoon representing comparison of imaging under sated (hand-fed sweetened condensed milk prior to experiment) or hungry (12-18 hr fasted) conditions (same animals, separated by 48 hrs, counterbalanced order of sated versus hungry day). (B) Example field of view across imaging sessions (sated: N=22 neurons, hungry: N=20 neurons). (C) Lick rates during food presentation (average across 20s trials, N=13 mice). (D) Distribution of change in activity in response to food from sated to hungry sessions (N=137 neurons). (E) Changes in neural tuning categories from identified cells tracked between sated and hungry sessions. (N=217 neurons) (F) Percentage of neurons responsive to food arrival, social (male and estrus female) arrival, or both on sated vs hungry sessions (error bars show standard deviation of binomial distribution, sated: N=427 neurons, hungry: N=394 neurons). (G) Cartoon representing comparison of imaging before and after freely moving opposite sex experience. (H) Example field of view across imaging sessions (before: N=34 neurons, after: N=35 neurons). (I) USV syllable counts while males are presented with female stimuli after versus before opposite sex experience (y-axis log scale, 5 out of 5 males increased in USVs detected during female trials). (J) In neurons that are food-responsive (before or after opposite sex experience), distribution of change in activity in response to opposite-sex arrival (N=95 neurons). (K) Changes in neural tuning (responsive to food arrival, opposite-sex arrival, both, or neither) before versus after opposite sex interaction (N=227 neurons). (L) Comparison of percentage of neurons responsive to food arrival, opposite-sex arrival, food and opposite-sex arrival, same-sex arrival, and food and same-sex arrival both before versus after freely moving opposite-sex interactions (error bars show stdev of binomial distribution, before: N=321 neurons, after: N=306 neurons). **See Table S17 for statistics.** ∗p < 0.05, ∗∗p < 0.01, ∗∗∗p < 0.001. Unless specified, data plotted as mean ± SEM.

Validating the manipulation, hunger induced behavioral changes consistent with increased motivation to acquire food (while there was licking in both states, there was more licking during hunger; Figure 3C).

Hunger broadly reorganized food responses across the population (Figure 3D-F). The magnitude of individual neurons’ responses to food increased on the hungry compared to the sated day (Figure 3D, N=137 neurons, t-test, T=13.18, p=4.5E-26), as did the proportion of neurons responsive to food and not social stimuli (Figure 3F; N=68/427 vs 173/394 neurons, Z=9.10, p=8.89E-20).

Accounting for lick rate, running speed, and pupil diameter did not change these conclusions. When including the lick rates along with sated versus hungry state as regressors of neural activity, state had significant explanatory power while lick rate did not (generalized estimating equation (GEE) regression by state: Z=2.21, p=0.027; lick rate Z=1.82, p=0.068). Running speed was not significantly different during the time window we analyzed neural data between sated and hungry states (Figure S5C,E,G). Pupil diameter also did not significantly differ between sated and hungry states change upon food arrival (Figure S5D,F). Thus, behavior or arousal differences across states are unlikely to fully explain neural response changes between sated and hungry conditions.

Interestingly, hunger also reorganized social responses across the VTA^DA^ population (Figure 3E-F). While the proportion of neurons responsive to social stimuli (and not food) decreased in a hungry state (Figure 3F; N=38/427 vs 14/394 neurons, Z=3.14, p=0.0017), the proportion of neurons responsive to both food and social stimuli increased (Figure 3F; N=42/427 vs 116/394 neurons, Z=7.20, p=6.18E-13). This increase in neurons responsive to both stimulus types implies overlapping representations of food and social stimuli, rather than a separate food-responsive subpopulation that is selectively modulated by food motivation.

When repeating the same experiment across days without changing food motivation, individual neurons had very similar magnitudes of responses to food (Figure S8B-C). Thus, the strength of responses in individual neurons was stable over time but depended on food motivation.

Moreover, the extent to which individual neurons increased food-responsiveness with hunger was highly consistent across 2 weeks of imaging, implying that VTA^DA^ neurons are also stable in their sensitivity to state changes (Figure S8D). Interestingly, the extent to which neurons increased their food responses with hunger was related to the anatomical depth of the fields imaged within the VTA (Figure S9, N=12 fields, R=-0.745, p=0.016).

### Opposite-sex experience increases the fraction of VTA^DA^ neurons responsive to both food and social stimuli

Social experience is known to induce profound changes to behavior and neural circuits, including in the midbrain DA system (Panksepp and Beatty 1980; C. R. Lee, Chen, and Tye 2021; Matthews et al. 2016; Tomova et al. 2020; Mumtaz et al. 2018; Tenk et al. 2009; Whitten 1956; Remedios et al. 2017; S. X. Zhang et al. 2021; McHenry et al. 2017; Willmore et al. 2022; Dai et al. 2022). If social and food representations overlap in VTA**^D^**^A^ neurons, one would predict that a change in social experience that affects social motivation might reorganize responses across both social- and food-responsive VTA**^D^**^A^ neurons.

We assessed this possibility by imaging animals several days before and after they received two hours of free social interaction with the opposite sex during which mounting was observed (Figure 3G). An example field of view before and after social experience is shown in Figure 3H. Mice in these experiments were at baseline with respect to food motivation (neither fasted nor pre-fed). Consistent with increasing social motivation, opposite sex interaction increased the number of ultrasonic vocalizations (USVs) detected while imaged males were presented with females (Figure 3I, Figure S10A-C). USVs are known to be emitted by freely moving males in response to female cues (Whitney et al. 1974; Chabout et al. 2015). Females are also known to vocalize while being courted by males (Neunuebel et al. 2015); however while presenting females with male stimuli, USVs did not consistently increase after opposite-sex experience (Figure S10D).

Opposite-sex experience reorganized VTA^DA^ responses to the opposite sex (Figure 3J-L). Specifically, opposite-sex experience increased the magnitude of opposite-sex (but not same-sex) responses when considering all imaged neurons (Figure S11; S12), as well as when considering only the food-responsive population (Figure 3J-L). As further evidence of opposite-sex experience reorganizing social responses, we could better decode whether social contact was with the same sex or opposite sex stimulus animal based on neural ensemble activity in response to bouts of social contact after opposite-sex experience (Figure S2D).

Social experience also reorganized responses to food, supporting the idea of social motivation altering VTA^DA^ population responses broadly across ingestive as well as social inputs. Specifically, after opposite-sex experience, the fraction of neurons responsive to both food and opposite-sex increased (Figure 3L; N=26/321 vs 42/306 neurons, Z=2.24, p=0.0253), while the proportion of food not social (or opposite-sex not food) neurons remained consistent (Figure 3L; food-only: N=136/321 vs 114/306 neurons, Z=-1.30, p= 0.192; opposite-sex only: N=8/321 vs 15/306 neurons, Z=1.54, p=0.12). The majority of neurons responsive to both food and opposite-sex after opposite-sex experience were responsive to only food before this experience (Figure 3K). This increase in neurons responsive to opposite-sex and food was observed in both males and females (Figure S11B). In contrast, the percent of neurons responsive to both *same-sex* conspecifics and food did not change with opposite-sex experience (Figure 3L, N=35/321 vs 17/306 neurons, Z=0.949, p=0.343; Figure S12A). This flexibility in representation of food and social stimuli provides further evidence that socially-responsive VTA^DA^ neurons are not distinct from food-responsive neurons, but rather overlapping populations. Control behavioral measurements such as average distance between imaged and stimulus mice (Figure S2E), imaged mouse pupil dilation (Figure S5H), and imaged mouse running speed (Figure S5I) did not differ before versus after opposite sex experience and thus were unlikely to explain these changes in neural activity with social experience.

The change in magnitude of responses to opposite-sex stimuli with hunger or experience depended on the anterior-posterior and medial-lateral locations of the neurons. S(Figure S13, N=137 neurons; for AP, R=-0.385, p=1.03E-5 and for ML, R=-0.214, p=0.036). Greater changes in social responses were found in neurons located in more posterior and medial portions of the VTA.

To control for time-dependence and confirm that changes in neural representations after opposite-sex experience are a result of the freely moving social interactions, we considered control data with extended habituation but no opposite-sex experience. In this case, the extent of overlap between food and social responses (opposite or same sex) was not changed by extended habituation to imaging, and in fact opposite-sex responses decreased with habituation (Figure S14B, Z=-4.14, p=3.48E-5).

### Changes in expression of excitability-related genes in VTA^DA^ nuclei across sated and hungry states

Our imaging data suggested greater overlap in responses to food and social stimuli in VTA^DA^ neurons in a higher motivational state (e.g. hunger). We wondered whether there might be changes in gene expression in these neurons across states that could help explain these functional changes. For example, altered expression of genes associated with neuronal excitability could potentially increase responsivity to multiple stimuli (food and social) and thus produce more overlapping representations. We thus performed droplet-based single-nucleus RNA sequencing (snRNA-seq) of VTA tissue from hungry and sated mice to examine gene expression at the level of individual neurons (Figure 4A, Figure S15; (Macosko et al. 2015). This approach allowed us to investigate both DA neurons as well as other neurotransmitter-defined neuronal subtypes (GABA and glutamate) within the VTA.

**Figure 4.**
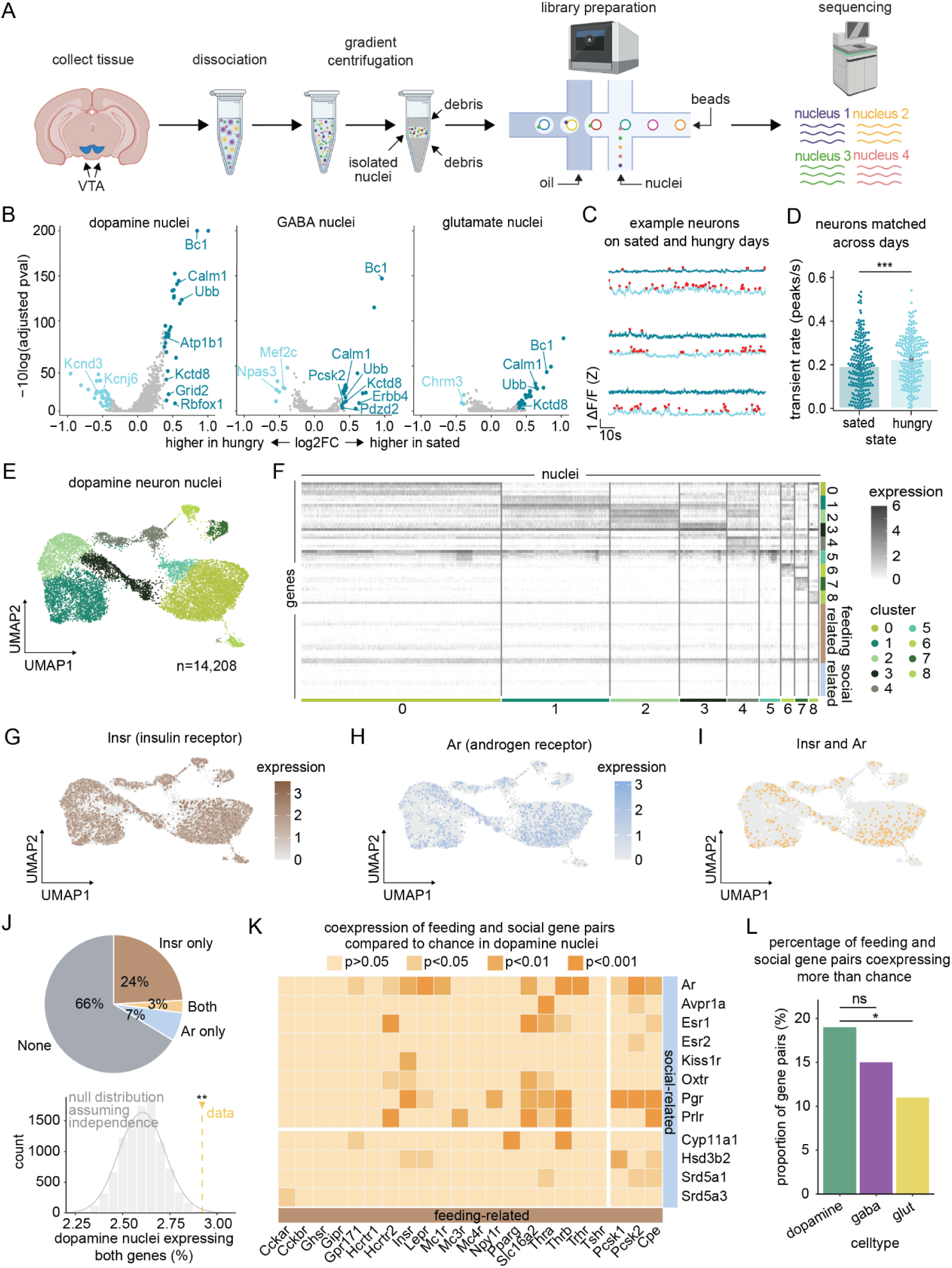
snRNA-seq reveals changes in excitability-related genes in VTA^DA^ neurons with hunger, as well as widespread and overlapping expression of feeding- and social-behavior related genes. (A) snRNA-seq pipeline includes tissue collection, nuclei isolation (dissociation and gradient centrifugation), library preparation, and sequencing. (B) Volcano plots showing genes significantly differentially expressed in nuclei from sated versus hungry animals across neuronal subtypes (dopamine, left; GABA, middle; glutamate, right; see Table S24 for genes and statistics). (C) Representative calcium traces from three neurons recorded on sated and hungry days (red dots indicate transients). (D) Average transient rate in neurons recorded on sated and hungry days (N=218 neurons). (E) Uniform approximation and projection (UMAP) of *Slc6a3*- or *Th*-expressing (dopamine nuclei) neuron subclusters in mouse VTA. (N=14,208 nuclei). (F) Heatmap showing expression of the top marker genes for each dopamine neuron subcluster as well as feeding and social hormone-related genes across subclusters. (G) UMAP of dopamine nuclei colored by expression level of *Insr* (see Table S25 for description of gene). (H) UMAP of dopamine nuclei colored by expression of *Ar* (see Table S25 for description of gene). (I) UMAP of dopamine nuclei, where nuclei are colored whether they express both *Insr* and *Ar*. (J) Percentage of dopamine nuclei expressing *Insr*, *Ar*, both, or neither (top). True percentage of dopamine nuclei co-expressing both *Insr* and *Ar* (yellow line) compared to a null distribution assuming the nuclei expressing each are independent samples (bottom). (K) Matrix showing the level of co-expression (compared to chance, assuming the nuclei expressing each gene are independent samples) of each food/social gene pair in dopamine nuclei. Horizontal break separates social-related hormone receptors (top) and enzymes (bottom). Vertical break separates feeding-related hormone receptors (left) and enzymes (right). (L) Percentage of gene pairs overlapping more than chance (based on comparison to null distribution constructed as described above) across neuron subtypes. **See Table S24-S28 for statistics and gene lists.** ∗p < 0.05, ∗∗p < 0.01, ∗∗∗p < 0.001 unless otherwise specified.

Hunger induced more changes in gene expression in VTA^DA^ neurons as a population than other neuronal subtypes in the VTA (Fig. 4B; 62 differentially-expressed genes for DA, 27 for GABA, 35 for glutamate; see Table S24). Interestingly, several genes implicated in excitability, excitatory/inhibitory balance, and sensitivity to synaptic inputs were altered in hunger. These include a number of potassium channels (*Kcnd3*, *Kcnj6* and *Kctd8*) as well as *Bc1*, *Npas3*, and *Mef2c*. Consistent with broad changes in excitability or input strength across hunger states, neurons in our imaging dataset had significantly higher transient rates on hungry versus sated imaging days (Figure 4C-D, t=5.6, p=7E-8).

### Overlapping expression of feeding- and social-hormone-related genes in VTA^DA^ neurons

Given the overlap in food and social responses in individual VTA^DA^ neurons (Fig 2,3), we wondered if there was also overlap in expression of feeding and social hormone-related gene expression in these neurons. Hunger and social experience trigger the production of circulating hormones (Coll et al. 2008; Hommel et al. 2006; Adkins-Regan 2009; O’Connell and Hofmann 2011; Song et al. 2016; Whitten 1956). Receptors for (and conversion enzymes associated with) many of these hormones are expressed in the VTA (Simerly et al. 1990; Mitra et al. 2003; Hung et al. 2017; Guan et al. 1997; Jerlhag et al. 2007; Hommel et al. 2006; Fulton et al. 2006; Song et al. 2016; Xiao et al. 2017; Lippert, Ellacott, and Cone 2014; Zigman et al. 2006). While the binding of these hormone receptors would not be expected to directly drive food or social responses, this binding endows neurons with state-dependent modulation of responses through changes in excitability or synaptic inputs (McHenry et al. 2017; Dey et al. 2015; Roepke et al. 2007; Marlin et al. 2015; Hung et al. 2017; Xiao et al. 2017; Hommel et al. 2006; Kendrick and Drewett 1979; Abizaid et al. 2006; van der Plasse et al. 2015; Labouèbe et al. 2013; Thompson and Borgland 2013).

Here we focused on genes that are more strongly associated with either feeding or social behavior (see Table S25 for gene list and references), including canonical receptors such as leptin receptor (*Lepr*) and oxytocin receptor (*Oxtr*) as well as enzymes involved in the conversion of hormones into active constituents such as proprotein convertases 1 and 2 (*Pcsk1/2*).

To characterize patterns of social and feeding hormone-related gene expression across VTA^DA^ subpopulations, we subclustered the DA nuclei (Figure 4E) and assessed which genes differed most in expression across clusters. In contrast to genes identified as cluster-specific markers (Table S26), expression of feeding and social hormone-related genes was dispersed across clusters, indicating that these genes do not define distinct DA neuron clusters (Figure 4F). For example, *Insr* (insulin receptor, a feeding-related gene) and *Ar* (androgen receptor, a social-related gene) had widespread expression (Figure 4G-I).

We next considered overlap of all pairs of feeding and social hormone-related genes and found significant co-expression in VTA^DA^ nuclei. For example, *Insr* and *Ar* are co-expressed more than expected assuming independent expression of each gene (Figure 4J, p<0.01 compared to a null distribution constructed based on the fraction of nuclei expressing each gene alone and assuming independence, see Methods). This significant level of co-expression was representative of many of the gene pairs (Figure 4K). There was more co-expression in VTA^DA^ compared to VTA^GLUT^ nuclei, and also a trend of greater co-expression when compared to VTA^GABA^ (Figure 4L, p<0.01; Figure S16).

Thus, we observed overlap in neural representations of food and social stimuli in VTA^DA^, as well as widespread and overlapping expression of food and social hormone-related genes in these neurons.

## Discussion

We aimed to determine whether the same or different populations of VTA^DA^ neurons are sensitive to food and social stimuli. We found that (1) the fraction of VTA^DA^ neurons functionally responsive to both food and social stimuli is significantly greater than chance (Figure 2), (2) the fraction of VTA^DA^ neurons responsive to both food and social stimuli increases with hunger and after social experience (Figure 3), and (3) co-expression of genes related to feeding and social behavior hormones in single VTA^DA^ neurons is significantly greater than chance (Figure 4). Each of these findings provides complementary evidence that overlapping VTA^DA^ populations are modulated by food and social stimuli and could underlie both types of motivation.

### Single DA neuron responses to social stimuli in a head-fixed prep

Our head-fixed social stimulus delivery paradigm offered a level of control not found in freely moving contexts, while maintaining the ability to elicit social behavior such as vocalizations (Weiner et al. 2016). In our assay, interactions were all passive (not self-initiated) and primarily reciprocal (nose-to-nose), allowing us to consistently probe the same types of interactions across days of imaging.

Despite head-fixation, many features of the responses to conspecific presentation mirrored previous published results in the freely moving context. For example, we found that roughly half as many VTA^DA^ neurons are responsive to social stimuli as food stimuli (21% of neurons respond to male arrival, while 46% respond to food arrival, Figure 2C), consistent with prior fiber photometry in freely moving mice, which found the magnitude of social responses about half that of food (Gunaydin et al. 2014). Moreover, our observation that about 23% of VTA^DA^ neurons responded to passive, reciprocal interactions is also consistent with recent findings (Solié et al. 2021) in which different types of freely moving interactions recruited between 7% to 53% of putative DA neurons. Finally, we noted rapidly attenuating responses to social stimuli across trials, which is also consistent with multiple previous reports in the freely moving setting (Gunaydin et al. 2014; Solié et al. 2021; Dai et al. 2022; Robinson, Heien, and Wightman 2002; Damsma et al. 1992).

### Implications of overlapping food and social representations in VTA^DA^ neurons

A successful decision involves weighting different types of outcomes into a so-called “common currency”. Our results suggest that in the case of unconditioned stimuli such as social stimuli and palatable food, responses overlap in single neurons in VTA^DA^ more than expected by chance. Computationally, such a signal could potentially function to produce downstream value (or decision-related) representations that combine food and social stimuli into a common currency.

In contrast to the overlap in responses to multiple unconditioned stimuli observed here, we and others have observed heterogenous coding of conditioned stimuli and task variables during complex behaviors (Kremer et al. 2020; Engelhard et al. 2019). The contrast between heterogeneous coding of conditioned stimuli and overlapping representations of unconditioned stimuli may be an organizing principle of the midbrain dopamine system (R. S. Lee et al. 2022).

An interesting question is if the shared response to food and social stimuli might be due to a shared salience, arousal, or novelty feature, rather than a shared reward-related feature (Lutas et al. 2019; Cai et al. 2020). While salience does drive some responses in our setup (positive responses to a salient airpuff, Figure S4B), these airpuff responses did not significantly overlap with food or social responses, and regressing out these responses could not explain the correlation between neurons’ responses to food and social stimuli (Figure S4C-H). Similarly, pupil dilation and running speed (readouts of arousal) do not appear to drive the relationship between food and social responses (Figure S5). On the other hand, the fast adaptation of the responses to social stimuli, both here and in previous work (Gunaydin et al. 2014; Solié et al. 2021; Dai et al. 2022; Robinson, Heien, and Wightman 2002; Damsma et al. 1992), argues that social responses are indeed novelty-related. However, novelty cannot entirely explain the responses to social stimuli or overlap with food. This is because we find that opposite sex experience enhances opposite sex (but not same sex) response and overlap with food, while novelty alone would instead predict attenuated response to a stimulus after exposure. Moreover, we do not think that a “shared novelty feature” between food and social stimuli explains the overlapping representations, given that the food was familiar and the response to the final food presentation of the day also significantly overlaps with the novel social stimulus response (Figure S6C-D). Thus, while salience, arousal, and novelty are encoded by dopamine neurons, these features are unlikely to completely account for the overlap in food and social responses.

### Changes in internal state and experience broadly alter DA responses

When an animal’s motivation changes, the magnitude of the VTA^DA^ responses for that motivator would be expected to change (Mazzone et al. 2020; Branch et al. 2013). However, it is less clear how responses to other motivators may change. Interestingly, we observed that hunger and social experience reorganized VTA^DA^ responses broadly, beyond neurons tuned exclusively to either stimulus type (Figure 3). Specifically, we found that both manipulations further increased the proportion of neurons that respond to *both* food and social stimuli. This demonstration that changing motivation for one stimulus reorganized the population response to the other stimulus was unexpected, and implies the food and social responding populations are fundamentally overlapping. These results may parallel prior observations that hunger not only increases behavioral sensitivity and dopamine release to food, but also to drugs such as cocaine (Shen et al. 2016; Zheng, Cabeza de Vaca, and Carr 2012). These findings are also potentially consistent with our observation that the high motivational state of hunger is associated with differential expression of genes known to be related to neuronal excitability and induced increased calcium transient rates in VTA^DA^ neurons (Figure 4), as a general change in excitability could produce stronger responses to weaker inputs and therefore could generate more overlap in food and social responses. Likewise, it is possible that opposite sex experience also alters expression of genes related to excitability, which can be examined in future studies.

Changes in an animal’s internal state have been shown to alter neural activity in the VTA via fluctuations in hormone receptor expression and binding (Marlin et al. 2015; Riediger et al. 2003; Shahrokh et al. 2010; Hung et al. 2017; Dölen et al. 2013; Song et al. 2016; Groppe et al. 2013; Xiao et al. 2017; Georgescu et al. 2005; Hommel et al. 2006; van den Heuvel et al. 2015; McHenry et al. 2017). Our finding that the same DA neurons tended to co-express both food and social hormone-related genes–and at levels higher than other neuronal subtypes (Figure 4)–implies that the same neurons may be sensitive to the hormonal changes that occur in both types of state changes. This is broadly consistent with our observation from the functional imaging of overlapping representations of food and social stimuli. Future work might directly investigate the causal relationships between internal state changes, hormone levels, other molecular changes that may vary with internal state (such as receptor availability in VTA^DA^ neurons), and the response profiles of these neurons.

## Acknowledgements

We would like to thank Meaghan Creed, Annegret Falkner, Fenna Krienen, Chris Zimmerman, Junuk Lee, and Lindsey Brown for providing feedback on this work, as well as the Witten and the Pena labs for their support. This research was funded by NIH T32MH065214 (L.W.), NSF GRFP DGE-2039656 (A.M. & L.W.), ARO W911NF1710554 (I.B.W.), NIH R01 DA047869 (I.B.W.), NYSCF (I.B.W.), SCGB (I.B.W.). PNI Research Innovator Award (C.J.P, I.B.W.), NIH R01MH129643 (C.J.P.).

## Contributions

L.W., B.E, M.M., A.R.M. and I.B.W. conceived of the project and designed the experiments; L.W., B.E., M.M., A.R.M., and B.M. collected the data with support from S.Y.T.; L.W., A.R.M., and N.O. analyzed the data with support from B.E.; L.W., B.E., A.R.M., C.J.P., and I.B.W. wrote the manuscript.

## Methods

### BEHAVIOR AND NEURAL RECORDING

#### Mice

Animal procedures were conducted in accordance with standards from the National Institutes of Health and under the approval of the Princeton University Institutional Animal Care and Use Committee.

Experimental animals for imaging were comprised of 20 mice (11 males and 9 females) cross bred between DAT::cre (Jackson Laboratory strain 006660; (Lammel et al. 2015)) and the Ai148 GCaMP6f reporter line (Jackson Laboratory strain 030328; (Daigle et al. 2018)), a cross extensively characterized previously (Engelhard et al. 2019). The above mice were housed with littermates until 2 weeks prior to imaging, at which time they were singly housed with enrichment materials (running wheel, nestlets). Stimulus mice and mice used for snRNA-sequencing were C57 / BL6 WT (Jackson Laboratory) males and females between the ages of 10 and 16 weeks. Stimulus mice were non-littermates of imaged animals. Stimulus animals were housed with their cage mates throughout the experiment. Food and water were given *ad libitum*, except during 2 experimental food deprivation days for both imaging mice and hunger state manipulated sequencing mice. Prior to and throughout experimental assays, experimental and stimulus animals were housed under a 12 H light-dark cycle with experiments exclusively taking place during the dark phase.

#### Surgery

Between the ages of 8 and 20 weeks, experimental animals were deeply anesthetized (3-5% isoflurane for induction and 1-3% for maintenance) and leveled in a stereotaxic frame. Hair from the scalp was removed, the scalp sterilized, incisions were made to expose the skull with the edges of the skin sealed to the skull with a thin layer of Vetbond (3M), and the periosteum was removed using a bone scraper. To assist with motion correction, we injected 800 nl of a viral AAV9.CB7.CI.mCherry.WPRE.rBG (University of Pennsylvania Vector Core) at a titer of 1.84e+12 parts per mL anterior to the ventral tegmental area (VTA, AP -2.6, ML +/- 0.5, DV -4.7 mm from the skull surface at bregma, unilaterally, randomly counterbalanced across animals). Following viral injections, a craniotomy and durotomy were carefully performed to create a clean brain surface before implanting a .5 NA, 0.6mm diameter, ∼7.3mm long GRIN lens (GLP-0673, Inscopix) over the VTA (AP -3.2, ML +/- 0.5, DV -4.2 mm, on the same side as the viral injection, zero’d at the skull surface at bregma with lens paper between the skull and lens). The lens was held by a custom 3D-printed holder and implanted by slowly lowering it 0.8mm and then raising it 0.4mm until the desired depth was reached. Then, the lens was affixed to the skull with a small amount of metabond (Parkell) injected into the space between the lens and skull using a 1- mL syringe and 18-gauge needle. After 20 minutes, the lens holder was loosed and carefully raised before additional metabond was applied between the lens and skull. Finally, a titanium headplate was affixed to the skull, and a titanium ring (to contain water for imaging through the water-immersed objective) was affixed to the headplate with the lens in its center. Animals were allowed to recover for at least 6 weeks between subsequent experimentation.

#### Stimulus delivery paradigm

Experimental animals were head-fixed while supported by an air-suspended spherical treadmill. Stimuli were delivered to head-fixed animals on a 2-foot slider powered via a stepper motor (G5-GVM-GT60, GMV). Slider movement was triggered by custom behavior-control MATLAB code on a dedicated computer, routed to the motor through a NI-DAQ board (National Instruments, NI CB-68LP), Master-9 Pulse Stimulator (AMPI) for generating precise square waves, and stepper motor driver (DM542T, OMC Corporation Ltd). Stimulus movement was controlled at a rate of 8 inches/second. Stimuli were held to the treadmill with magnets and exchanged during the inter-trial interval. Stimulus types included a food stimulus (20% sweetened condensed milk diluted in water), 3 social stimuli (male C57, female C57 in estrus, and female C57 not in estrus), and a control empty stimulus. To determine whether a female stimulus mouse was in estrus, vaginal smears were evaluated as previously described (Byers et al. 2012) at the start of each imaging day. Food stimuli were presented on a custom 3D-printed spoon and social stimuli were presented in custom 3D-printed cages (3.5’ x 1.4’ x 1.7’), all made with fine-detail, smooth plastic (by Shapeways). The social stimulus cages included transparent acrylic bars at the front (to allow for social interaction) and a transparent acrylic side door (to allow for video acquisition of stimulus mouse behavior.

In an imaging session, each stimulus type (5 types total) was presented 10 times (50 trials total) in a randomly interleaved order. Stimulus presentation lasted for 20 seconds per trial, and intertrial-intervals were random in duration, 30s plus an additional interval following an exponential distribution with a mean of 20s. Stimulus loading occurred at the end of the slider away from the imaged mouse, and stimulus movement toward and away from the mouse took 3s. Stimulus animals were novel to the imaging mice at the beginning of each session, with the same stimulus animal used across trials in a single session, and were free to move within the stimulus presentation cage. Records of the stimulus order were collected after each session.

In addition to the main stimuli (5 types above), three salient yet mild air puffs were delivered at the end of the main stimulus delivery session. Immediately following the last main stimulus delivery and while the imaged mouse and imaging setup remained in the same position, a narrow air spout was aimed at the right whisker pad of the imaged mouse. At random intervals (at least 30s plus an additionally exponentially-distributed period, mean=20s), an air puff of 18 PSI and 200 ms in duration was released via a solenoid under software control. Neural imaging was continuous throughout the main stimulus delivery, the air puff setup, and the air puff delivery.

#### Experimental timeline

Following surgery recovery of at least 6 weeks, animals were briefly assessed for quality of imaging in a 10-20 minute 2-photon calcium imaging session. Spontaneous activity without delivering stimuli was monitored, and only the animals with the clearest fields of view were carried forward with the experiment (20 animals across 4 cohorts, 44% of all mice that underwent surgery).

Animals selected for undergoing experiments were singly housed 2 weeks before undergoing imaging experiments. One week before imaging, mice were habituated to handling and sampling sweetened condensed milk in their home cages. At this time, stimulus animals were habituated to dwelling within stimulus-presentation cages. In a final habituation step, mice were imaged and exposed to the final stimulus delivery paradigm without seeing social stimuli. That is, while head-fixed, mice were imaged and presented with randomly interleaved empty control and food stimuli for approximately half an hour. During this time, the spoon location was manually adjusted until mice learned to lick the food reward. Data from this final habituation session was not included in our study.

Subsequent imaging sessions, for which we present data in Figures 1-3, included the use of the same stimulus delivery apparatus, same spoon, same food, and same cages as used in the habituation sessions. Social stimuli were novel to the imaged mouse on each day of imaging.

Data presented in Figures 1-3 captures the neural responses to the first trial in which an imaged mouse encounters each novel conspecific.

For the following 2-3 weeks, mice each underwent 3 imaging sessions in which mice experienced reward delivery as described above. On 2 of the imaging weeks, animals were imaged under conditions of varying food motivation: 1) baseline, with no manipulation; 2) hunger, in which food was removed and bedding cleaned from animals’ home cages 15-20 hours before imaging the following day, at which point ad libitum food was resumed; and 3) sated, in which animals were free to consume 20% sweetened condensed milk (SCM) for 1 hour prior to imaging and received additional, hand-fed 20% SCM after head fixation but immediately prior to beginning imaging and stimulus delivery. These food motivation state manipulations were repeated twice for most mice in a counterbalanced order. All animals in the same cohort received the same motivation change and imaging schedule. In order to alter the animals’ social experience and motivation, between weeks 1 and 2 imaged mice were given 2 hours to freely interact with a C57 mouse of the opposite sex. Interactions were monitored via video and manual annotation. Mounting was observed across all pairings but there was no evidence of vaginal plugs, suggesting these interactions did not lead to intromission. This interaction took place 1-2 days before the subsequent imaging session. See Figure S3A for a timeline of food access and social experience manipulations.

The timeline of data collection and internal state or experience changes can be found in Figure S8A.

#### Two-photon calcium imaging

Calcium imaging was performed using two custom-built two-photon microscopes as previously described (Engelhard et al. 2019). The microscopes were equipped with either a pulsed Ti:sapphire laser (Chameleon Vision, Coherent) tuned to 920 nm or an Alcor 920 laser. The laser power reaching the sample was controlled by an Electro-Optic Modulator (350-80-LA-02 KD*P, Conoptics). The scanning unit used a 5-mm galvanometer and an 8-kHz resonant scanning mirror (Cambridge Technologies). The collected photons were split into two channels by a dichroic mirror (FF562-Di03, Semrock). The light for the respective green and red channels was filtered using bandpass filters (FF01-520/60 and FF01-607/70, Semrock), and then detected using GaAsP photomultiplier tubes (1077PA-40, Hamamatsu). The signals from the photomultiplier tubes were amplified using either a high-speed current amplifier (59-179, Edmund) or a transimpedance amplifier TIA60 (Thorlabs). Black rubber tubing was attached to the objective (Zeiss 420957-9900-000, 20x, 0.5NA) as a light shield covering the space from the objective to the titanium ring surrounding the GRIN lens. Double-distilled water was used as the immersion medium. The microscopes could be rotated along the mediolateral axis of the mice, allowing alignment of the optical axes and GRIN lens. Control of the microscopes and image acquisition was performed using the ScanImage software (Vidrio Technologies, (Pologruto, Sabatini, and Svoboda 2003)) that was run on a dedicated computer. Images were acquired at 30 Hz and 512 x 512 pixels. Average beam power measured at the front of the objective was 40–60 mW. From each animal, approximately 10-40 neurons were recorded.

#### Behavior data acquisition and processing

Videos of behavior were recorded using two BlackFly S cameras (FLIR); one focused on the imaged mouse right pupil and the other capturing the body movements of both the imaged mouse and social stimulus. For the body-focused camera, we used a 6mm fixed focal lens (TA1186AMPF, Tamron); for the pupil-focused camera, we used a close-focusing macro lens (ZOOM 7000, Navitar). For both cameras, a 304-785 nm bandpass glass filter (FGS550, ThorLabs) was used to remove flashing light from the 2-photon microscope. Simultaneous video acquisition and real-time compression from both cameras was performed via custom software Motif (LoopBio, on a computer running Ubuntu 18.04.5, equipped with an Intel Core i7-4790 CPU and Quadro P2000/PCle/SSE2 GPU). The body camera was mounted approximately 5.1 inches from the 2-photon objective. The pupil camera was mounted such that the tip of the lens was 140mm from the 2-photon objective. Images were recorded at 1024 x 1280 pixels and 100 frames per second. Animals were illuminated with infrared light.

Signals from separate systems were recorded via a Digidata 1440A interfacing to a Clampex 10.5 software (Molecular Devices) run on a dedicated computer. The following analogue readouts were recorded: (1) 2-photon galvanometer Y-scanning signal; (2) video frame capture signals mirroring the exposure opening on the behavior camera; (3) square-wave signal engaging the stepping-motor controlling stimulus treadmill movement.

To quantify licking (Figure 3C), frames in which the tongue was extended were detected. Specifically, an area surrounding the tongue was defined and the average difference in pixel intensity compared to when the tongue was not out was used to identify when the animal was licking (based on a manually selected threshold). In lick analysis, mice whose tongues could not be differentiated from the image background were not used. This resulted in omitting 5 mice from the first cohort in which the food delivery apparatus was the same color as the mouse tongues and 2 mice that drank the milk without licking.

To quantify pupil dilation, running speed, and social contact, videos were analyzed via DeepLabcut version 2.1. For pupil dilation, 8 points (making 4 diameters) across the pupil were tracked in images from the pupil-focused camera. 235 video frames from 19 animals and 19 video sessions and 4 cohorts were used in training, which was done under the default parameters. From the body camera running speed and social contact were tracked. For running speed, the longest toe tip of the left paw (nearest the camera) was tracked. For social contact, the center of eye of the simulus mouse and the nose time of the imaging mouse were tracked. All contextual points labeled but not used in the analyses were:

NoseTip, NoseTip, NoseTop, NoseTop, NoseBottom, NoseBottom, EyeTop, EyeTop, EyeBottom, Eye Bottom, EyeLeft, EyeLeft, EyeRight, EyeRight, Whisker1, Whisker1, Whisker2, Whisker2, Whisker3, Whisker3, Tongue, Tongue, LeftForepawTip, LeftForepawTip, LeftForepawHeel, LeftForepawHeel, R ightForepawTip, RightForepawTip, RightForepawHeel, RightForepawHeel, BoxTopFarCorner, BoxTopFarCorner, BoxTopNearCorner, BoxTopNearCorner, BoxBottomFarCorner, BoxBottomFarCorner, BoxBottomNearCorner, BoxBottomNearCorner, TargetEyeCenter, TargetEyeCenter, TargetNoseTi p, TargetNoseTip, TargetBottomJaw, TargetBottomJaw, TargetForepawTip, TargetForepawTip, TargetForepawHeel, TargetForepawHeel, TargetHindpawTip, TargetHindpawTip, TargetHindpawHeel, TargetHindpawHeel, TargetBackForepawTip, TargetBackForepawTip, TargetBackForepawHeel, TargetBackForepawHeel, TargetBackHindpawTip, TargetBackHindpawTip, TargetBackHindpawHeel, TargetBackHindpawHeel, TargetRear, TargetRear.

The input for training the body-tracking DLC network included 1371 video frames from 18 animals and 32 video sessions and 4 cohorts.

To quantify lick rates, because tongue tracking was not accurate in our hands, for every session and mouse a bounding box was drawn around a small region of interest in which dark background pixels were present when the mouse wasn’t licking and the tongue was out when the mouse was licking. The intensity of these pixels was measured across the first food trial and spikes in intensity were defined as licks.

#### USV acquisition and processing

To record vocalizations in the ultrasonic range, the AultraSoundGate 416H system (Avisoft Bioacoustics) was used via a set of two externally polarized condenser microphones. Audio data was recorded at 250kHz via Avisoft-RECORDER USGH software. Vocal syllables were detected using MUPET (Van Segbroeck et al. 2017). Cohort 1 and cohorts 2 and 3 were processed separately (due to different microphone placements). Syllable number was set to 60 with two rounds of refinement, to remove any broadcast noise or non-ultrasonic calls. For all other parameters, default values were used. For alignment to stimulus delivery, custom code was used to detect the sound of the slider movement, which occurred at the onset and offset of each stimulus delivery trial.

#### Calcium imaging pre-processing and fluorescence extraction

Motion correction and region of interest (ROI) extraction were performed using the Suite2P(Pachitariu et al., n.d.) software package for fast, large-scale two-photon imaging analysis, followed by manual curation. Unless expression of red fluorescence failed, motion correction was done on the red channel, otherwise the green channel was used. Default settings were utilized other than ’nimg_init’: 300, ’batch_size’: 500, ’spatial_hp’: 50.0, ’pre_smooth’: 2.0, spatial_taper’: 50.0, ’threshold_scaling’: 1.0, ‘sparse_mode’=True. Following one round motion correction, visual inspection of video quality was used to determine if videos should be run again. For low SNR sessions, suite2p was rerun with ‘two_step_registration’=True. For imaging sessions with large jumps, ‘maxregshift’ was increased.

ROIs detected by suite2p were pruned to only include neurons with obvious morphological features and GCaMP dynamics present in fluorescence traces.

Each neuron’s activity is presented as Z-scored ΔF/F. These traces were derived from the fluorescence in each neuron’s ROI, less the fluorescence of the surrounding neuropil (calculated via suite2p) scaled by a correction factor of 0.58. Neuropil subtracted signal was defined as F.

ΔF/F defined as (F-F_0_)/F_0_, where F_0_ is the mean of F. Z-scored ΔF/F was calculated by dividing ΔF/F by the standard deviation of F. Standard deviation was calculated across the entire recording session and separately for each session and neuron.

#### 1- Photon Imaging Analysis

##### Definition of Responsive Neurons

Neurons were defined as responsive to a stimulus type if their average Z-scored ΔF/F in the first second after the first time that stimulus was presented was over 1.5, which was slightly greater than 2 standard deviations (1.4) of the activity taken from randomly chosen windows across the intertrial interval across all neurons.

##### Test for Significant Overlap

To directly test whether food-responsive and socially responsive sets of neurons significantly overlapped, we compared the observed fraction of neurons both responsive to the first presentation of food and social stimuli to the fraction expected by chance. The chance overlap level was defined based on a null distribution constructed as follows. We randomly shuffled food-responsive and social-responsive labels across all imaged neurons, maintaining the overall number of neurons responsive to each stimulus. Then we noted what percentage of neurons with these random labels were assigned to be responsive to both food and social. We repeated this random shuffling 10,000 times, each time recording the percent of neurons with both food and social responses. The significance of the overlap (fraction of neurons with food and social responses) observed in our data was then defined as the likelihood that randomly simulated assignments produced as much or more overlap.

##### Matching Neurons Imaged Across Sessions

Neurons were tracked across sessions using a semi-automated pipeline with custom Python code. In a pairwise fashion, mean images of fields from 2 imaging sessions were shifted in x and y using phase cross correlation alignment (skimage.registration.phase_cross_correlation). These shifts were applied to images of the cell masks defined by suite2p. Cell masks that overlapped between the two days were labeled a match. The unique label for that match was then used if a cell from another session matched. Following this automated cell-matching pipeline, the matches were visualized and mistakes corrected.

##### Decoding Same vs Opposite Sex Contact

In Figure S2D, using neural ensemble activity from a single mouse in response to bouts of social contact, we trained a binary classifier on whether the bout of social contact was with a stimulus mouse of the same or opposite sex. Input to the classifier was a reduced representation of ensemble activity. For each bout, the original activity included the 1s trace (30 data points) of neural activity from all N neurons in the mouse’s field of view, generating an array of size 30N. All bouts M from the same or opposite sex were then stacked to create an input matrix of 30N x M. PCA was then used to reduce the 30N feature dimension to 10. Only sessions in which a minimum of 8 neurons were imaged and in which a minimum of 5 contact bouts with each sex occurred were used. The 10-dimensional vector was the input to a decision tree classifier (simple non-linear classifier, sklearn). For each mouse, 5-fold cross-validation was used; of the 5 splits, 70% of the data was used to train the classifier and 30% was used to test the accuracy. Accuracy was measured as the average F1 score across the 5 cross-validation splits. The procedure was done for mice on baseline (*ad libitum* food as opposed to fasted or sated) sessions before and after opposite sex experience. Results were presented in Figure S2D only for mice with valid analysis on both sessions (enough neurons and contact bouts).

#### Histology

Following imaging experiments, animals were deeply anesthetized with a lethal dose of ketamine/xylazine cocktail and perfused with 4% paraformaldehyde (PFA) dissolved in 1x phosphate buffer saline (PBS). Incisions were made between the skull and implanted headplate, avoiding cutting too deeply and damaging the GRIN lens to be removed, cleaned, and recovered for future use. After removing the headgear, brains were excised and soaked for post-fixation (also in 4% PFA in PBS) for 12-24 hours. Post-fixed brains were transferred to 30% sucrose for cryoprotection before being frozen in optimal cutting temperature mounting medium (Fisher Healthcare Tissue-Plus O.C.T. Compound, Fisher Scientific). Frozen brains were sliced at a thickness of 30 μM and directly mounted onto glass microscope slides. After washing slices 3 times with PBS+0.4% triton (PBST), slices were blocked for immunostaining by soaking for 30 minutes in blocking buffer (PBST + 2% normal donkey serum and 1% bovine serum albumin).

Slices were stained for green fluorescent protein (GFP) with a rabbit monoclonal anti-GFP antibody (G10362, Molecular Probes, 1:1000 dilution in blocking buffer) and for tyrosine hydroxylase (TH) with a chicken polyclonal anti-TH antibody (E.C. 1.14.16.2, Aves Labs, 1:200 dilution in blocking buffer) overnight at 4°C. After washing with PBST, slices were incubated with secondary antibodies (Alexa fluor 647 donkey-anti-chicken and Alexa fluor donkey-anti-rabbit, Jackson ImmunoResearch, 1:1000 dilution in blocking buffer). Once stained, slices were rinsed with PBS, briefly dried, and then coverslipped with mounting medium including a nuclear stain (EMS Immuno Mount DAPI and DABSCO, Electron Microscopy Sciences, Cat # 17989-98, Lot 180418). Coverslip edges were sealed with transparent nail polish. After 24 hours of drying, slides were digitally scanned at 20x magnification (NanoZoomer S60, C13210-01, Hamamatsu).

#### Anatomical registration

To investigate the relationship between the activity of the neurons and their location in the VTA, we estimated the location of each neuron by combining information about the position of the GRIN lens from histology with the location of the imaged neurons within the field of view.

To obtain the position of the GRIN lens in atlas coordinates, we registered the histological images using the WholeBrain software package (Fürth et al. 2018). In the software, we applied registration points using the VTA, substantia nigra pars compacta and cerebral peduncle as primary markers. After registration, we marked the center of the bottom of the lesion produced by the lens and extracted its atlas coordinates using the software (these are Allen CCF coordinates). We then combined the coordinates of the center of each lens with estimates of the distance of each neuron to the center of the lens to obtain the location of each neuron in atlas coordinates. The distances of the neurons to the lens centers were estimated from the in-vivo imaging as follows: first, we determined the depth of imaging (distance of imaged plane to the bottom of the lens) from z-stack movies taken in-vivo. We used this distance to obtain the depth-dependent magnification factor of imaging based on a calibration procedure described below and performed previously (Engelhard et al. 2019). This magnification factor was needed because GRIN lenses have different magnifications at different imaging depths. The magnification factor determined the microns-per-pixel scale factor of each imaging session. We calculated the distance in pixels from the center of the lens to the center of mass of each cell ROI and used the magnification factor to calculate the distance in microns of each cell to the center of the lens. To calculate the distance in pixels from the center of the lens to the center of mass of each neuron we estimated the location of the center of the lens in pixels from the circular shape of the lens in the in-vivo images, obtained the location in pixels of the center of mass of each neuronal ROI from the segmentation software (normcore) and then calculated the distance between the two. For those mice where z-stack movies were not available, we estimated the magnification factor using an interpolation procedure that compared the mean size of all the cell ROIs of the mouse to the mean size of cell ROIs and magnification factors of all the mice that did have z-stack movies. Finally, the absolute location of each neuron in atlas coordinates was determined as the vector sum of its estimated distance from the lens center in the field of view to the measured location of the lens center in atlas coordinates.

To calculate the depth-dependent magnification factor, we generated samples from a solution of agarose and fluorescent beads (10 μm, Molecular Probes). We first confirmed the size of the beads by imaging the samples directly with the two-photon microscope that was calibrated by previous imaging of a 10 × 10 μm grid (Thorlabs). We then proceeded to image the samples through the GRIN lens. We calibrated the magnification factor at each depth by measuring the observed size of the beads in the x–y axes, and used that size to estimate the magnification factor. To relate the movement of the stage in the z-axis with the imaging depth of the imaged fields, we also measured the observed size of the beads across the z-axis. The reference (0) plane was in all cases the first plane where we could form an image as the microscope’s objective approached the GRIN lens.

#### Statistical analysis of neural data

Tests for linear relationships were ordinary least squares regression (statsmodels.regression.linear_model.OLS) when repeated measures were not made (Figures 2E, S3, S4G, S4H, S5B, S2B, S8C, S9, S12B, S13) and GEE (generalized estimating equations, statsmodels.formula.api.GEE, groups=’mice’, family=’Gaussian’, cov_structure=’Independence’) when repeated measures were made (Figures 1F and S2C, and lick rate vs internal state analysis). Statsmodels is a Python-based package. We used version 0.13.2.

### SINGLE NUCLEUS RNA SEQUENCING

#### Tissue collection and single-nuclei dissociation

We collected VTA tissue from male and female mice in adulthood (P60). For hunger state manipulations, we manipulated access to food as described above – hungry: food was removed and bedding cleaned from animals’ home cages 15-20 hours before tissue collection the following day; sated: animals were hand-fed 20% sweetened condensed milk (SCM) 1 hour before tissue collection.

Animals from all conditions were cervically dislocated and brains were extracted and sectioned into 1mm coronal sections on ice using a brain block. Bilateral punches of VTA were taken, flash frozen in Eppendorf tubes, and kept at -80C until nuclei isolation and library preparation.

Nuclei were isolated into suspensions for snRNA-seq as previously described (Hrvatin et al. 2020), with minor modifications. Bilateral VTA punches from two animals of the same sex and condition were pooled together and Dounce homogenized. Pairs were selected randomly from within each sex and condition. Pooling ensured a high enough concentration of nuclei per sample for quality sequencing. The sample was filtered through a 40um strainer and nuclei were isolated via iodixanol gradient centrifugation. Nuclei were hand-counted using a hemocytometer on an EVOS M5000 microscope and final concentrations were 850-1,450 nuclei per ul. The volume of suspension used for library preparation was adjusted in order to load a target of 20,000 nuclei and capture 10,000 nuclei per sample.

#### snRNA-seq library prep and sequencing

Nuclei were captured and RNA was reverse transcribed and barcoded for library preparation using the 10X Genomics Chromium v3 platform. cDNA was amplified and adapters were added for sequencing on an Illumina NovaSeq SP 100nt Lane v1.5. Library preparation and sequencing was performed by the Princeton University Genomics Core.

#### snRNA-seq read mapping

The 10X Genomics package CellRanger (v6.1.1) was used to map transcripts to the mm10 mouse reference genome. In order to capture the high percentage of pre-spliced intronic mRNA present within the nucleus, the ‘--include-introns’ flag was used to map unspliced reads to corresponding genes. After quality control and initial filtering, we recovered 118,054 total nuclei expressing a total of 23,385 genes (median UMIs/nucleus=1,221; median genes/nucleus=858; Figure S15A-C). Neurons had greater numbers of UMIs/nucleus and gene/nucleus than non-neurons, which is comparable to previously reported snRNA-seq datasets of mouse brain (Hrvatin et al. 2020).

#### snRNA-seq Analysis

##### Preprocessing

CellRanger output count matrices were further analyzed using Seurat (v4.0.0) (Butler et al. 2018; Stuart et al. 2019) in R (v4.0.3). During microfluidic encapsulation of nuclei on 10X devices, some droplets may contain more than one nucleus. During reverse transcription, transcripts from co-encapsulated nuclei are labeled with the same cell barcode, creating a ‘doublet’. To remove likely doublets in each sample, nuclei with >2500 genes were removed. In addition, nuclei with <200 genes and >5% of reads mapping to mitochondrial genes were removed. Mitochondrial and ribosomal genes were then removed in order to remove these genes as a confounding source of variation in downstream analysis.

##### Dataset integration, UMAP embedding, and clustering

To integrate all datasets and cluster nuclei (Fig. S15), each sample dataset was log normalized and the top 2000 highly variable features were identified for downstream sample integration using the *FindVariableFeatures()* function. These features were used as input to the function *SelectIntegrationFeatures()*. We then scaled and ran PCA on each dataset separately. The 10 separate datasets–6 control (3 male and 3 female), 2 sated (1 male and 1 female), and 2 hungry (1 male and 1 female)–were then integrated using the *FindIntegrationAnchors()* and *IntegrateData()* functions. Next, we scaled and ran PCA on the integrated dataset. The *RunUMAP()* function was used to embed the nuclei into UMAP space and the *FindNeighbors()* and *FindClusters()* functions were used to cluster nuclei based on overall gene expression similarity using the top 10 PCs and a clustering resolution of 0.05. The majority of clusters represented established cell types in the VTA. However, one cluster contained nuclei that were dispersed across all other clusters and had a high percentage of reads mapped to mitochondrial genes, suggesting contamination. These nuclei were removed from downstream analysis.

##### Quality Control Metrics

Clustering confirmed the presence of expected neuronal and non-neuronal populations, which we labeled based on expression of established marker genes (Figure S15A-B). These populations included neuronal and non-neuronal cell types, such as astrocytes and oligodendrocytes, known to be present in the VTA (Figure S15A-B). Nuclei from male and female samples were distributed across all broad cell type clusters. (Figure S15D-S15E). The distributions of UMIs and genes were also comparable across the sexes (Figure S15F).

Similarly well distributed nuclei, UMIs, and genes were collected across samples from different hunger states (control, sated, hungry).

##### Identification of neuronal subtypes

Our analyses in Figure 4 focused on neuronal subclusters further defined by neurotransmitter type. Clusters were identified as neuronal or non-neuronal based on expression of *Syt1* (Figure S15A-B). Nuclei were assigned to neuronal subclusters as follows: DA nuclei had expression of *Slc6a3* (DAT) >0 or *Th* (tyrosine hydroxylase) (N=14,208); GABA nuclei had expression of *Slc32a1*, *Gad1*, or *Gad2* and *not Slc6a3*, *Th*, *Slc17a6*, *Slc17a7*, or *Slc17a8* (N=11,100); glutamate nuclei had expression of *Slc17a6*, *Slc17a7*, or *Slc17a8* and *not Slc6a3*, *Th*, *Slc32a1*, *Gad1*, or *Gad2* (N=4,435).

##### Determination of Differentially-Expressed Genes

Within VTA^DA^, VTA^GABA^, and VTA^GLUT^ neuronal subclusters, we examined differential gene expression in nuclei collected from hungry versus sated animals using DESeq2 ((Love, Huber, and Anders 2014)). We considered differentially expressed genes those with an adjusted value of 0.05 and an absolute value log2 fold change of greater than 0.38 (Figure 4B). Positive fold changes indicated higher expression in nuclei from sated animals whereas negative fold changes indicated higher expression in nuclei from hungry animals. See Table S24 for individual gene statistics.

##### Examination of Cluster-Specific Gene Expression

In Fig. 4E-I, we investigated whether there was systematically different expression of feeding versus social hormone-related genes across DA subclusters. First, nuclei assigned as DA neurons were scaled and clustered similarly to above, using “nfeatures” = 2000, “PCs” = 10, “n.neighbors” = 70, “min.dist” = 0.3, “spread” = 0.7, and “resolution” = 0.3. These parameters produced 9 DA subclusters (Figure 4E). We then identified the top 5 putative marker genes for each cluster using the Seurat *FindAllMarkers()* function (Table S26). No genes from our hormone-related list of interest were identified as cluster markers. In Fig. 4F, we plotted expression of these marker genes across DA clusters to confirm that they were selectively expressed. In the same heatmap, we also plotted feeding and social behavior hormone-related genes across DA clusters to visualize whether they were similarly selectively expressed or widely expressed across clusters.

##### Test for Significant Co-Expression

In Fig. 4J-K, we tested whether feeding- and social hormone-related genes within individual DA nuclei were co-expressed significantly more than chance. Chance co-expression was defined based on a null distribution constructed as follows. For each gene pair, we randomly reassigned expression of the genes across all DA nuclei, maintaining the overall number of nuclei expressing each gene. We then noted what percentage of nuclei with these random assignments expressed both genes. For every single gene pair, we iterated this process 10,000 times to construct a null distribution. The *p*-value for co-expression was defined as the percentile of the real data on this null distribution (Figure 4J-K). This process was repeated for GABA and glutamate nuclei in order to generate the percentage of gene pairs co-expressed more than chance in each neuronal subtype. In Fig. 4L, we compared co-expression between DA and GABA as well as DA and glutamate nuclei using two separate 2-sample tests for equality of proportions.

---

**Figure S1.**
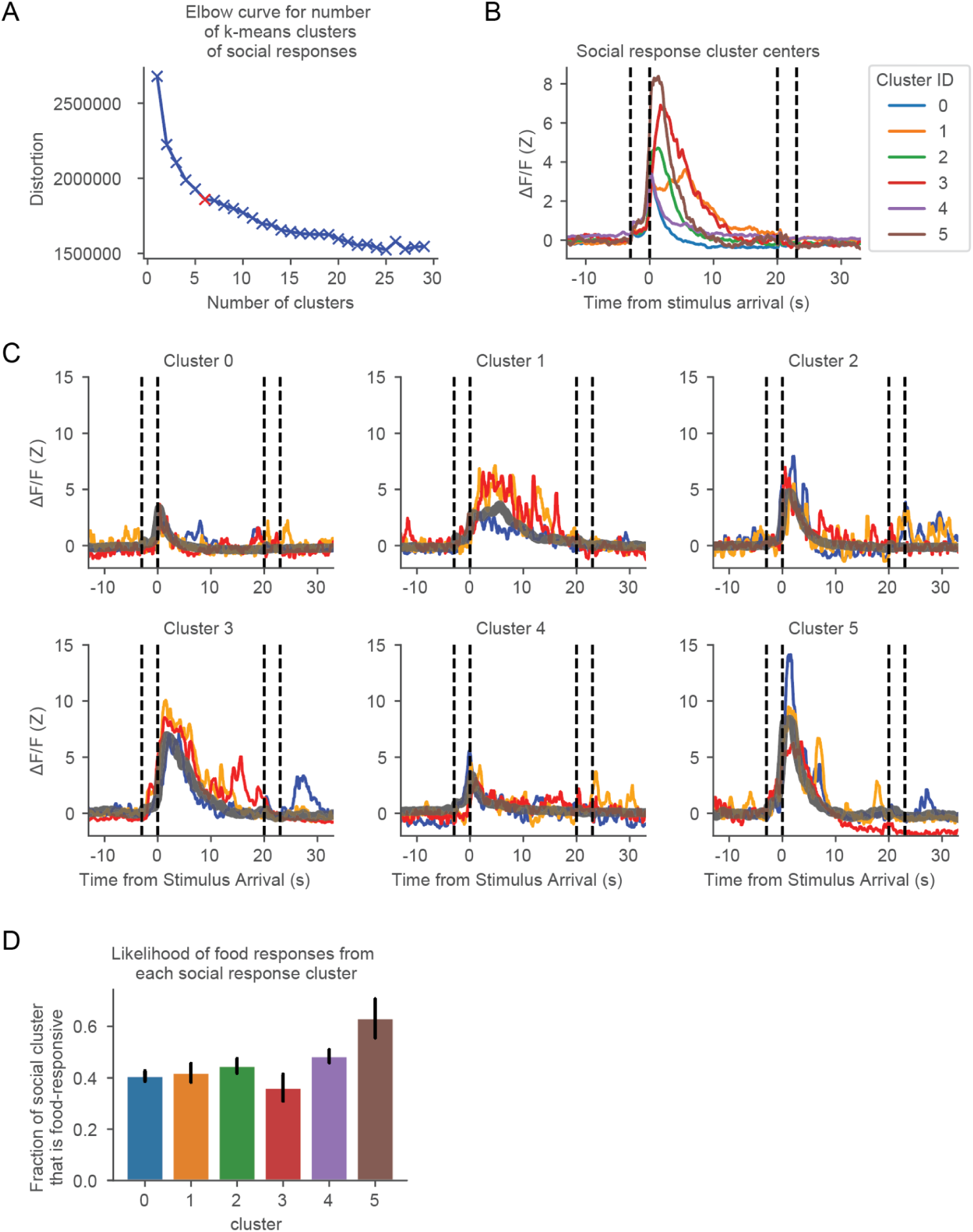
Social response profile diversity. (A) With K-means clustering of single neuron responses to male arrival, elbow plot of the distortion (inversely related to how well the dataset is clustered) versus number of clusters. Where the slope transitions from high to low (elbow at 6 clusters) signifies optimal number of clusters to fit the data. Data clustered is male-responsive neurons from Figure 1E. (B) Cluster centers from each of the 6 response profiles to male social stimuli. (C) Thin colored lines are randomly chosen single neuron responses within each cluster and bold gray lines show the cluster centers (same traces as in B). Fraction of neurons in each male-responsive cluster that are also food-responsive (error bars represent standard error from the binomial distribution of food-responsive neurons from each cluster).

**Figure S2.**
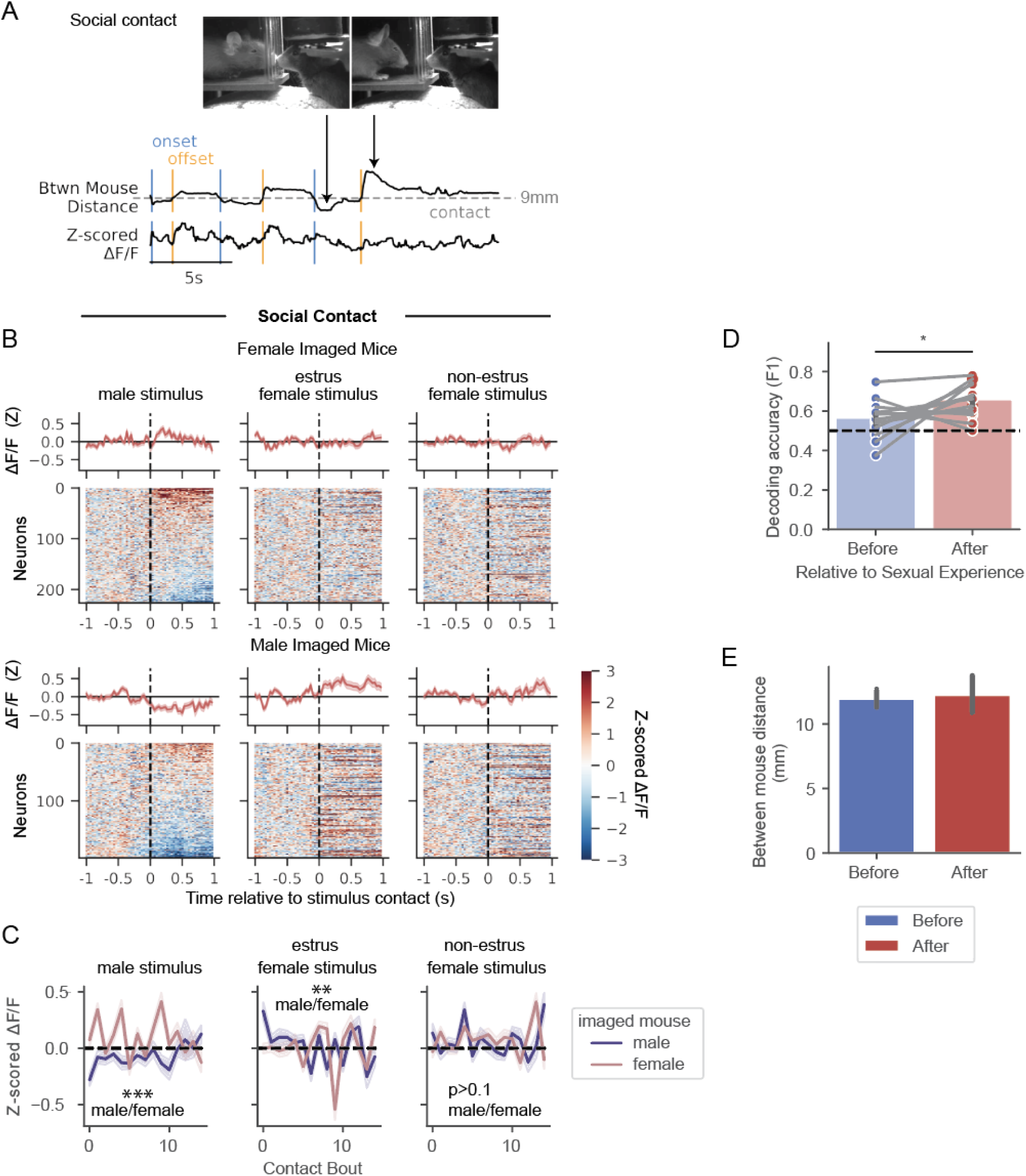
VTA^DA^ responses to conspecific contact differ between sexes. (A) Schematic of social contact, defined by the distance between the nose of the imaged mouse and eye of the social stimulus. (B) Neural responses to the first bout of social contact on a single baseline imaging session, separated by imaged mouse and social stimulus sex (N=8 imaged females with 210 neurons and 11 imaged males with 199 neurons). (C) Average activity across neurons recorded from male and female imaged mice (GEE for average activity by contact bout, imaged mouse sex, and the interaction between trial and imaged mouse sex, N=199 neurons from males, 210 neurons from females). (D) Decoding accuracy of same vs opposite sex contact based on neural ensemble responses, comparing before and after opposite sex experience (N=13 mice). Panels A-C show after opposite sex experience responses. (E) Distance between imaged and stimulus mice during the first 10 seconds of the first opposite sex stimulus delivery before and after opposite sex experience (N=9 males and 7 females before opposite sex experience, 11 males and 8 females after opposite sex experience). **See Table S7-10 for statistics.** ∗p < 0.05, ∗∗p < 0.01, ∗∗∗p < 0.001. Unless specified, data plotted as mean ± SEM.

**Figure S3.**
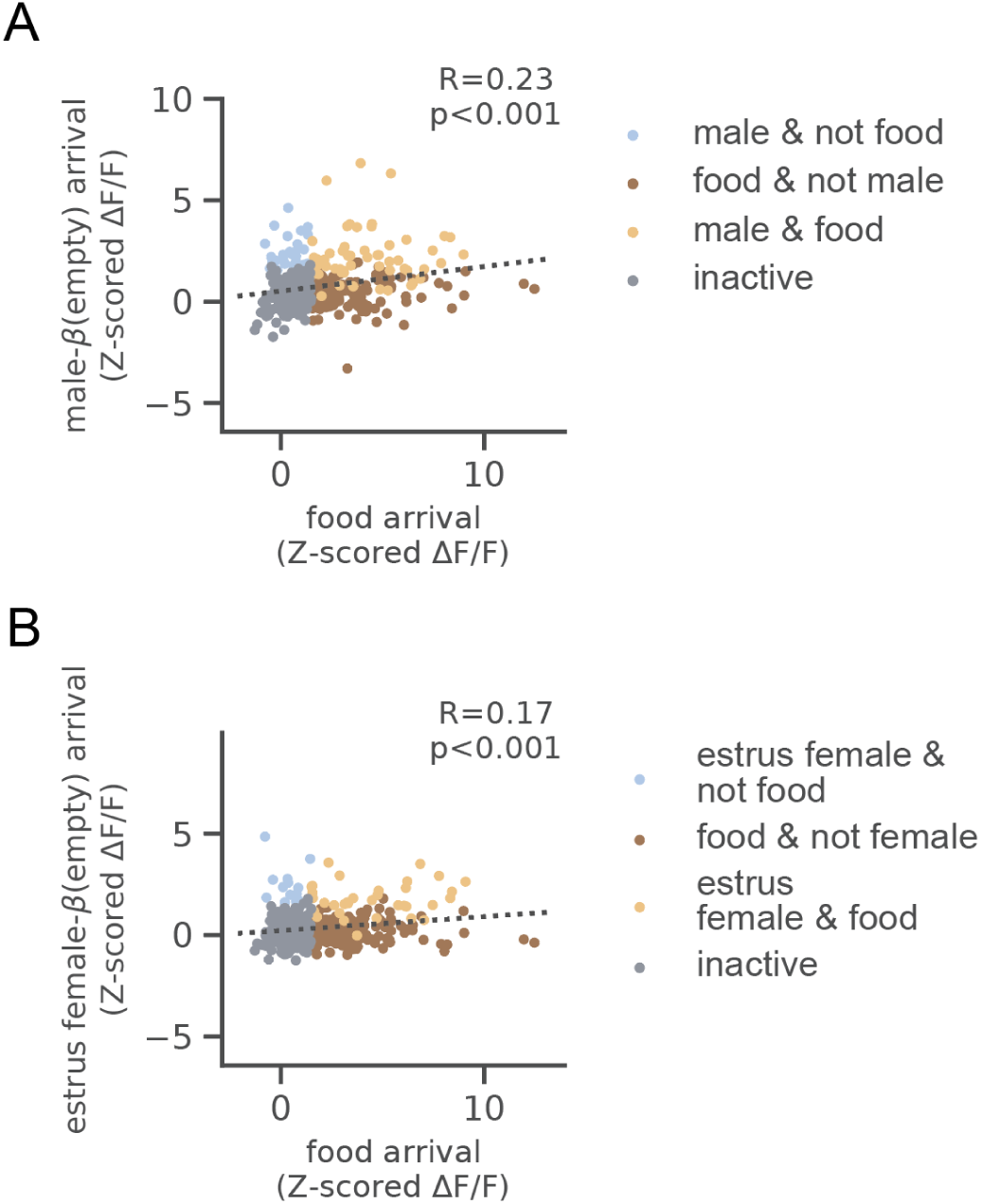
Correlation between VTA^DA^ responses to food and social stimuli, accounting for object responses. (A) Partial correlation between neural activity in response to food arrival and male arrival, accounting for object arrival; colored by selectivity to food arrival, male arrival, or both (in A-B, N=411 neurons). (B) Same as (A) for estrus female stimulus mice. **See Table S12 for statistics.**

**Figure S4.**
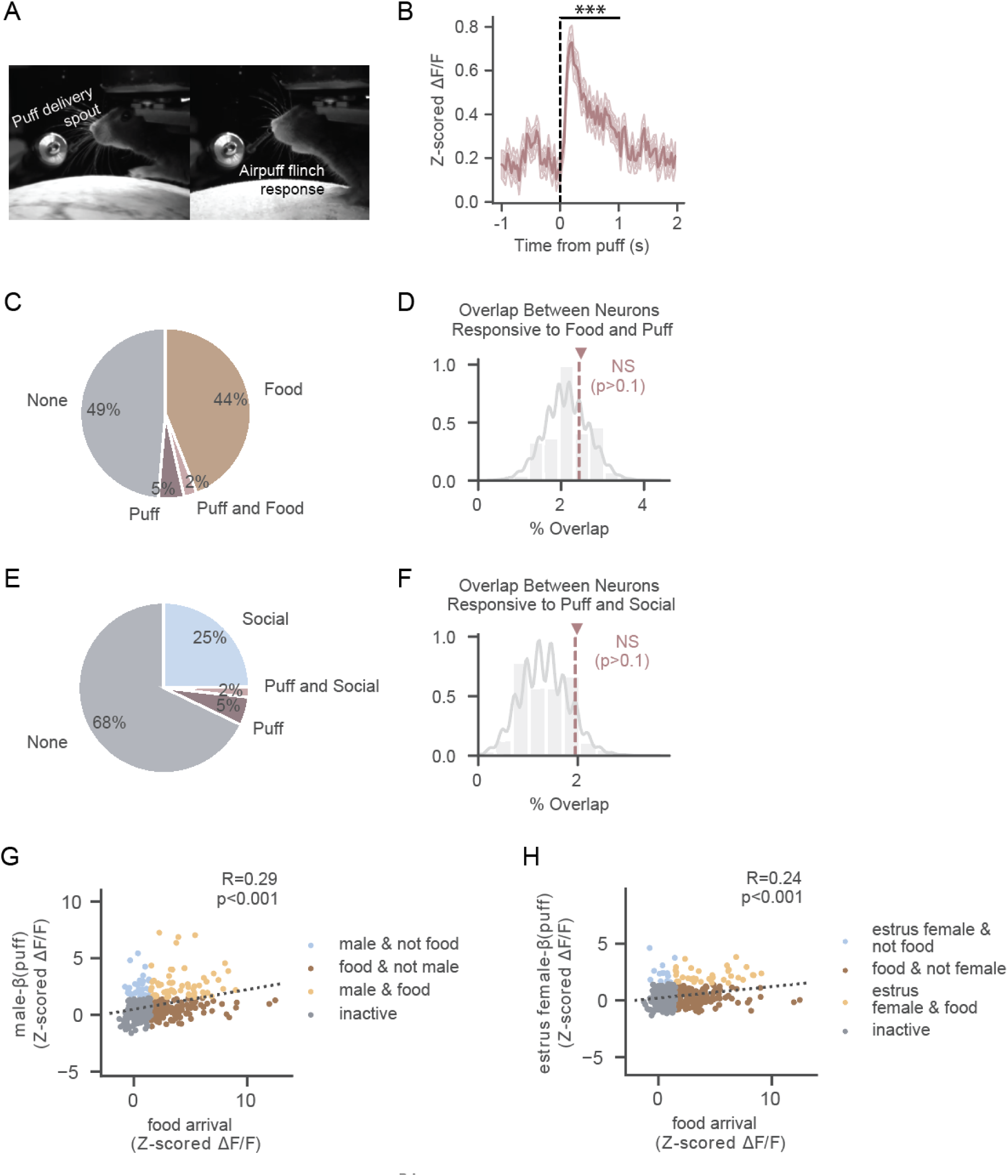
Negligible overlap in VTA^DA^ populations responsive to air puffs and food or social stimuli. (A) Example video frames of air-puff delivery setup before (left) and immediately after (right) puff to the snout. Air puffs were delivered at the end of recording sessions, following all food, social and object stimulus delivery. (B) Average response across all recorded neurons to the first air puff delivery. (In B-H, N=411 neurons) (C) Percentage of recorded neurons responsive to food arrival, air puff, both, or neither. Tuning specificity here and in subsequent panels is based on the mean activity in the first second after arrival or air puff. (D) Overlap compared to null distribution assuming food- and puff-responsive neurons are independent samples. (E) Same as (C) for neurons responsive to social stimuli (male and/or estrus female) and air puff. (F) Same as (D) for neurons responsive to social stimuli (male and/or estrus female) and air puff. (G) Partial correlation between neural activity in response to food arrival and male arrival, accounting for air puff response; colored by selectivity to food arrival, male arrival, or both. (H) Same as (G) for estrus female stimulus mice. **See Table S13 for statistics.** ∗p < 0.05, ∗∗p < 0.01, ∗∗∗p < 0.001. Unless specified, data plotted as mean ± SEM.

**Figure S5:**
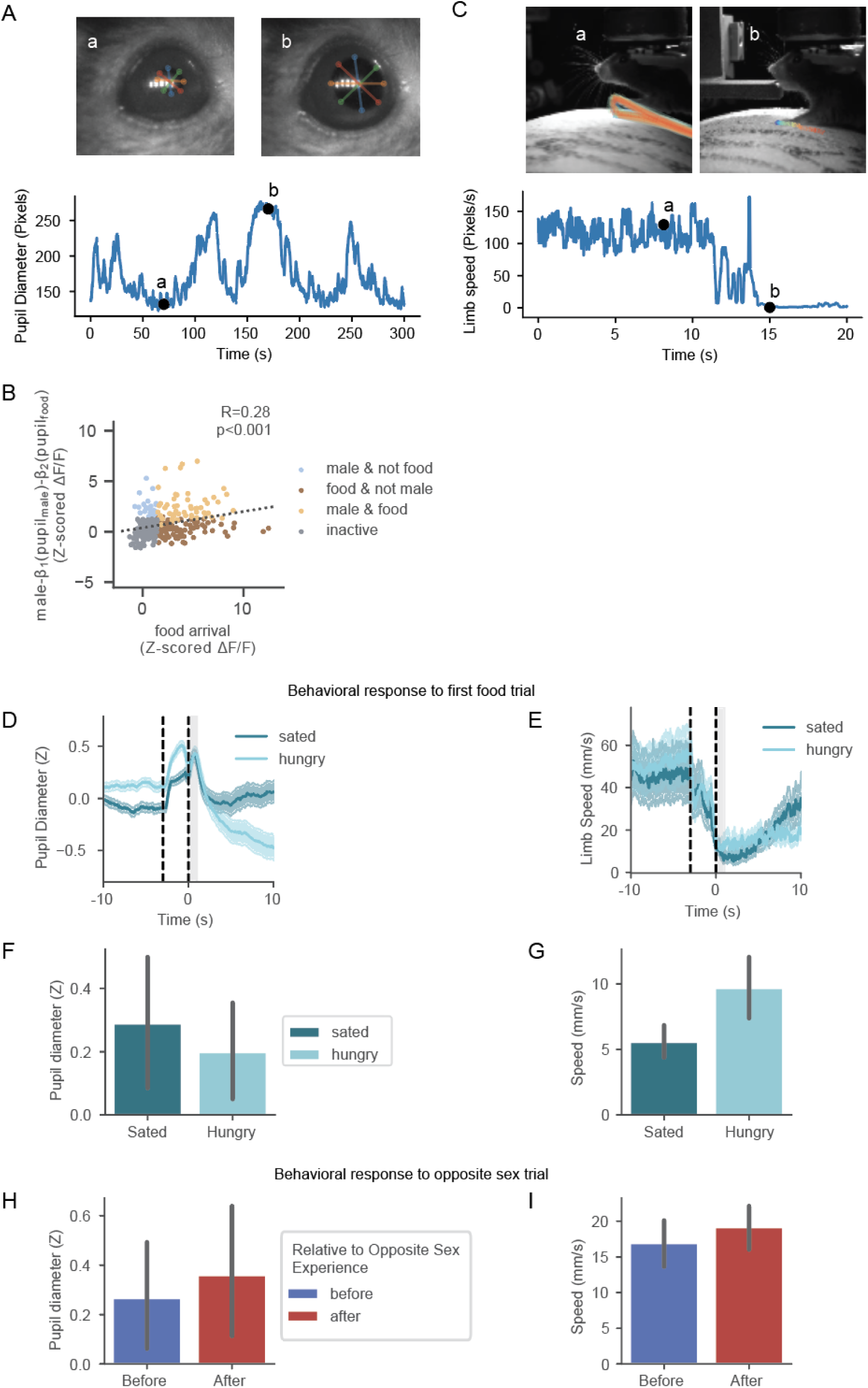
Pupil diameter and locomotion as measures of arousal. (A) Top: Example frames from pupil-focused camera with post-hoc diameter tracking overlaid (top). Bottom: Example trace of estimated diameter (mean of 4 tracked diameters) with example frames labeled. (B) Partial correlation between neural activity in the first second after food arrival and male arrival, accounting for pupil dilation response (Z-scored pupil diameter in first second after the first food or male trial); colored by neurons’ selectivity to food arrival, male arrival, or both. (N=306 neurons) (C) Top: Example frames from body-focused camera with a 3-second trajectory of post-hoc forepaw tracking overlaid (top). Bottom: Example trace of instantaneous forelimb speed with example frames labeled. (D) Mean z-scored pupil diameter aligned to first food trial, separated by hunger state (N=20 sated mice, 18 hungry mice in D-G). (E) Mean limb speed aligned to the first food trial, separated by hunger state. (F) Comparison of pupil dilation across hunger states, in the first second following food delivery (average during gray interval from D). (G) Comparison of limb speed across hunger states, in the first second following food delivery (average during gray interval from E). (H) Comparison of pupil dilation before and after opposite sex experience, in the first second following the first opposite sex stimulus delivery (N=16 mice before and 20 mice after for H-I). (I) Comparison of limb speed before and after opposite sex experience, in the first second following the first opposite sex stimulus delivery. **See Table S514 for statistics.**

**Figure S6.**
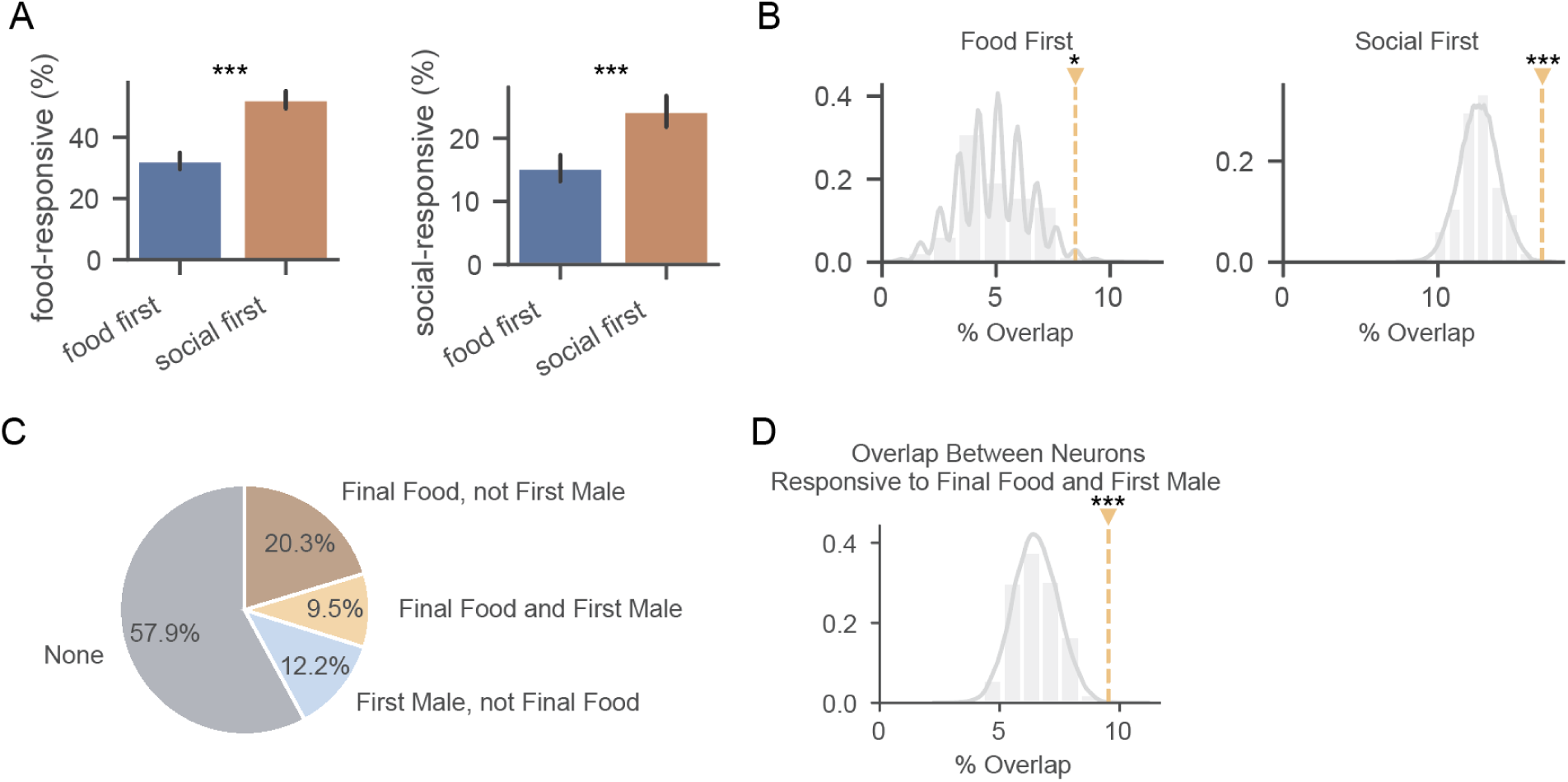
Order of stimuli changes responsiveness but food and social overlap is significant regardless of order. (A) Fraction of neurons that are responsive to food or male (social) stimuli on the first presentation of each, split by which stimulus arrived before the other one in the session (in A-B, N=411 neurons, same neurons, mice and session as in Figures 1-2). (B) Overlap compared to null distribution assuming food- and male-responsive neurons are independent samples, split by if the food or male stimulus arrived before the other stimulus type in the session. (C) Percentage of recorded neurons responsive to last food arrival, first male male arrival, both, or neither. Tuning specificity here and in subsequent panels is based on the mean activity in the first second after arrival. (D) Overlap compared to null distribution assuming last-food- and first-male-responsive neurons are independent samples. **See Table S15 for statistics.** ∗p < 0.05, ∗∗p < 0.01, ∗∗∗p < 0.001. Unless specified, data plotted as mean ± SEM.

**Figure S7.**
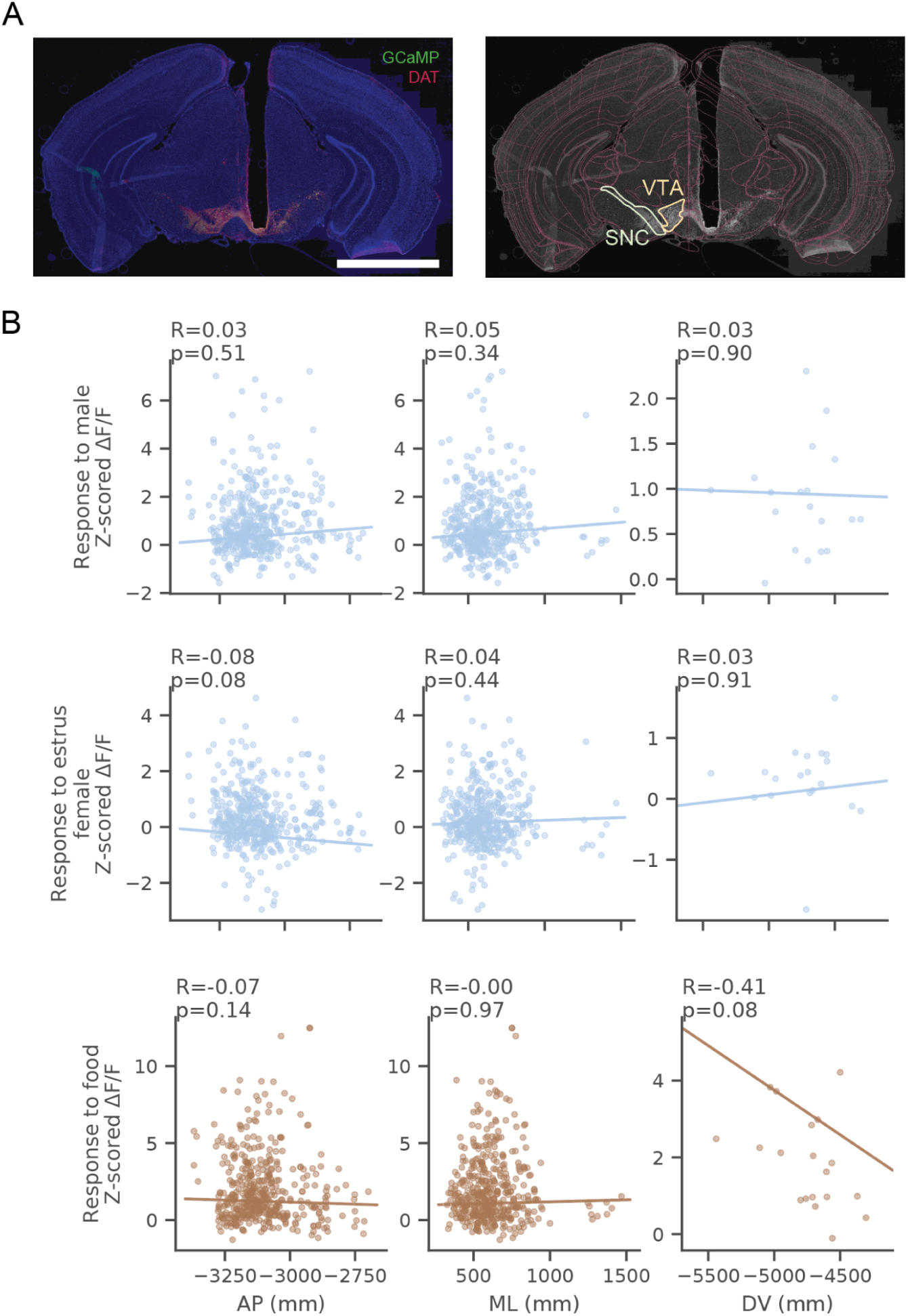
No relationship between anatomical location of neurons and responses to social stimuli or food. (A) Example histological image GRIN lens implant over the VTA, stained for DAT+ and expressing GCaMP6f in DAT neurons (left). Example registration of the brain slice to common Allen Atlas reference coordinates using WholeBrain (Fürth et al. 2018) (right). (B) Relationships between anterior-posterior (AP), medial-lateral (ML), and dorsal-ventral (DV) locations of cells (for AP, ML) and fields of view (for DV) and magnitudes of responses to male (top), estrus female (middle), and food (bottom) stimuli (Left, middle: N=435 neurons; Right: N=19 fields). **See Table S16 for statistics.**

**Figure S8.**
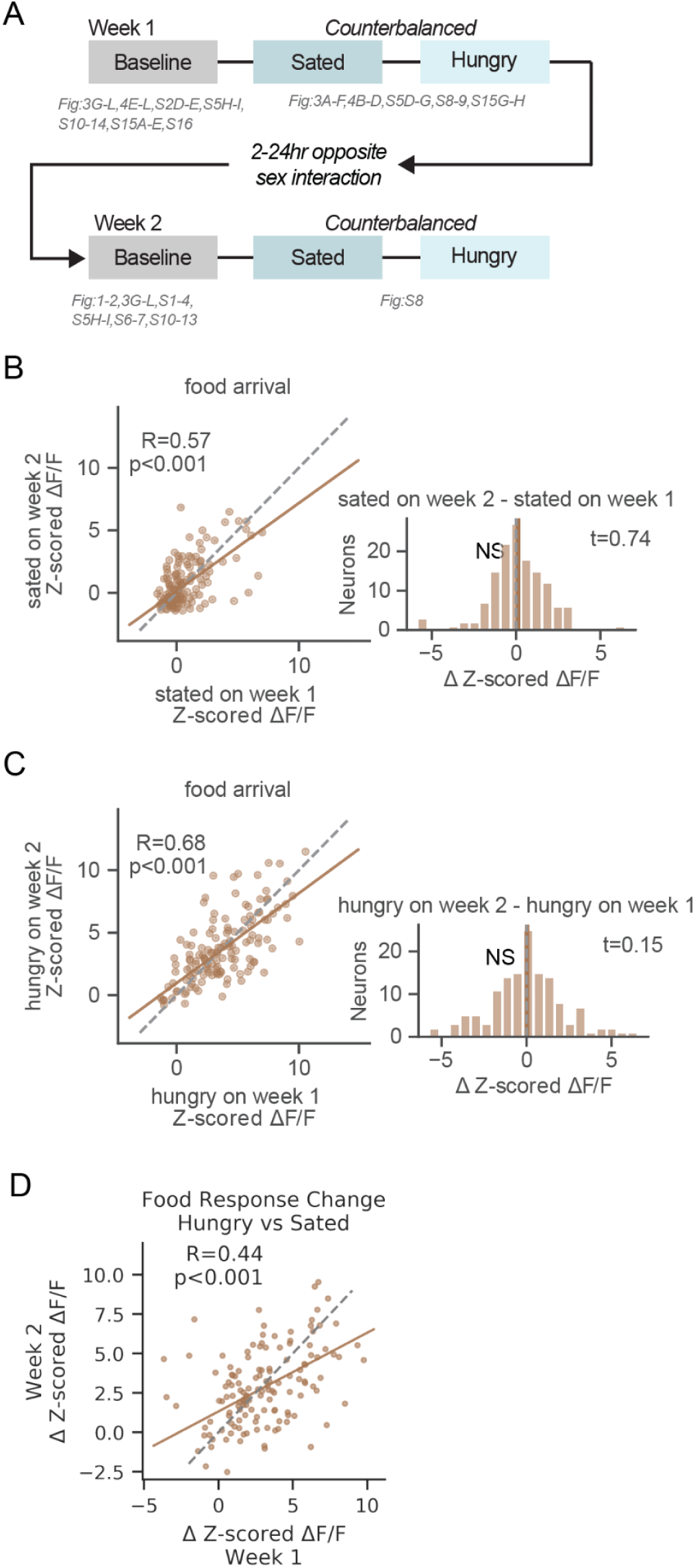
Consistent food responses across different days with the same sated or hunger states. (A) Timeline of change in food access and change in social experience experiments. (B) Relationship between neural activity in response to food arrival on first and second sated sessions (left). Distribution of change in activity in response to second to first sated sessions (right). (In all panels, N=137 neurons). (C) Relationship between neural activity in response to food arrival on first and second hungry sessions (left). Distribution of change in activity in response to second to first hungry sessions (right). (D) Relationship between neural activity changes between sated to hungry sessions on the first versus second week of imaging. **See Table S18 for statistics.** ∗p < 0.05, ∗∗p < 0.01, ∗∗∗p < 0.001.

**Figure S9.**
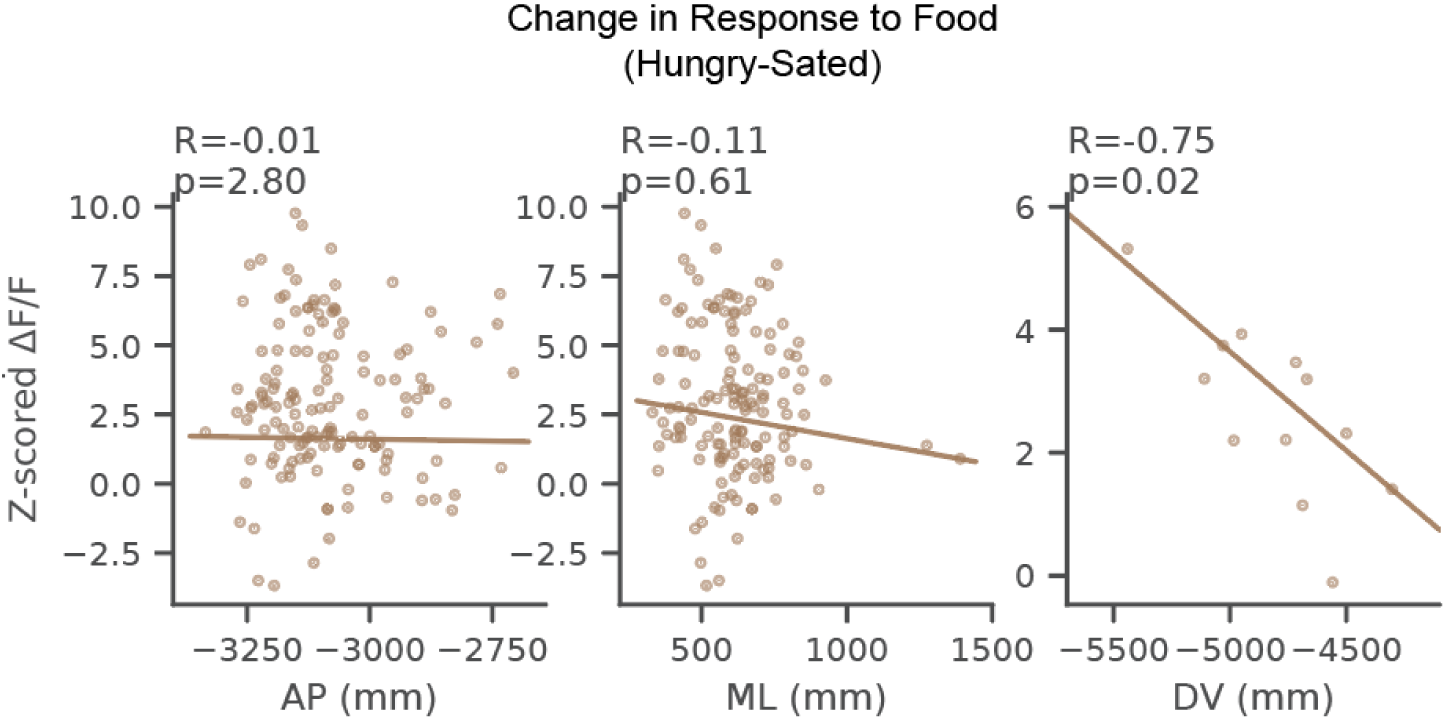
Change in magnitude of response to food when hungry is higher in neurons located in ventral VTA. Relationship between change in response to food between sated and hungry days and anatomical location of imaged neurons and fields of view. Left: anterior-posterior (A), middle: medial-lateral (ML), right: dorsal-ventral. All coordinates in Allen Atlas reference frame. (left, middle: N=140 neurons, right: N=12 fields). **See Table S19 for statistics.**

**Figure S10.**
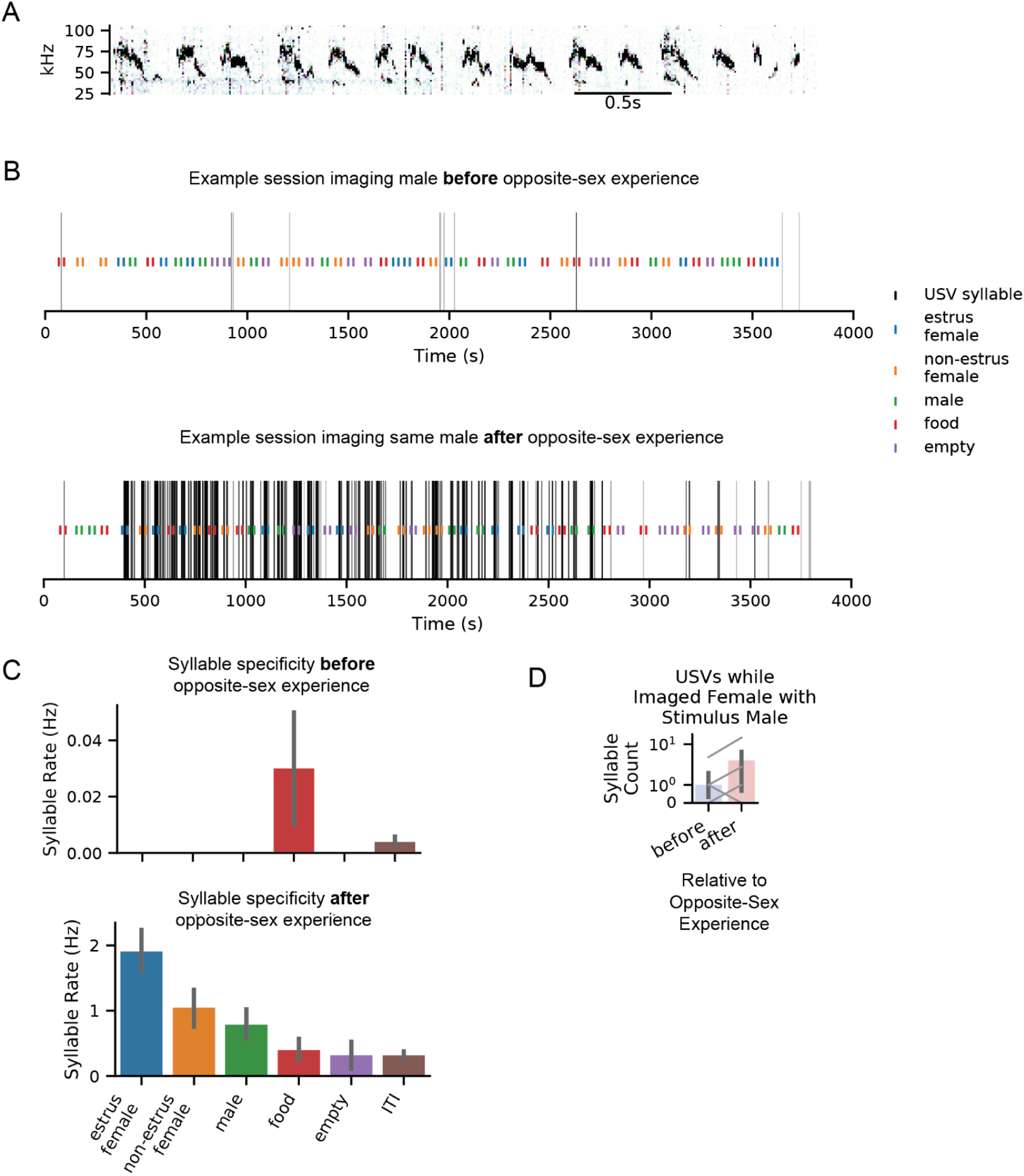
More ultrasonic vocalizations during opposite-sex presentation before versus after opposite-sex experience in imaged males. (A) Example spectrogram from raw recordings of vocal syllables in the ultrasonic frequency range. (B) From an example male, imaging sessions before and after opposite-sex experience. Black lines show where USV syllables were detected aligned to presentation of different stimulus types. Colored ticks indicated the onset and offset of stimulus presentation. (C) Average syllable vocalization rate during presentation of different stimuli in the example sessions shown in (B). (D) USV syllable counts while females are presented with male stimuli after versus before opposite sex experience (3 out of 5 females increased in USVs detected during male trials).

**Figure S11.**
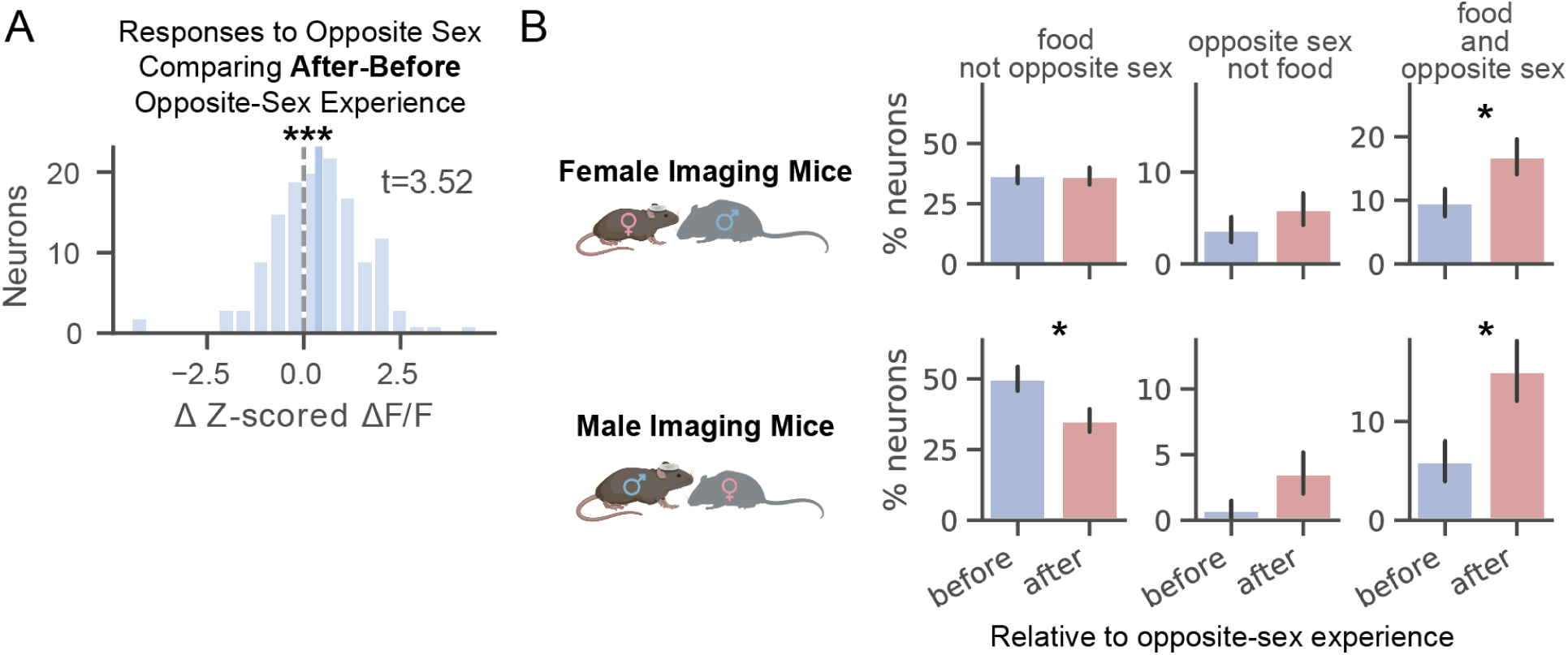
In males and females, opposite-sex experience increases fraction of neurons responding to both food and social stimuli. (A) Across all neurons recorded on both days, distribution of change in activity in response to opposite-sex arrival before and after opposite sex experience (N=137 neurons). (B) Comparison of percentage of neurons responsive to food arrival, opposite-sex arrival, and food and opposite-sex arrival before versus after freely moving opposite-sex interactions. Female imaging mice shown in top row, males in bottom (error bars show stdev of binomial distribution, females: N=187 neurons before, 184 neurons after, males: N=134 neurons before, 139 neurons after). **See Table S20 for statistics.**

**Figure S12.**
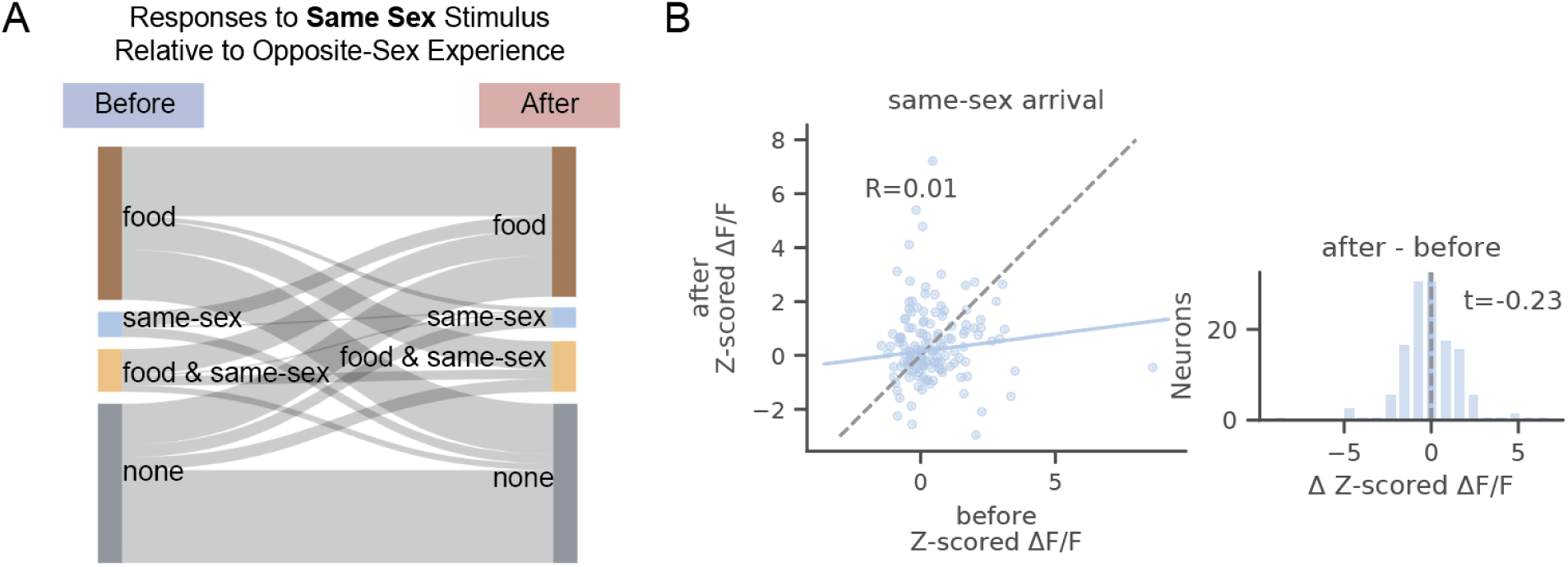
Opposite-sex experience does not change responses to same-sex stimuli. (A) Changes in neural tuning (responsive to food arrival, same-sex arrival, both, or neither) before versus after opposite sex interaction (N=227 neurons). (B) Relationship between neural activity in response to same-sex arrival on before versus after same-sex experience (left). Distribution of change in activity in response to same-sex arrival (right) (N=137 neurons). **See Table S21 for statistics.**

**Figure S13.**
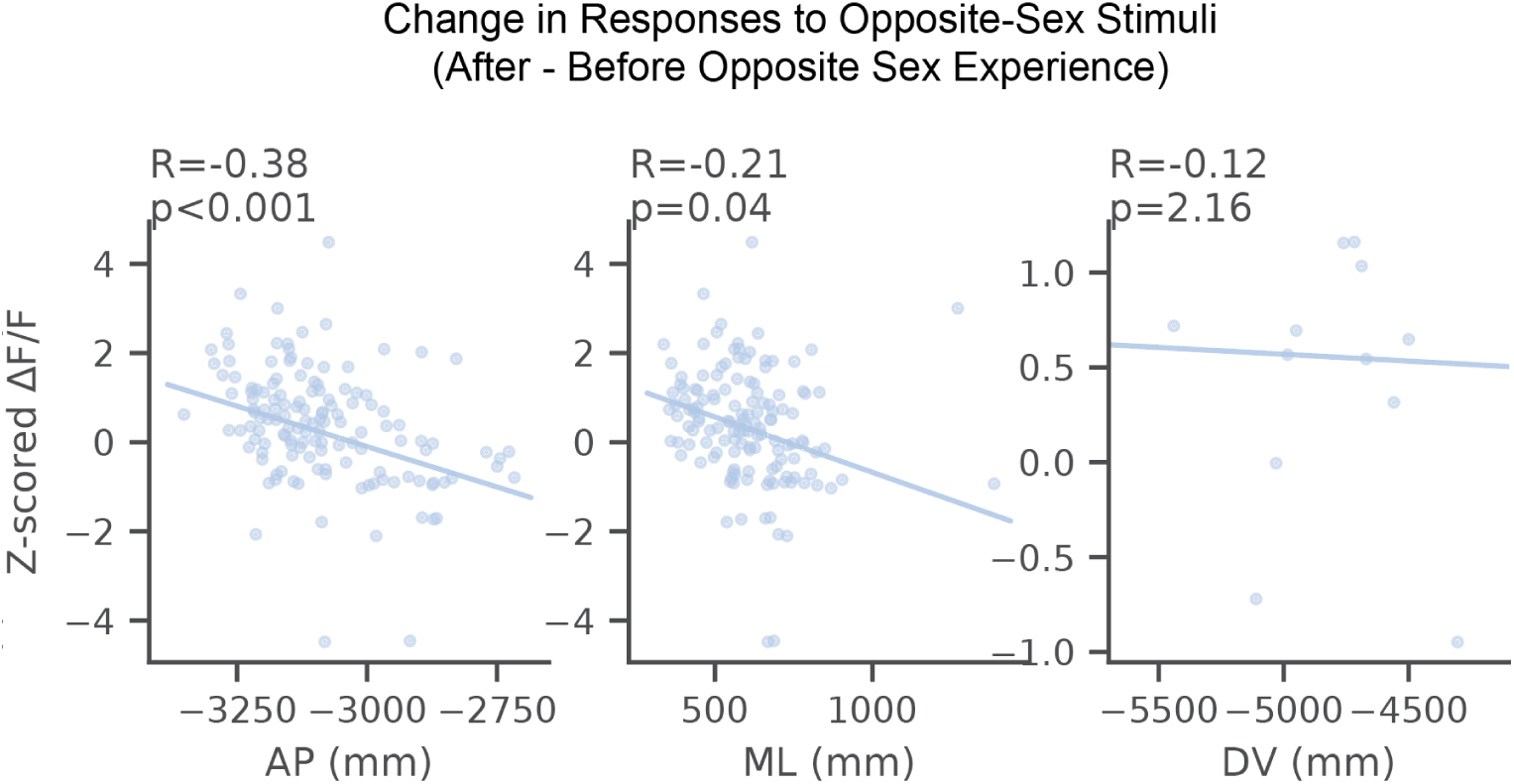
Change in magnitude of response to opposite-sex stimuli after opposite sex experience is higher in neurons located in more posterior and medial VTA. Relationship between change in response to opposite-sex stimuli between after and before opposite sex experience and anatomical location of imaged neurons and fields of view. Left: anterior-posterior (A), middle: medial-lateral (ML), right: dorsal-ventral. All coordinates in Allen Atlas reference frame (left, middle: N=137 neurons, right: N=12 fields). **See Table S22 for statistics.**

**Figure S14.**
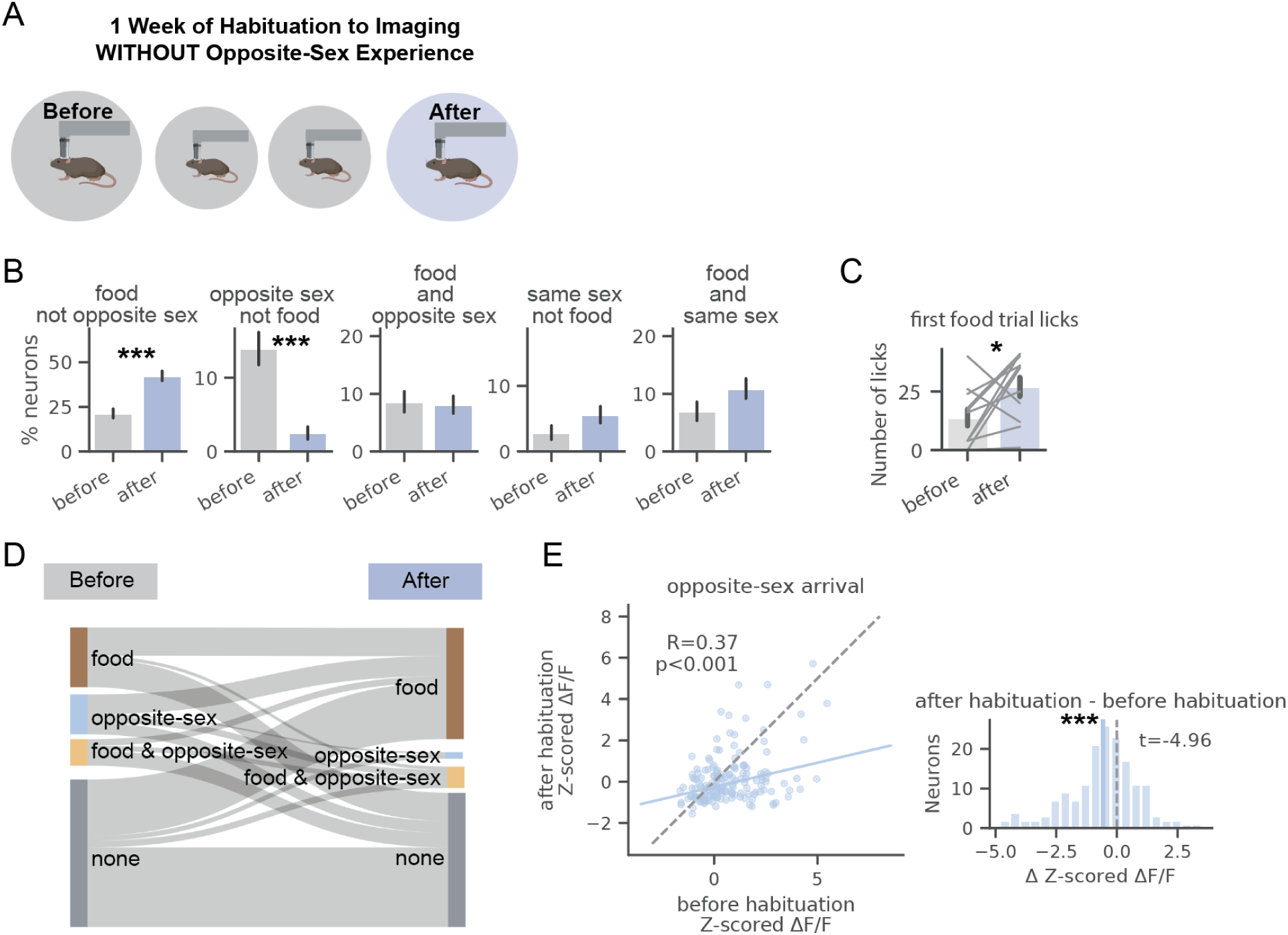
Habituation to imaging and stimulus delivery decreases opposite-sex responses. (A) Cartoon representing comparison of imaging before versus after habituation. (B) Comparison of percentage of neurons responsive to food arrival, opposite-sex arrival, food and opposite-sex arrival, same-sex arrival, and food and same-sex arrival both before versus after habituation (error bars show standard deviation of binomial distribution, before: N=244 neurons, after: N=408 neurons). (C) Lick rates during the first trial of food presentation before vs after habituation (N=11 mice). (D) Changes in neural tuning (responsive to food arrival, opposite-sex arrival, both, or neither) before versus after habituation (N=165 neurons). (E) Relationship between neural activity in response to opposite-sex arrival on before versus after habituation (left). Distribution of change in activity in response to opposite-sex arrival (right). (N=165 neurons). **See Table S23 for statistics.**

**Figure S15.**
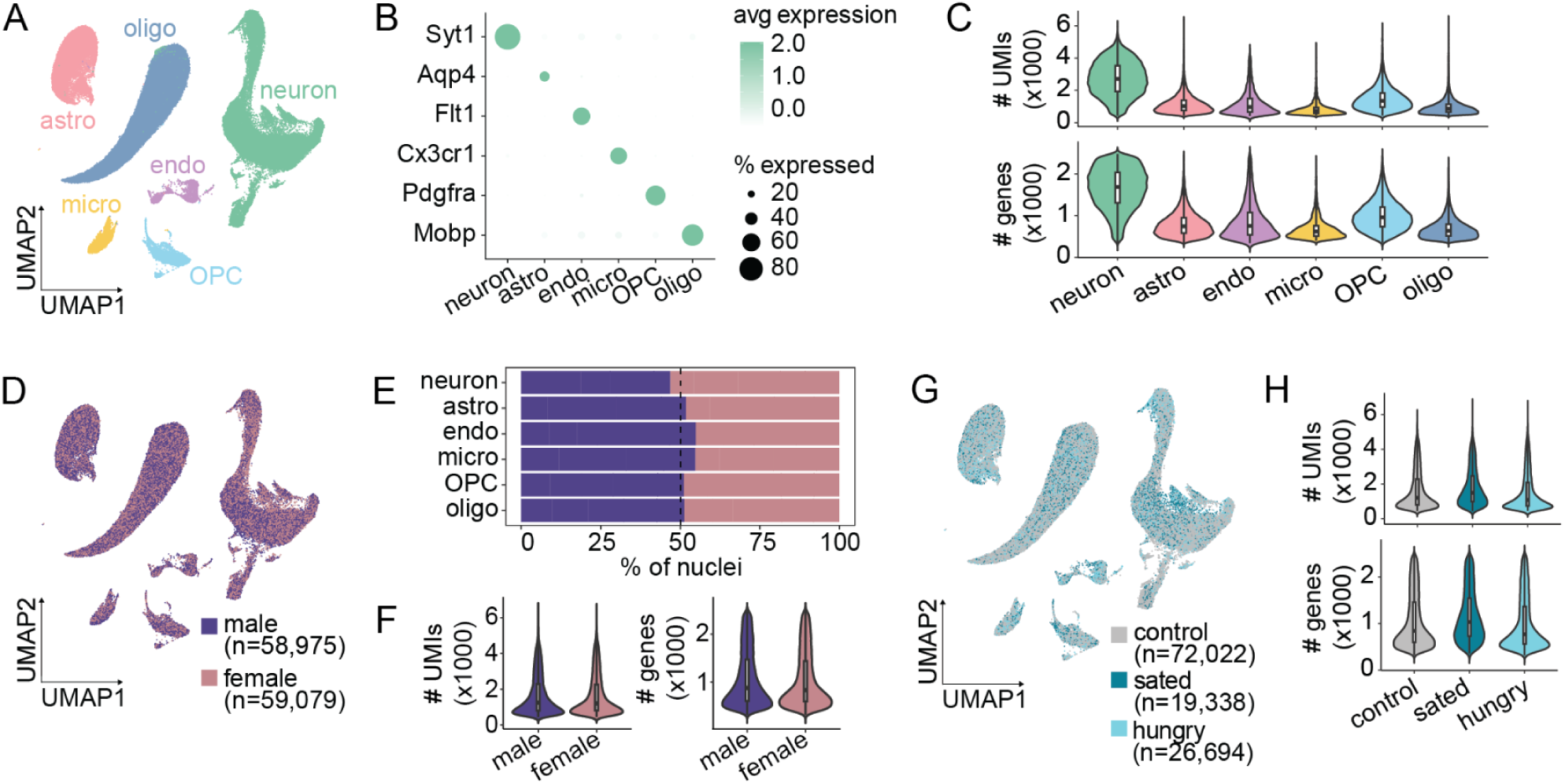
Quality control measurements for snRNA-seq of mouse VTA. (A) Uniform approximation and projection (UMAP) of all VTA nuclei with points colored by cell type classification (N=118,054 nuclei). (B) Dot plot showing the percentage of nuclei in each cell type expressing known marker genes (size of dot) as well as the average level of expression of each marker gene (color of dot) across clusters. (C) Violin plots showing the unique molecular identifier (UMI; top) and gene (bottom) distributions in each cell type. (D) UMAP of all VTA nuclei with points colored by sex of the sample they are from (N=58,975 nuclei from male samples and N=59,079 nuclei from female samples for E-F). (E) Percentage of nuclei in each cell type from male or female samples. (F) Violin plots showing the UMI (left) and gene (right) distributions in male versus female samples. (G) UMAP of all VTA nuclei with points colored by the hunger state condition of the sample they are from (N=72,022 nuclei from control animals, N=19,338 nuclei from sated (pre-fed) animals, and N=26,694 nuclei from hungry animals for G-H). (H) Violin plots showing the UMI (left) and gene (right) distributions in control, sated, and hungry samples.

**Figure S16.**
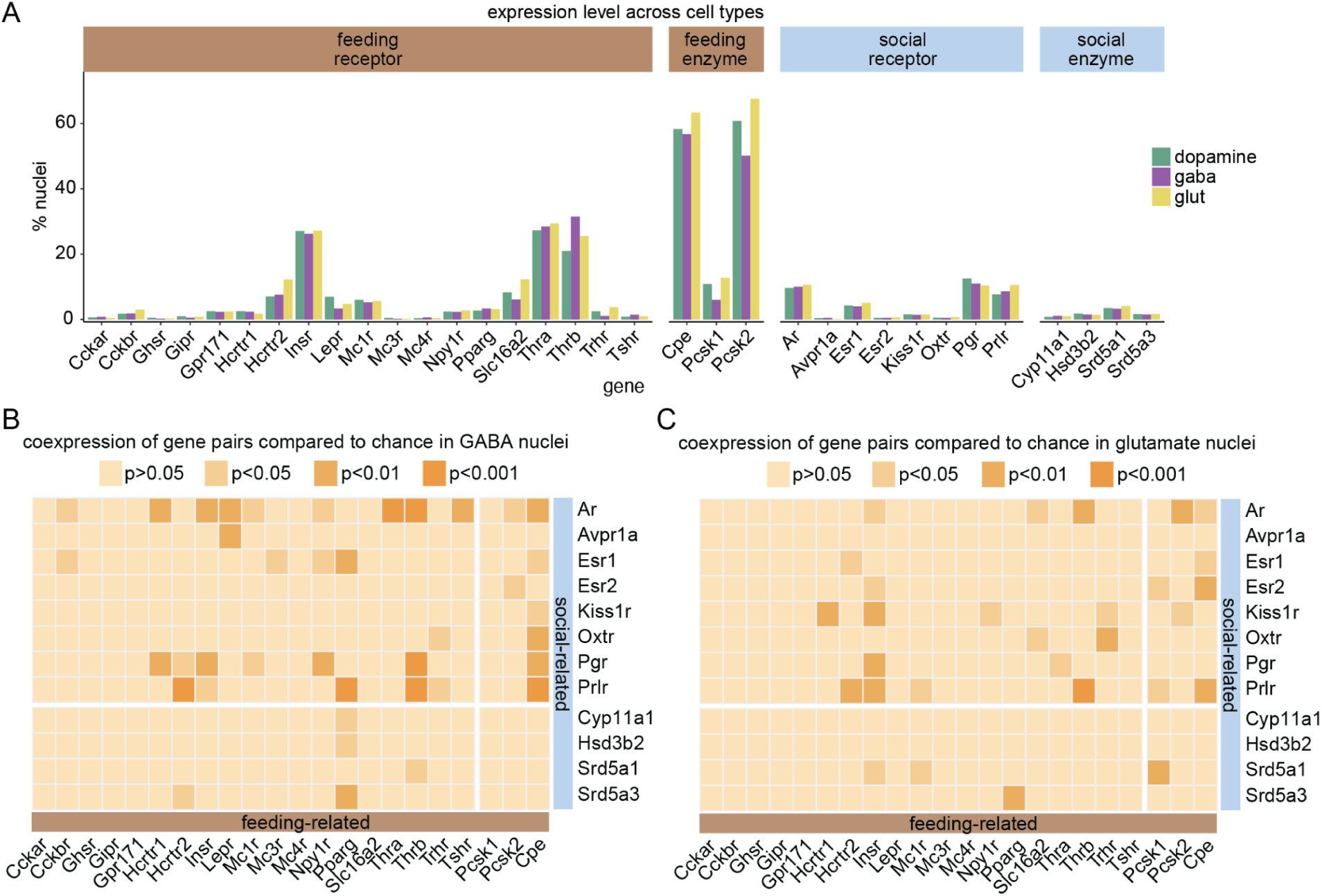
Expression of feeding- and social-related genes across neuron subtypes. (A) Percentage of nuclei expressing each feeding and social hormone-related gene in each neuronal subtype. (B) Matrix showing the level of co-expression (compared to chance, assuming the nuclei expressing each gene are independent samples from the full population; see Methods) of each gene pair in GABA nuclei. (C) Matrix showing the level of co-expression (compared to chance, assuming the nuclei expressing each gene are independent samples from the full population; see Methods) of each gene pair in glutamate nuclei.

## Supplementary Tables

**Table S1:**
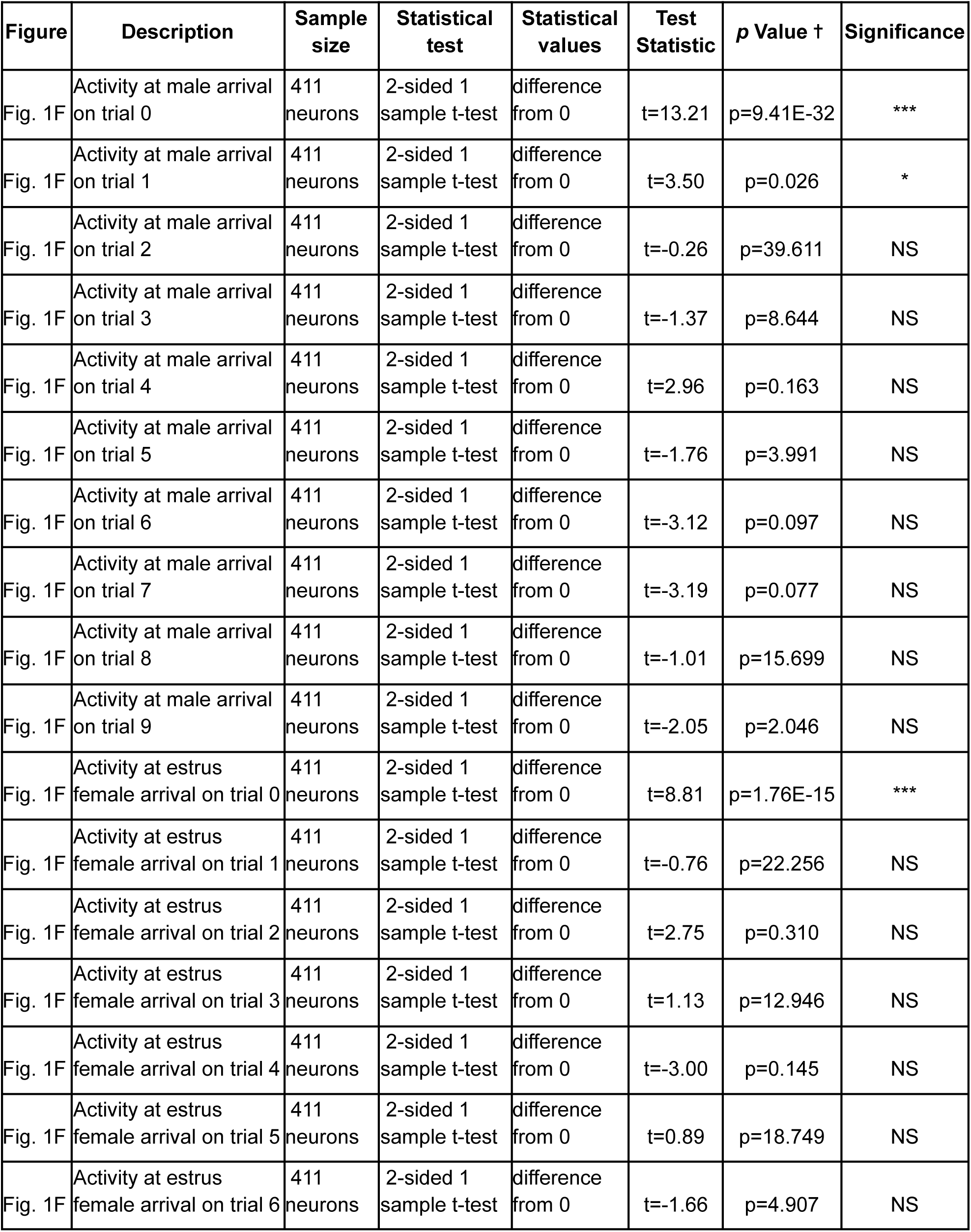
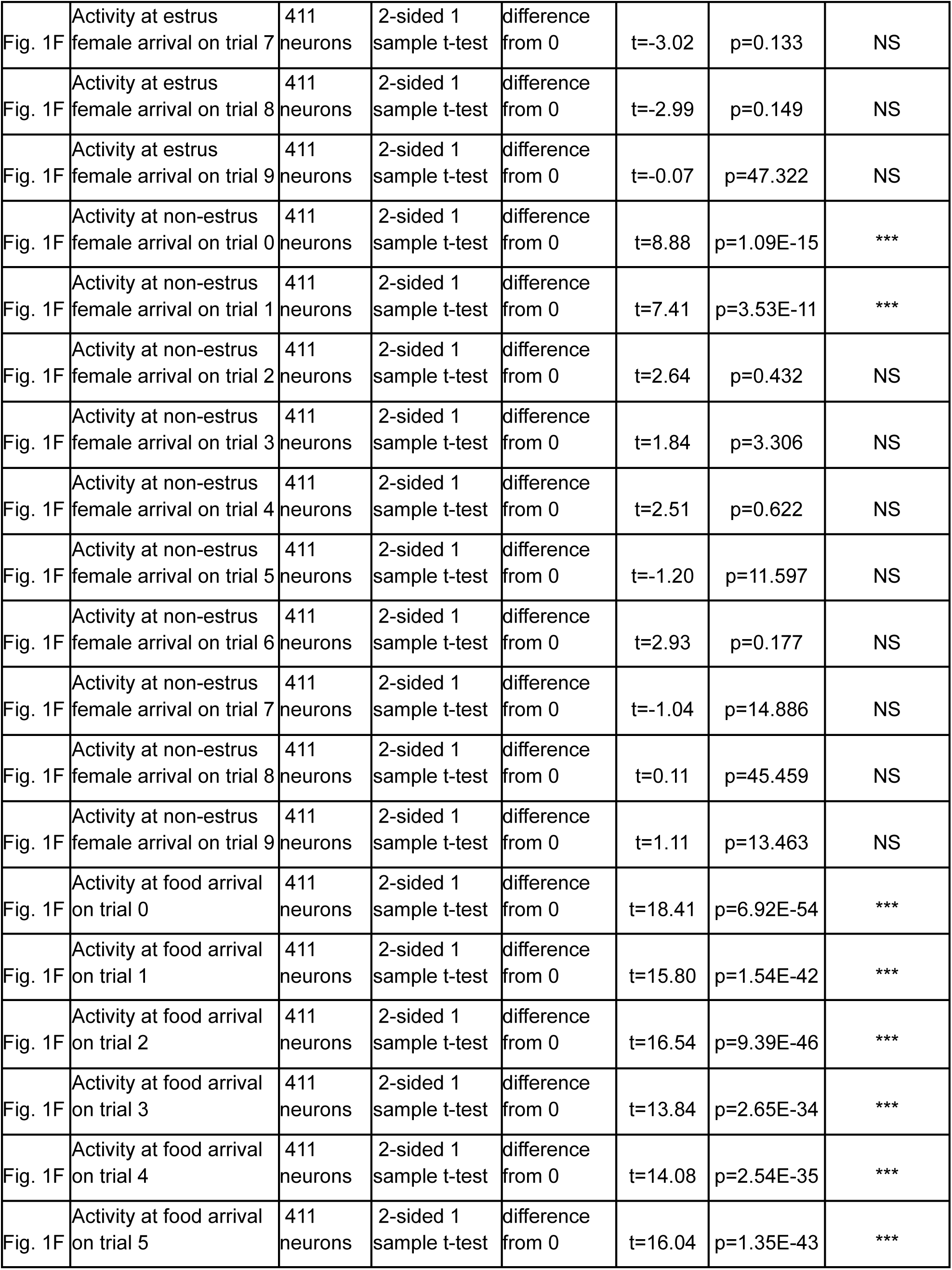
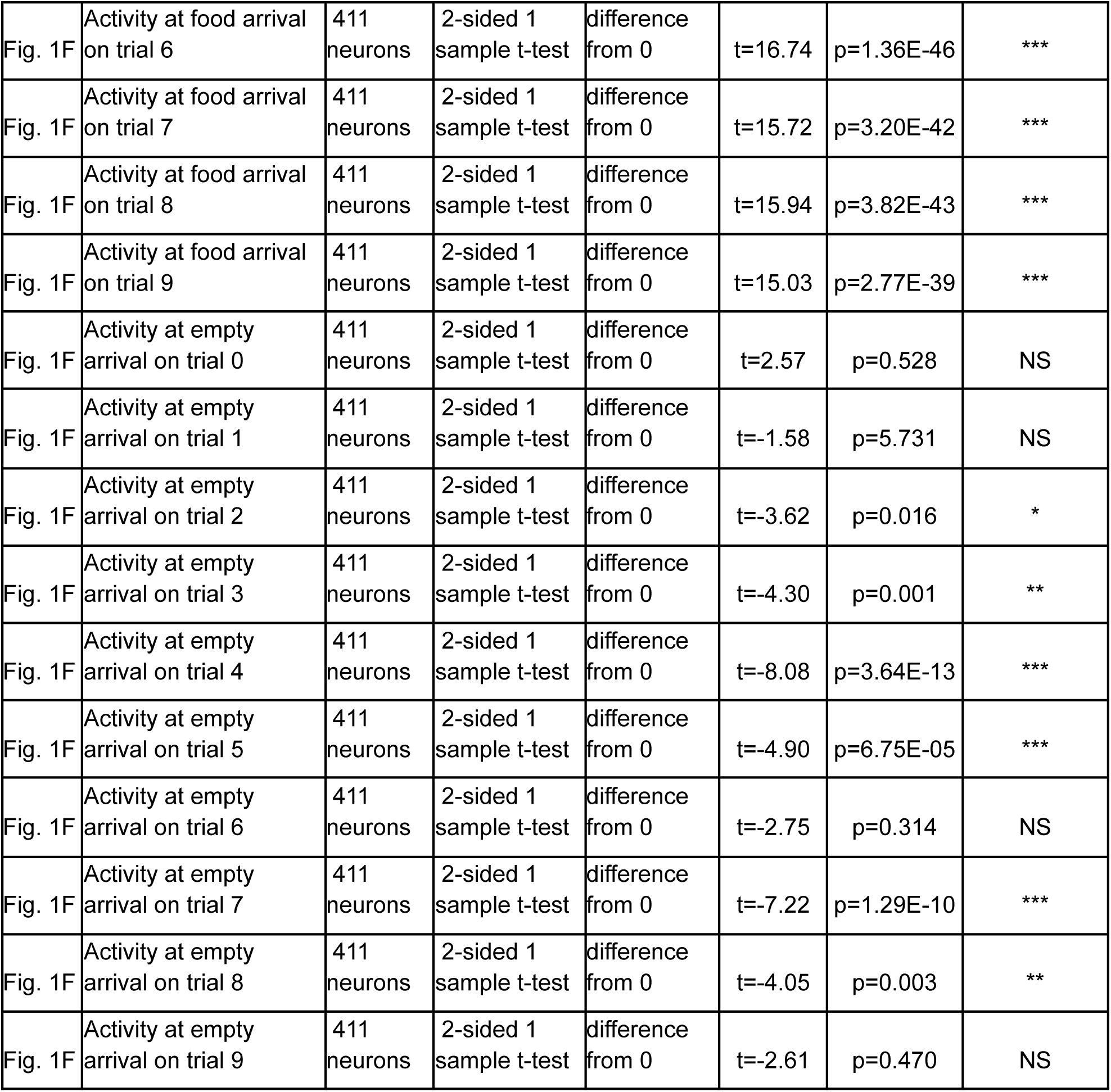
Statistics for Figure 1 (✝ Bonferroni correction for 50 tests)

**Table S2:** 2-sided GEE regression of GCaMP response to male (Figure 1F): *Z-scored ΔF/F = trial + sex + sex*trial + intercept, grouped by cell*. Number of cells (groups) = 411, minimum samples per group 199, maximum samples per group 212, dependence structure = independence, family = Gaussian. P-values Bonferroni corrected for 5 comparisons.

| *Model: Z-scored $\Delta F/F$ responding to male = trial + sex + trial\*sex + intercept, grouped by cell* | $\beta \pm$ standard error | z-stat | p-value | 95% CI [lower, upper] | |
| --- | --- | --- | --- | --- | --- |
| **Intercept** | 0.291±0.041 | 7.025 | 1.07E-11 | 0.21 | 0.372 |
| **sex** | 0.102±0.07 | 1.446 | 0.74 | -0.036 | 0.239 |
| **trial** | -0.053±0.005 | -10.125 | 2.13E-23 | -0.063 | -0.042 |
| **sex \* trial** | -0.018±0.009 | -2.044 | 0.205 | -0.035 | -0.001 |

**Table S3:**
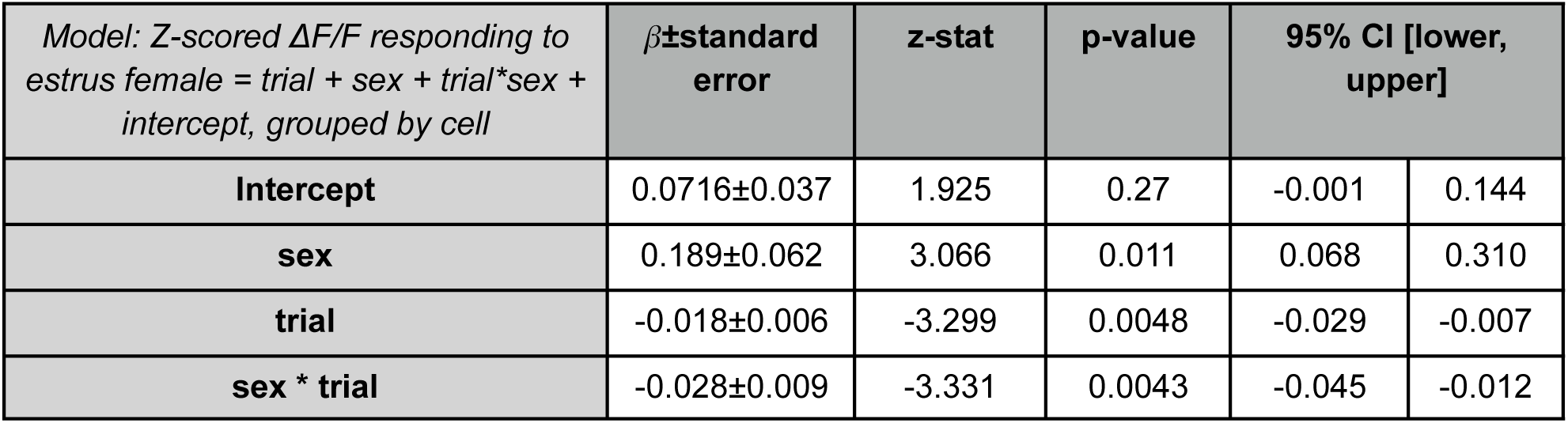
2-sided GEE regression of GCaMP response to estrus female (Figure 1F): *Z-scored ΔF/F = trial + sex + sex*trial + intercept, grouped by cell*. Number of cells (groups) = 411, minimum samples per group 199, maximum samples per group 212, dependence structure = independence, family = Gaussian. P-values Bonferroni corrected for 5 comparisons.

| *Model: Z-scored $\Delta F/F$ responding to estrus female = trial + sex + trial\*sex + intercept, grouped by cell* | $\beta \pm \text{standard error}$ | z-stat | p-value | 95% CI [lower, upper] | |
| --- | --- | --- | --- | --- | --- |
| **Intercept** | 0.0716±0.037 | 1.925 | 0.27 | -0.001 | 0.144 |
| **sex** | 0.189±0.062 | 3.066 | 0.011 | 0.068 | 0.310 |
| **trial** | -0.018±0.006 | -3.299 | 0.0048 | -0.029 | -0.007 |
| **sex \* trial** | -0.028±0.009 | -3.331 | 0.0043 | -0.045 | -0.012 |

**Table S4:**
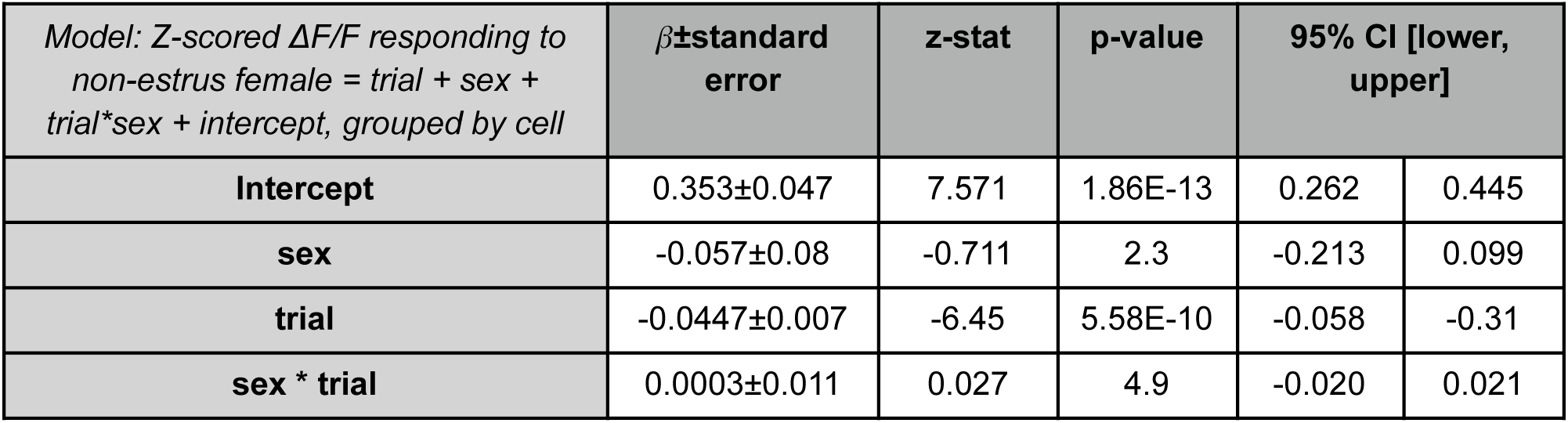
2-sided GEE regression of GCaMP response to non-estrus female (Figure 1F): *Z-scored ΔF/F = trial + sex + sex*trial + intercept, grouped by cell*. Number of cells (groups) = 411, minimum samples per group 199, maximum samples per group 212, dependence structure = independence, family = Gaussian. P-values Bonferroni corrected for 5 comparisons.

**Table S5:** 2-sided GEE regression of GCaMP response to food (Figure 1F): *Z-scored ΔF/F = trial + sex + sex*trial + intercept, grouped by cell*. Number of cells (groups) = 411, minimum samples per group 199, maximum samples per group 212, dependence structure = independence, family = Gaussian. P-values Bonferroni corrected for 5 comparisons.

| *Model: Z-scored $\Delta F/F$ responding to food = trial + sex + trial\*sex + intercept, grouped by cell* | $\beta \pm \text{standard error}$ | z-stat | p-value | 95% CI [lower, upper] | |
| --- | --- | --- | --- | --- | --- |
| **Intercept** | 1.452±0.108 | 13.41 | 2.64E-40 | 1.24 | 1.66 |
| **sex** | 0.414±0.175 | 2.37 | 0.089 | 0.072 | 0.756 |
| **trial** | -0.0477±0.012 | -4.123 | 1.87E-4 | -0.07 | -0.025 |
| **sex \* trial** | -0.0302±0.016 | -1.901 | 0.29 | -0.061 | 0.001 |

**Table S6:** 2-sided GEE regression of GCaMP response to object (Figure 1F): *Z-scored ΔF/F = trial + sex + sex*trial + intercept, grouped by cell*. Number of cells (groups) = 411, minimum samples per group 199, maximum samples per group 212, dependence structure = independence, family = Gaussian. P-values Bonferroni corrected for 5 comparisons.

| *Model: Z-scored $\Delta F/F$ responding to object = trial + sex + trial\*sex + intercept, grouped by cell* | $\beta \pm$ standard error | z-stat | p-value | 95% CI [lower, upper] | |
| --- | --- | --- | --- | --- | --- |
| **Intercept** | -0.102±1.698 | -3.042 | 0.012 | -0.167 | -0.036 |
| **sex** | 0.127±0.052 | 2.452 | 0.0711 | 0.025 | 0.229 |
| **trial** | -0.0113±0.005 | -2.482 | 0.065 | -0.02 | -0.002 |
| **sex \* trial** | -0.0108±0.008 | -1.38 | 0.837 | -0.026 | 0.005 |

**Table S7:** 2-sided GEE regression of GCaMP response to male contact (Figure S2C): *Z-scored ΔF/F = bout + sex + sex*trial + intercept, grouped by cell*. Number of cells (groups) = 409, minimum samples per group 199, maximum samples per group 210, dependence structure = independence, family = Gaussian. P-values Bonferroni corrected for 5 *+ sex*trial + intercept, grouped by cell*. Number of cells (groups) = 409, minimum samples per group 199, maximum samples per group 210, dependence structure = independence, family = Gaussian. P-values Bonferroni corrected for 5 comparisons.

| *Model: Z-scored $\Delta F/F$ responding to male contact = bout + sex + trial\*sex + intercept, grouped by cell* | $\beta \pm$ standard error | z-stat | p-value | 95% CI [lower, upper] | |
| --- | --- | --- | --- | --- | --- |
| **Intercept** | 0.134±0.030 | 4.447 | 4.36E-5 | 0.075 | 0.193 |
| **sex** | -0.298±0.05 | -6.494 | 4.17E-10 | -0.387 | -0.208 |
| **trial** | -0.004±0.004 | -0.789 | 0.43 | -0.012 | 0.005 |
| **sex \* trial** | 0.016±0.007 | 2.332 | 0.02 | 0.003 | 0.03 |

**Table S8:** 2-sided GEE regression of GCaMP response to estrus female contact (Figure S2C): *Z-scored ΔF/F = bout + sex + sex*trial + intercept, grouped by cell*. Number of cells (groups) = 409, minimum samples per group 199, maximum samples per group 210, dependence structure = independence, family = Gaussian. P-values Bonferroni corrected for 5 comparisons.

| *Model: Z-scored $\Delta F/F$ responding to estrus female contact = bout + sex + trial\*sex + intercept, grouped by cell* | $\beta \pm$ standard error | z-stat | p-value | 95% CI [lower, upper] | |
| --- | --- | --- | --- | --- | --- |
| **Intercept** | -0.005±0.028 | -0.189 | 0.85 | -0.059 | 0.049 |
| **sex** | 0.154±0.047 | 3.26 | 0.0056 | 0.061 | 0.246 |
| **trial** | 0.002±0.004 | 0.46 | 0.65 | -0.006 | 0.009 |
| **sex \* trial** | -0.021±0.007 | -3.14 | 0.008 | -0.034 | -0.008 |

**Table S9:** 2-sided GEE regression of GCaMP response to non-estrus female contact (Figure S2C): *Z-scored ΔF/F = bout + sex + sex*trial + intercept, grouped by cell*. Number of cells (groups) = 409, minimum samples per group 199, maximum samples per group 210, dependence structure = independence, family = Gaussian. P-values Bonferroni corrected for 5 comparisons.

| *Model: Z-scored $\Delta F/F$ responding to non-estrus female contact = bout + sex + trial\*sex + intercept, grouped by cell* | $\beta \pm$ standard error | z-stat | p-value | 95% CI [lower, upper] | |
| --- | --- | --- | --- | --- | --- |
| **Intercept** | 0.039±0.023 | 1.73 | 0.42 | -0.005 | 0.084 |
| **sex** | 0.030±0.044 | 0.66 | 0.51 | -0.058 | 0.117 |
| **trial** | 0.006±0.003 | 1.803 | 0.357 | -0.001 | 0.013 |
| **sex \* trial** | -0.0087±0.006 | -1.513 | 0.13 | -0.02 | 0.003 |

**Table S10:** Statistics for Figures S2.

| Figure | Description | Sample size | Statistical test | Statistical values | Test Statistic | p Value | Significance |
| --- | --- | --- | --- | --- | --- | --- | --- |
| Fig. S2D | Difference in same vs opposite sex contact decoding accuracy before versus after opposite sex experience | 13 mice | 2-sided, 2-sample relative t-Test | Difference in means | t=2.23 | p=0.0459 | * |
| Fig. S2E | Difference in distance between imaged and stimulus mouse before vs after opposite sex experience | 16, 19 mice | 2-sided, 2 sample, t-Test | Difference in means | t=0.163 | p=0.871 | NS |

**Table S11:** Statistics for Figure 2.

| Figure | Description | Sample size | Statistical test | Statistical values | Test Statistic | P-value | Significance |
| --- | --- | --- | --- | --- | --- | --- | --- |
| Fig. 2C | Overlap in neurons responsive to male and food | 411 neurons | Each point in the null distribution is constructed by shuffling whether a neuron responds to each stimulus type, and then calculating the percentage of cells responsive to both stimulus types from the whole population. | P-value is the fraction of simulated values from the null distribution greater than the true percentage of cells responsive to both stimuli | NA | 0 | *** |
| Fig. 2D | Overlap in neurons responsive to estrus female and food | 411 neurons | Each point in the null distribution is constructed by shuffling whether a neuron responds to each stimulus type, and then calculating the percentage of cells responsive to both stimulus types from the whole population. | P-value is the fraction of simulated values from the null distribution greater than the true percentage of cells responsive to both stimuli | NA | 0 | *** |
| Fig. 2E | Relationship between response to food arrival and response to male | 411 neurons | Pearson's correlation | Linear relationship | r=0.295 | p=1.1 E-9 | *** |
| Fig. 2F | Relationship between response to food arrival and response to estrus female | 411 neurons | Pearson's correlation | Linear relationship | r=0.237 | p=1.1 5E-6 | *** |

**Table S12:** Statistics for Figure S3.

| Figure | Description | Sample size | Statistical test | Statistical values | Test Statistic | p Value | Significance |
| --- | --- | --- | --- | --- | --- | --- | --- |
| Fig. S3A | Relationship between response to food arrival and object-corrected response to male | 411 neurons | Pearson's correlation | Linear relationship | r=0.23 | p=2.5 5E-6 | *** |
| Fig. S3B | Relationship between response to food arrival and object-corrected response to estrus female | 411 neurons | Pearson's correlation | Linear relationship | r=0.173 | p=4.2 E-4 | *** |

**Table S13:**
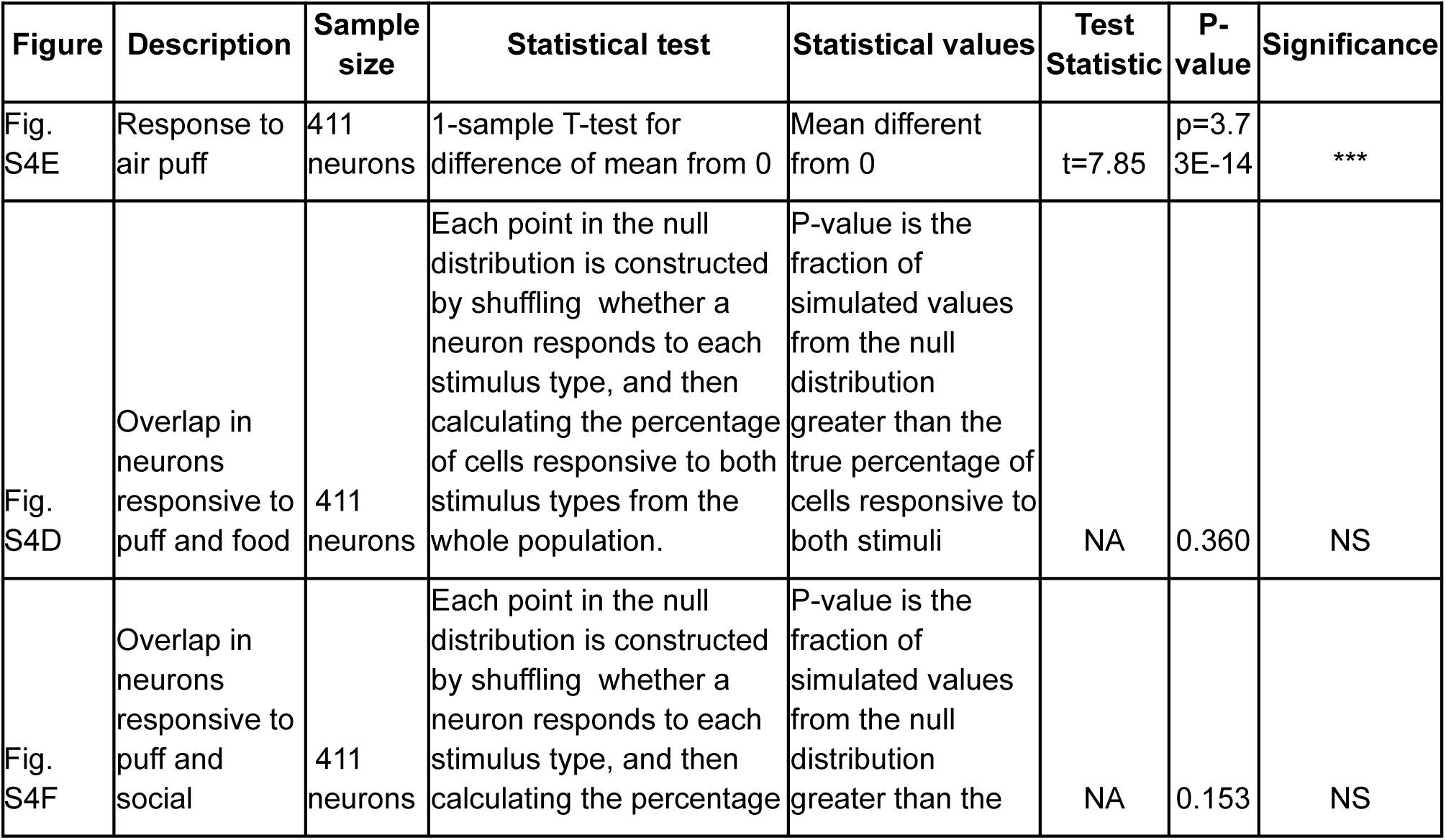
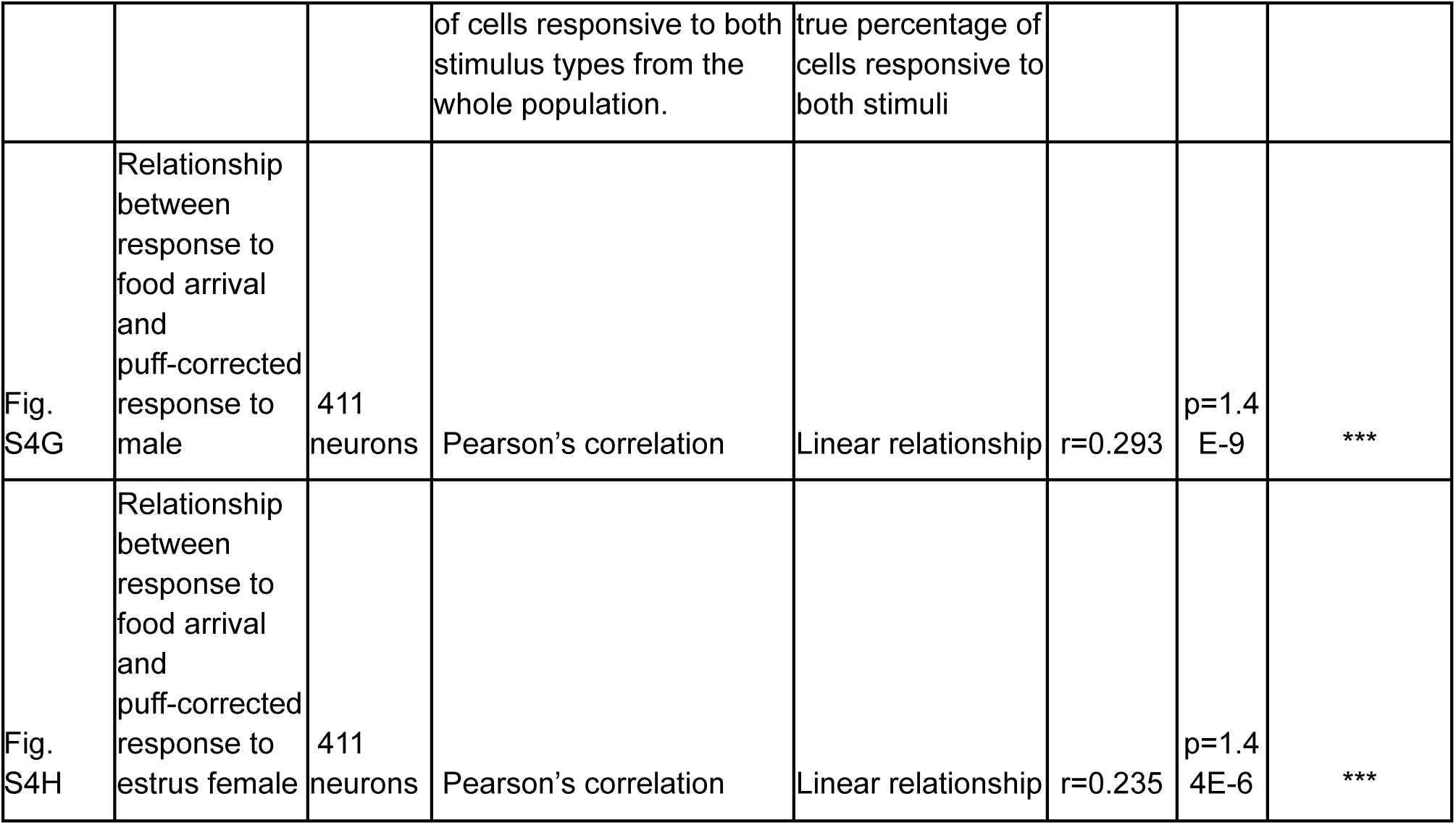
Statistics for Figure S4.

**Table S14:** Statistics for Figure S5.

| Figure | Description | Sample size | Statistical test | Statistical values | Test Statistic | P-value | Significance |
| --- | --- | --- | --- | --- | --- | --- | --- |
| Fig. S5B | Relationship between response to food arrival and pupil-corrected response to male | 306 neurons | Pearson's correlation | Linear relationship | r=0.277 | p=8.9 6E-0.7 | *** |
| Fig. S5F | Difference in pupil diameter while sated versus hungry | 20, 18 mice | 2-sided, 2 sample, t-Test | Difference in means | t=-0.33 | p=0.7 43 | NS |
| Fig. S5G | Difference in limb speed while sated versus hungry | 20, 18 mice | 2-sided, 2 sample, t-Test | Difference in means | t=1.58 | p=0.1 24 | NS |
| Fig. S5H | Difference in pupil diameter before versus after opposite sex experience | 16, 20 mice | 2-sided, 2 sample, t-Test | Difference in means | t=0.261 | p=0.7 96 | NS |
| Fig. S5I | Difference in limb speed before versus after opposite sex experience | 16, 20 mice | 2-sided, 2 sample, t-Test | Difference in means | t=0.487 | p=0.6 29 | NS |

**Table S15:** Statistics for Figure S6.

| Figure | Description | Sample size | Statistical test | Statistical values | Test Statistic | p Value | Significance |
| --- | --- | --- | --- | --- | --- | --- | --- |
| Fig. S6A | Difference in proportion of neurons responsive to food, when food arrives first or male stimulus arrives first | 118 neurons for food first, 293 neurons for social first | 2-sided, 2-sample test of proportions | Difference in proportion | Z=10.13 | p=3.86 E-24 | *** |
| Fig. S6A | Difference in proportion of neurons responsive to food, when food arrives first or male stimulus arrives first | 118 neurons for food first, 293 neurons for social first | 2-sided, 2-sample test of proportions | Difference in proportion | Z=6.10 | p=1.06 E-9 | *** |
| Fig. S6B | Overlap in neurons responsive to male and food when food trial occurs before male trial | 411 neurons | Each point in the null distribution is constructed by shuffling whether a neuron responds to each stimulus type, and then calculating the percentage of cells responsive to both stimulus types from the whole population. | P-value is the fraction of simulated values from the null distribution greater than the true percentage of cells responsive to both stimuli | NA | 0.0243 | *** |
| Fig. S6B | Overlap in neurons responsive to male and food when food trial occurs before male trial | 411 neurons | Each point in the null distribution is constructed by shuffling whether a neuron responds to each stimulus type, and then calculating the percentage of cells responsive to both stimulus types from the whole population. | P-value is the fraction of simulated values from the null distribution greater than the true percentage of cells responsive to both stimuli | NA | 0.0009 | *** |

**Table S16:** Statistics for Figure S7.

| Figure | Description | Sample size | Statistical test | Statistical values | Test Statistic | p Value | Significance |
| --- | --- | --- | --- | --- | --- | --- | --- |
| Fig. S7B | Relationship between AP location and response to male stimulus | 435 neurons | Pearson's correlation | Linear relationship | r=0.032 | p=0.51 | NS |
| Fig. S7B | Relationship between ML location and response to male stimulus | 435 neurons | Pearson's correlation | Linear relationship | r=0.045 | p=0.34 | NS |
| Fig. S7B | Relationship between DV location and response to male stimulus | 19 fields | Pearson's correlation | Linear relationship | r=0.030 | p=0.90 | NS |
| Fig. S7B | Relationship between AP location and response to estrus female stimulus | 435 neurons | Pearson's correlation | Linear relationship | r=-0.084 | p=0.080 | NS |
| Fig. S7B | Relationship between ML location and response to estrus female stimulus | 435 neurons | Pearson's correlation | Linear relationship | r=0.037 | p=0.44 | NS |
| Fig. S7B | Relationship between DV location and response to estrus female stimulus | 19 fields | Pearson's correlation | Linear relationship | r=0.028 | p=0.91 | NS |
| Fig. S7B | Relationship between AP location and response to food | 435 neurons | Pearson's correlation | Linear relationship | r=-0.07 | p=0.14 | NS |
| Fig. S7B | Relationship between ML location and response to food | 435 neurons | Pearson's correlation | Linear relationship | r=0.0016 | p=0.97 | NS |
| Fig. S7B | Relationship between DV location and response to food | 19 fields | Pearson's correlation | Linear relationship | r=-0.41 | p=0.080 | NS |

**Table S17:** Statistics for Figure 3.

| Figure | Description | Sample size | Statistical test | Statistical values | Test Statistic | p Value | Significance |
| --- | --- | --- | --- | --- | --- | --- | --- |
| Fig. 3C | Average lick rate during food presentation when mice were sated versus hungry | 13 mice | 2-sided, 2-sample relative t-Test | Difference in means | t=6.697 | p=2.21E-5 | *** |
| Fig. 3D | Change in magnitude in response to food on hungry-sated sessions | 137 neurons | 2-sided, 1-sample t-Test | Difference in mean from 0 | t=13.18 | p=4.5E-26 | *** |
| Fig. 3F | Proportion of neurons responsive to food but not social | 427 neurons, 394 neurons | 2-sided, 2-sample test of proportions | Difference in proportion | Z=9.1 | p=8.89E-20 | *** |
| Fig. 3F | Proportion of neurons responsive to social but not food | 427 neurons, 394 neurons | 2-sided, 2-sample test of proportions | Difference in proportion | Z=-3.14 | p=0.0017 | ** |
| Fig. 3F | Proportion of neurons responsive to food and social | 427 neurons, 394 neurons | 2-sided, 2-sample test of proportions | Difference in proportion | Z=7.197 | p=6.18E-13 | *** |
| Fig. 3I | Fraction of males for which more syllables were detected during female stimulus trials after than before opposite sex experience | 5 male mice | bootstrap | Greater than chance | - | p=0.031 | * |
| Fig. 3J | Change in magnitude in response to opposite-sex after-before opposite-sex experience, in food responsive-neurons | 95 neurons | 2-sided, 1-sample t-Test | Difference in mean from 0 | t=2.25 | p=0.0268 | * |
| Fig. 3L | Proportion of neurons responsive to food but not opposite-sex | 321 neurons, 306 neurons | 2-sided, 2-sample test of proportions | Difference in proportion | Z=-1.30 | p=0.1922 | NS |
| Fig. 3L | Proportion of neurons responsive to opposite-sex but not food | 321 neurons, 306 neurons | 2-sided, 2-sample test of proportions | Difference in proportion | Z=1.54 | p=0.12 | NS |
| Fig. 3L | Proportion of neurons responsive to food and opposite-sex | 321 neurons, 306 neurons | 2-sided, 2-sample test of proportions | Difference in proportion | Z=2.24 | p=0.0253 | * |
| Fig. 3L | Proportion of neurons responsive to same-sex but not food | 321 neurons, 306 neurons | 2-sided, 2-sample test of proportions | Difference in proportion | Z=-0.020 | p=0.984 | NS |
| Fig. 3L | Proportion of neurons responsive to food and same-sex | 321 neurons, 306 neurons | 2-sided, 2-sample test of proportions | Difference in proportion | Z=0.945 | p=0.343 | NS |

**Table S18:** Statistics for Figure S8.

| Figure | Description | Sample size | Statistical test | Statistical values | Test Statistic | p Value | Significance |
| --- | --- | --- | --- | --- | --- | --- | --- |
| Fig. S8B | Magnitude in response to food on first versus second sated session | 137 neurons | Pearson's correlation | Linear relationship | R=0.57 | p=3.2E-13 | *** |
| Fig. S8B | Change in magnitude in response to food second-first sated session | 137 neurons | 2-sided, 1-sample t-Test | Difference in mean from 0 | t=0.743 | p=0.46 | NS |
| Fig. S8C | Magnitude in response to food on first versus second hungry session | 137 neurons | Pearson's correlation | Linear relationship | R=0.48 | p=6.10E-20 | *** |
| Fig. S8C | Change in magnitude in response to food second-first hungry session | 137 neurons | 2-sided, 1-sample t-Test | Difference in mean from 0 | t=0.152 | p=0.879 | NS |
| Fig. S8D | Change in magnitude in response to food from sated to hungry days on week 1 versus week 2 | 137 neurons | Pearson's correlation | Linear relationship | R=0.44 | p=1.02 E-7 | *** |

**Table S19:** Statistics for Figure S9 (✝ with Bonferroni correction for 3 comparisons)

| Figure | Description | Sample size | Statistical test | Statistical values | Test Statistic | p Value † | Significance |
| --- | --- | --- | --- | --- | --- | --- | --- |
| Fig. S9 | Relationship between change in food response when hungry vs sated and AP location | 140 neurons | Pearson's correlation | Linear relationship | R=-0.01 | p=2.8 | NS |
| Fig. S9 | Relationship between change in food response when hungry vs sated and ML location | 140 neurons | Pearson's correlation | Linear relationship | R=-0.11 | p=0.61 | NS |
| Fig. S9 | Relationship between change in food response when hungry vs sated and DV location | 12 fields | Pearson's correlation | Linear relationship | R=-0.75 | p=0.02 | * |

**Table S20:**
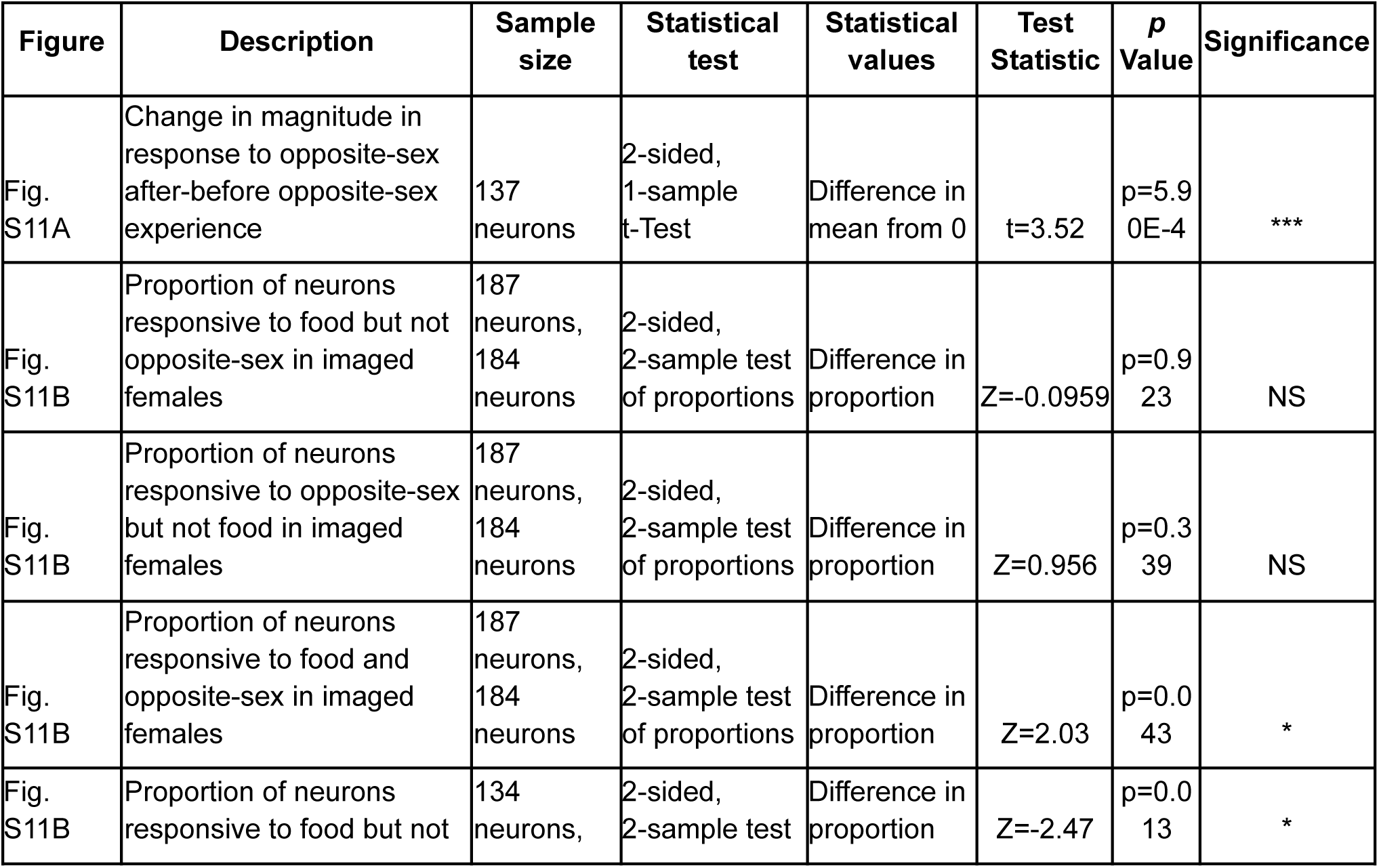
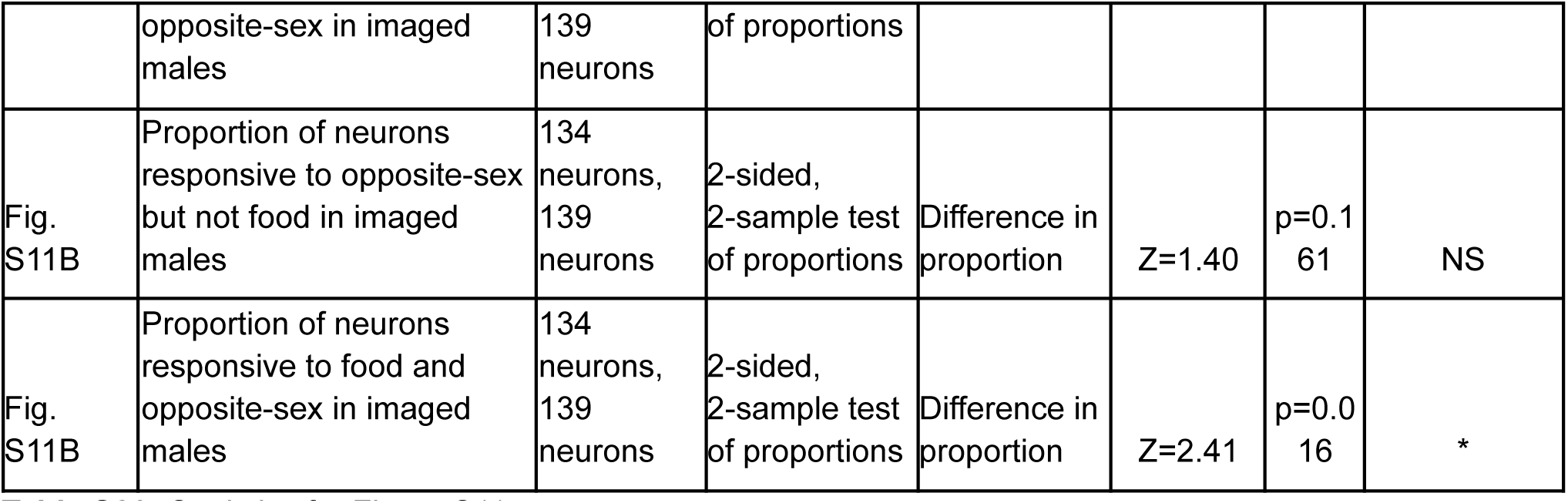
Statistics for Figure S11.

**Table S21:** Statistics for Figure S12.

| Figure | Description | Sample size | Statistical test | Statistical values | Test Statistic | p Value † | Significance |
| --- | --- | --- | --- | --- | --- | --- | --- |
| Fig. S12B | Magnitude in response to same-sex before vs after opposite-sex experience | 137 neurons | Pearson's correlation | Linear relationship | R=0.007 | p=0.939 | NS |
| Fig. S12B | Change in magnitude in response to same-sex after-before opposite-sex experience | 137 neurons | 2-sided, 1-sample t-Test | Difference in mean from 0 | t=-0.23 | p=0.819 | NS |

**Table S22:** Statistics for Figure S13. (✝ with Bonferroni correction for 3 tests)

| Figure | Description | Sample size | Statistical test | Statistical values | Test Statistic | p Value † | Significance |
| --- | --- | --- | --- | --- | --- | --- | --- |
| Fig. S13 | Relationship between change in opposite-sex stimulus response after vs before opposite-sex experience and AP location | 137 neurons | Pearson's correlation | Linear relationship | R=-0.385 | p=1.03E-5 | *** |
| Fig. S13 | Relationship between change in opposite-sex stimulus response after vs before opposite-sex experience and ML location | 137 neurons | Pearson's correlation | Linear relationship | R=-0.214 | p=0.036 | * |
| Fig. S13 | Relationship between change in opposite-sex stimulus response after vs before opposite-sex experience and DV location | 12 fields | Pearson's correlation | Linear relationship | R=-0.115 | p=2.16 | NS |

**Table S23:** Statistics for Figure S14.

| Figure | Description | Sample size | Statistical test | Statistical values | Test Statistic | p Value | Significance |
| --- | --- | --- | --- | --- | --- | --- | --- |
| Fig. S14B | Proportion of neurons responsive to food but not opposite-sex | 244 neurons, 408 neurons | 2-sided, 2-sample test of proportions | Difference in proportion | Z=3.851 | p=1.18E-4 | *** |
| Fig. S14B | Proportion of neurons responsive to opposite-sex but not food | 244 neurons, 408 neurons | 2-sided, 2-sample test of proportions | Difference in proportion | Z=-4.14 | p=3.48E-5 | *** |
| Fig. S14B | Proportion of neurons responsive to food and opposite-sex | 244 neurons, 408 neurons | 2-sided, 2-sample test of proportions | Difference in proportion | Z=0.071 | p=0.943 | NS |
| Fig. S14B | Proportion of neurons responsive to same-sex but not food | 244 neurons, 408 neurons | 2-sided, 2-sample test of proportions | Difference in proportion | Z=0.900 | p=0.368 | NS |
| Fig. S14B | Proportion of neurons responsive to food and same-sex | 244 neurons, 408 neurons | 2-sided, 2-sample test of proportions | Difference in proportion | Z=1.309 | p=0.191 | NS |
| Fig. S14C | Lick rate during first food presentation before vs after habituation | 11 mice | 2-sided, 2-sample relative t-Test | Difference in means | t=2.273 | p=0.0463 | * |
| Fig. S14E | Magnitude in response to opposite-sex before vs after habituation | 165 neurons | Pearson's correlation | Linear relationship | R=0.37 | p=8.45E-7 | *** |
| Fig. S14E | Change in magnitude in response to opposite-sex after-before habituation | 165 neurons | 2-sided, 1-sample t-Test | Difference in mean from 0 | t=-4.96 | p=1.73E-6 | *** |

**Table S24:** Differentially expressed genes (DEGs) in nuclei from sated versus hungry animals (Figure 4B)

| Gene | Celltype | Log2FC | P-value | Adjusted p-value | Direction of change |
| --- | --- | --- | --- | --- | --- |
| 1700042O10Rik | dopamine | -0.956 | 5.12E-53 | 1.09E-48 | hungry |
| 4930419G24Rik | dopamine | -0.809 | 4.97E-41 | 1.06E-36 | hungry |
| Col25a1 | dopamine | -0.705 | 1.14E-36 | 2.43E-32 | hungry |
| Gm26905 | dopamine | -0.616 | 1.16E-18 | 2.48E-14 | hungry |
| Gm37459 | dopamine | -0.595 | 1.34E-23 | 2.86E-19 | hungry |
| 4921534H16Rik | dopamine | -0.585 | 1.70E-45 | 3.64E-41 | hungry |
| Kcnd3 | dopamine | -0.582 | 7.35E-35 | 1.57E-30 | hungry |
| Gm26936 | dopamine | -0.560 | 2.50E-44 | 5.35E-40 | hungry |
| Fryl | dopamine | -0.534 | 1.05E-37 | 2.24E-33 | hungry |
| Ddc | dopamine | -0.532 | 9.22E-25 | 1.97E-20 | hungry |
| Erc2 | dopamine | -0.529 | 8.19E-14 | 1.75E-09 | hungry |
| Nwd2 | dopamine | -0.514 | 4.10E-26 | 8.77E-22 | hungry |
| Prkca | dopamine | -0.508 | 2.82E-19 | 6.03E-15 | hungry |
| Kcnj6 | dopamine | -0.500 | 6.17E-21 | 1.32E-16 | hungry |
| Gm15398 | dopamine | -0.497 | 2.46E-10 | 5.27E-06 | hungry |
| Gm15261 | dopamine | -0.488 | 6.69E-19 | 1.43E-14 | hungry |
| Fam184a | dopamine | -0.463 | 1.69E-26 | 3.62E-22 | hungry |
| Hdac9 | dopamine | -0.449 | 3.68E-15 | 7.86E-11 | hungry |
| Clstn2 | dopamine | -0.447 | 1.18E-10 | 2.52E-06 | hungry |
| D030068K23Rik | dopamine | -0.441 | 6.24E-18 | 1.33E-13 | hungry |
| 9530059O14Rik | dopamine | -0.437 | 3.51E-24 | 7.51E-20 | hungry |
| Cntnap4 | dopamine | -0.429 | 9.89E-18 | 2.11E-13 | hungry |
| Robo2 | dopamine | -0.427 | 9.68E-11 | 2.07E-06 | hungry |
| Cpne7 | dopamine | -0.422 | 5.88E-20 | 1.26E-15 | hungry |
| C79798 | dopamine | -0.420 | 1.25E-22 | 2.67E-18 | hungry |
| AC165271.1 | dopamine | -0.418 | 5.11E-18 | 1.09E-13 | hungry |
| Rreb1 | dopamine | -0.412 | 5.04E-12 | 1.08E-07 | hungry |
| Apba1 | dopamine | -0.402 | 5.11E-12 | 1.09E-07 | hungry |
| Gm47271 | dopamine | -0.399 | 2.12E-17 | 4.53E-13 | hungry |
| Sox6 | dopamine | -0.391 | 5.89E-07 | 0.01258638 | hungry |
| Cdh11 | dopamine | -0.390 | 5.70E-13 | 1.22E-08 | hungry |
| Atp5k | dopamine | 0.381 | 4.32E-74 | 9.24E-70 | sated |
| Ppia | dopamine | 0.384 | 5.20E-83 | 1.11E-78 | sated |
| Tmsb4x | dopamine | 0.394 | 9.73E-97 | 2.08E-92 | sated |
| Tpt1 | dopamine | 0.395 | 2.31E-78 | 4.93E-74 | sated |
| Atp1b1 | dopamine | 0.399 | 1.15E-84 | 2.46E-80 | sated |
| Kctd8 | dopamine | 0.405 | 8.21E-39 | 1.76E-34 | sated |
| Aldoa | dopamine | 0.409 | 8.53E-89 | 1.82E-84 | sated |
| Ndufa4 | dopamine | 0.409 | 1.76E-68 | 3.77E-64 | sated |
| Zfp804b | dopamine | 0.409 | 2.23E-16 | 4.77E-12 | sated |
| Dgkb | dopamine | 0.428 | 4.90E-48 | 1.05E-43 | sated |
| Ckb | dopamine | 0.436 | 3.02E-85 | 6.46E-81 | sated |
| Grid2 | dopamine | 0.439 | 5.49E-23 | 1.17E-18 | sated |
| Cst3 | dopamine | 0.440 | 6.40E-88 | 1.37E-83 | sated |
| Chchd2 | dopamine | 0.450 | 4.28E-92 | 9.15E-88 | sated |
| Plp1 | dopamine | 0.471 | 1.81E-95 | 3.88E-91 | sated |
| Cox8a | dopamine | 0.492 | 1.09E-134 | 2.33E-130 | sated |
| Cox4i1 | dopamine | 0.514 | 1.65E-135 | 3.53E-131 | sated |
| Cox6c | dopamine | 0.515 | 4.01E-127 | 8.58E-123 | sated |
| Hspa8 | dopamine | 0.519 | 6.20E-129 | 1.33E-124 | sated |
| Hsp90ab1 | dopamine | 0.528 | 4.98E-153 | 1.06E-148 | sated |
| Rbfox1 | dopamine | 0.532 | 3.64E-13 | 7.79E-09 | sated |
| Pcdh9 | dopamine | 0.545 | 2.55E-62 | 5.44E-58 | sated |
| Calm1 | dopamine | 0.559 | 7.92E-142 | 1.69E-137 | sated |
| Hsp90aa1 | dopamine | 0.584 | 2.64E-145 | 5.65E-141 | sated |
| Apoe | dopamine | 0.600 | 5.12E-121 | 1.10E-116 | sated |
| Ubb | dopamine | 0.629 | 7.21E-125 | 1.54E-120 | sated |
| AC149090.1 | dopamine | 0.782 | 3.46E-273 | 7.39E-269 | sated |
| Gm42418 | dopamine | 0.785 | 5.94E-225 | 1.27E-220 | sated |
| Bc1 | dopamine | 0.847 | 0 | 0 | sated |
| Fth1 | dopamine | 1.005 | 0 | 0 | sated |
| Gm38505 | gaba | -0.548 | 1.80E-15 | 3.85E-11 | hungry |
| Npas3 | gaba | -0.540 | 1.06E-27 | 2.27E-23 | hungry |
| D030068K23Rik | gaba | -0.526 | 4.97E-37 | 1.06E-32 | hungry |
| Gm26883 | gaba | -0.501 | 1.41E-25 | 3.01E-21 | hungry |
| Mef2c | gaba | -0.444 | 2.56E-30 | 5.47E-26 | hungry |
| Dab1 | gaba | -0.425 | 2.11E-29 | 4.50E-25 | hungry |
| AY036118 | gaba | -0.385 | 1.29E-51 | 2.75E-47 | hungry |
| Dgkb | gaba | 0.390 | 7.21E-11 | 1.54E-06 | sated |
| Pdzd2 | gaba | 0.400 | 3.56E-08 | 0.000761786 | sated |
| Hsp90ab1 | gaba | 0.407 | 1.65E-25 | 3.54E-21 | sated |
| Pcsk2 | gaba | 0.413 | 9.65E-14 | 2.06E-09 | sated |
| Hsp90aa1 | gaba | 0.418 | 1.05E-23 | 2.25E-19 | sated |
| Cox6c | gaba | 0.418 | 2.38E-27 | 5.09E-23 | sated |
| Cox8a | gaba | 0.431 | 8.15E-32 | 1.74E-27 | sated |
| Cox4i1 | gaba | 0.435 | 1.01E-29 | 2.16E-25 | sated |
| Plp1 | gaba | 0.440 | 3.21E-27 | 6.87E-23 | sated |
| Ubb | gaba | 0.442 | 1.33E-19 | 2.84E-15 | sated |
| Tsix | gaba | 0.445 | 8.39E-33 | 1.79E-28 | sated |
| Kctd8 | gaba | 0.448 | 3.15E-15 | 6.73E-11 | sated |
| Calm1 | gaba | 0.453 | 3.62E-30 | 7.74E-26 | sated |
| Zfp804b | gaba | 0.587 | 1.76E-07 | 0.003772174 | sated |
| AC149090.1 | gaba | 0.613 | 1.47E-45 | 3.15E-41 | sated |
| ErbB4 | gaba | 0.635 | 1.98E-13 | 4.23E-09 | sated |
| Pcdh9 | gaba | 0.688 | 4.91E-24 | 1.05E-19 | sated |
| Ntng1 | gaba | 0.732 | 1.21E-24 | 2.59E-20 | sated |
| Fth1 | gaba | 0.851 | 1.14E-116 | 2.43E-112 | sated |
| Bc1 | gaba | 0.961 | 8.15E-148 | 1.74E-143 | sated |
| Robo2 | glut | -0.392 | 2.78E-22 | 5.95E-18 | hungry |
| Hs3st4 | glut | -0.423 | 1.06E-13 | 2.26E-09 | hungry |
| Chrm3 | glut | -0.396 | 1.43E-12 | 3.06E-08 | hungry |
| Fth1 | glut | 1.030 | 2.14E-83 | 4.57E-79 | sated |
| AC149090.1 | glut | 0.853 | 1.02E-52 | 2.18E-48 | sated |
| Bc1 | glut | 0.791 | 6.24E-47 | 1.33E-42 | sated |
| Calm1 | glut | 0.630 | 1.59E-34 | 3.39E-30 | sated |
| Hsp90aa1 | glut | 0.636 | 6.92E-31 | 1.48E-26 | sated |
| Gm42418 | glut | 0.745 | 5.19E-30 | 1.11E-25 | sated |
| Ubb | glut | 0.655 | 4.59E-29 | 9.82E-25 | sated |
| Cox4i1 | glut | 0.549 | 1.16E-26 | 2.48E-22 | sated |
| Hsp90ab1 | glut | 0.556 | 6.37E-26 | 1.36E-21 | sated |
| Hspa8 | glut | 0.544 | 5.32E-23 | 1.14E-18 | sated |
| Cox8a | glut | 0.486 | 2.91E-22 | 6.23E-18 | sated |
| Cox6c | glut | 0.500 | 1.30E-21 | 2.79E-17 | sated |
| Plp1 | glut | 0.543 | 6.77E-21 | 1.45E-16 | sated |
| Apoe | glut | 0.566 | 7.62E-21 | 1.63E-16 | sated |
| Chchd2 | glut | 0.512 | 9.69E-21 | 2.07E-16 | sated |
| Cst3 | glut | 0.497 | 1.69E-19 | 3.62E-15 | sated |
| Aldoa | glut | 0.432 | 2.28E-16 | 4.87E-12 | sated |
| Ppia | glut | 0.447 | 9.12E-16 | 1.95E-11 | sated |
| Tpt1 | glut | 0.446 | 1.58E-15 | 3.39E-11 | sated |
| Stmn3 | glut | 0.400 | 5.58E-14 | 1.19E-09 | sated |
| Ckb | glut | 0.413 | 1.84E-13 | 3.93E-09 | sated |
| Slc25a3 | glut | 0.400 | 4.61E-13 | 9.87E-09 | sated |
| Uqcrh | glut | 0.396 | 6.17E-13 | 1.32E-08 | sated |
| Luzp2 | glut | 0.474 | 1.09E-12 | 2.32E-08 | sated |
| Ndufa4 | glut | 0.433 | 1.91E-12 | 4.09E-08 | sated |
| Tecr | glut | 0.401 | 2.89E-12 | 6.18E-08 | sated |
| Chchd10 | glut | 0.385 | 1.09E-11 | 2.32E-07 | sated |
| Kctd8 | glut | 0.409 | 5.23E-10 | 1.12E-05 | sated |
| Ptgds | glut | 0.390 | 1.41E-09 | 3.02E-05 | sated |
| Cox5a | glut | 0.392 | 2.63E-08 | 0.000562871 | sated |
| Ebf1 | glut | 0.460 | 1.32E-06 | 0.028204384 | sated |
| Gulp1 | glut | 0.446 | 1.76E-06 | 0.037545199 | sated |

**Table S25:**
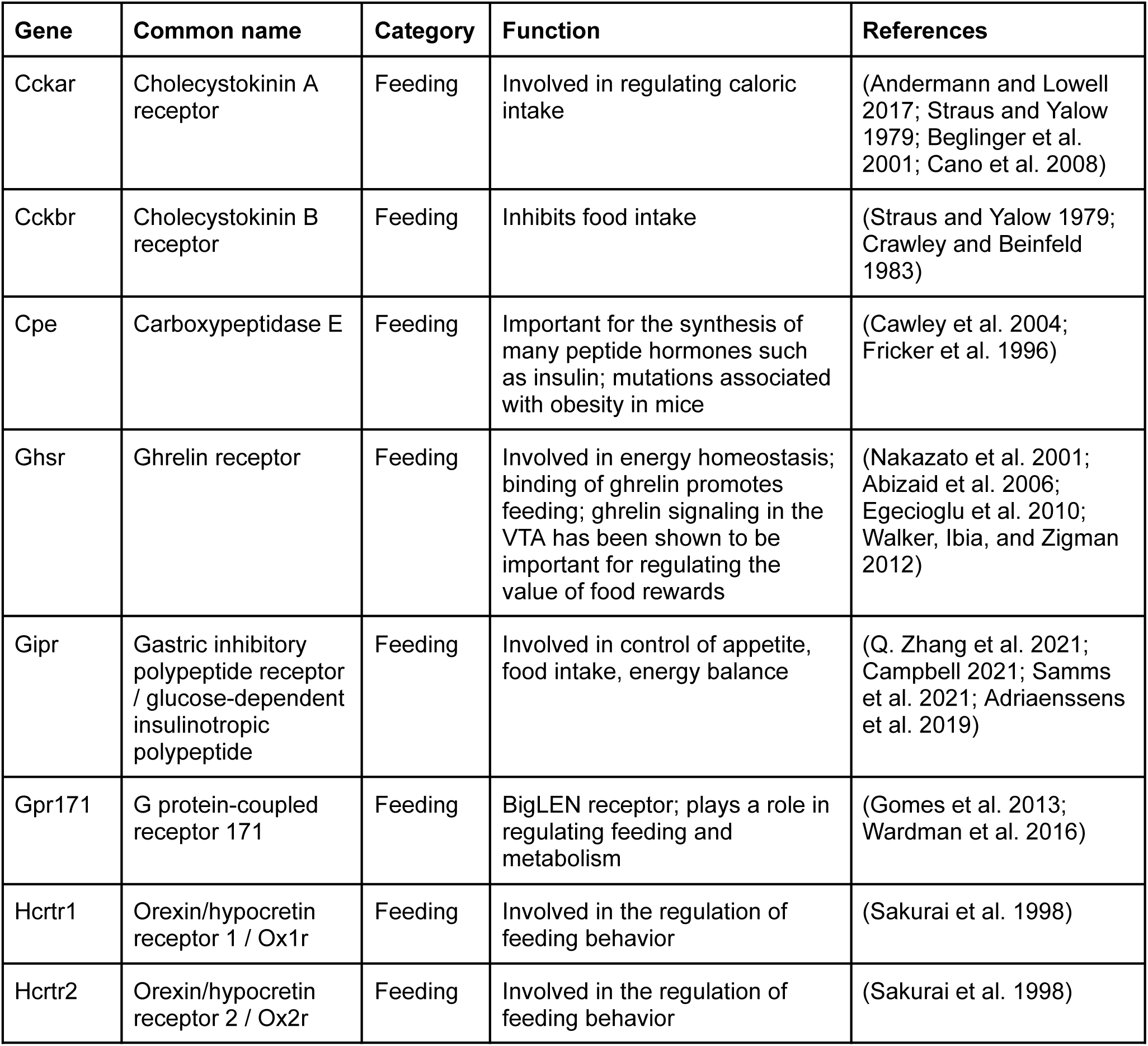
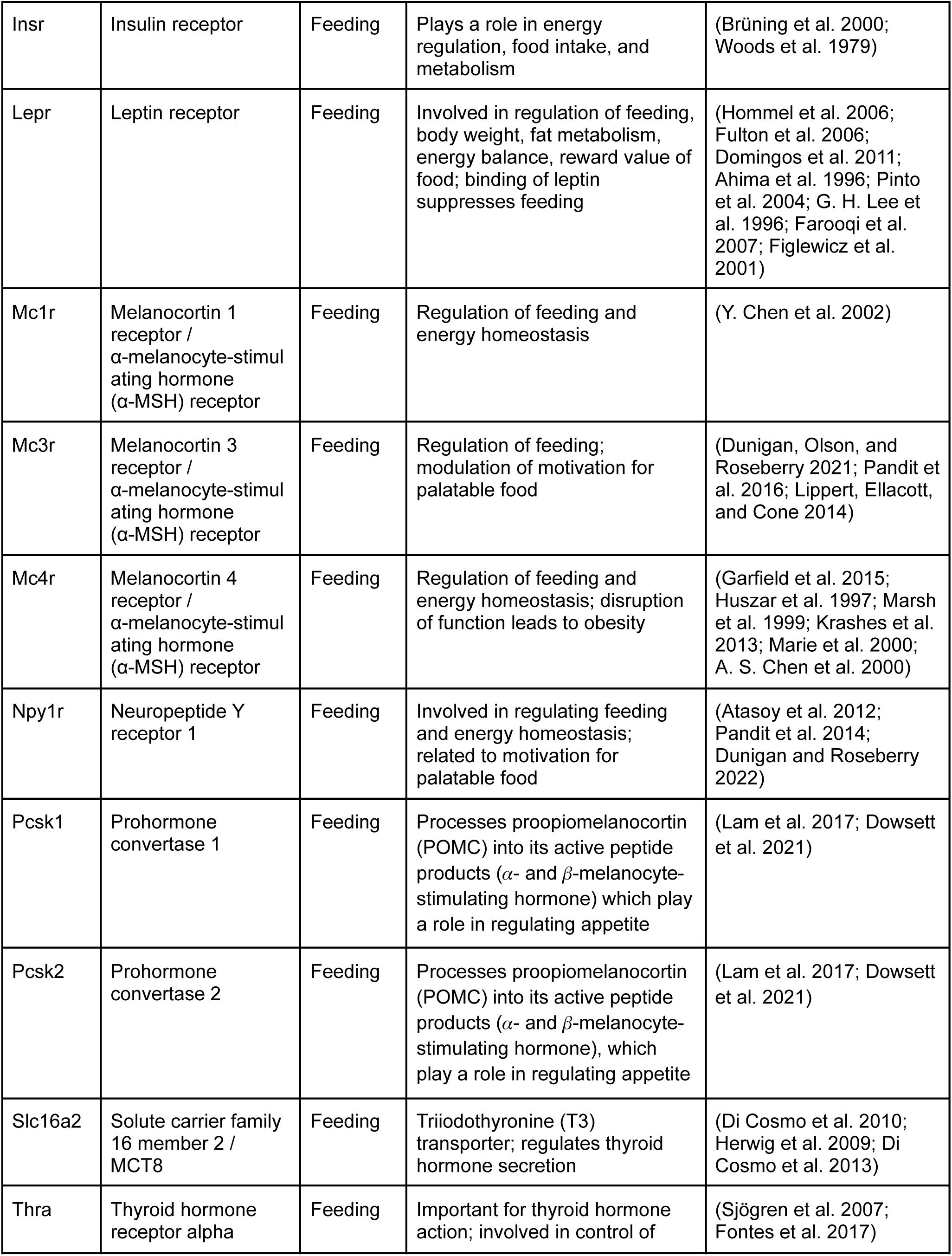
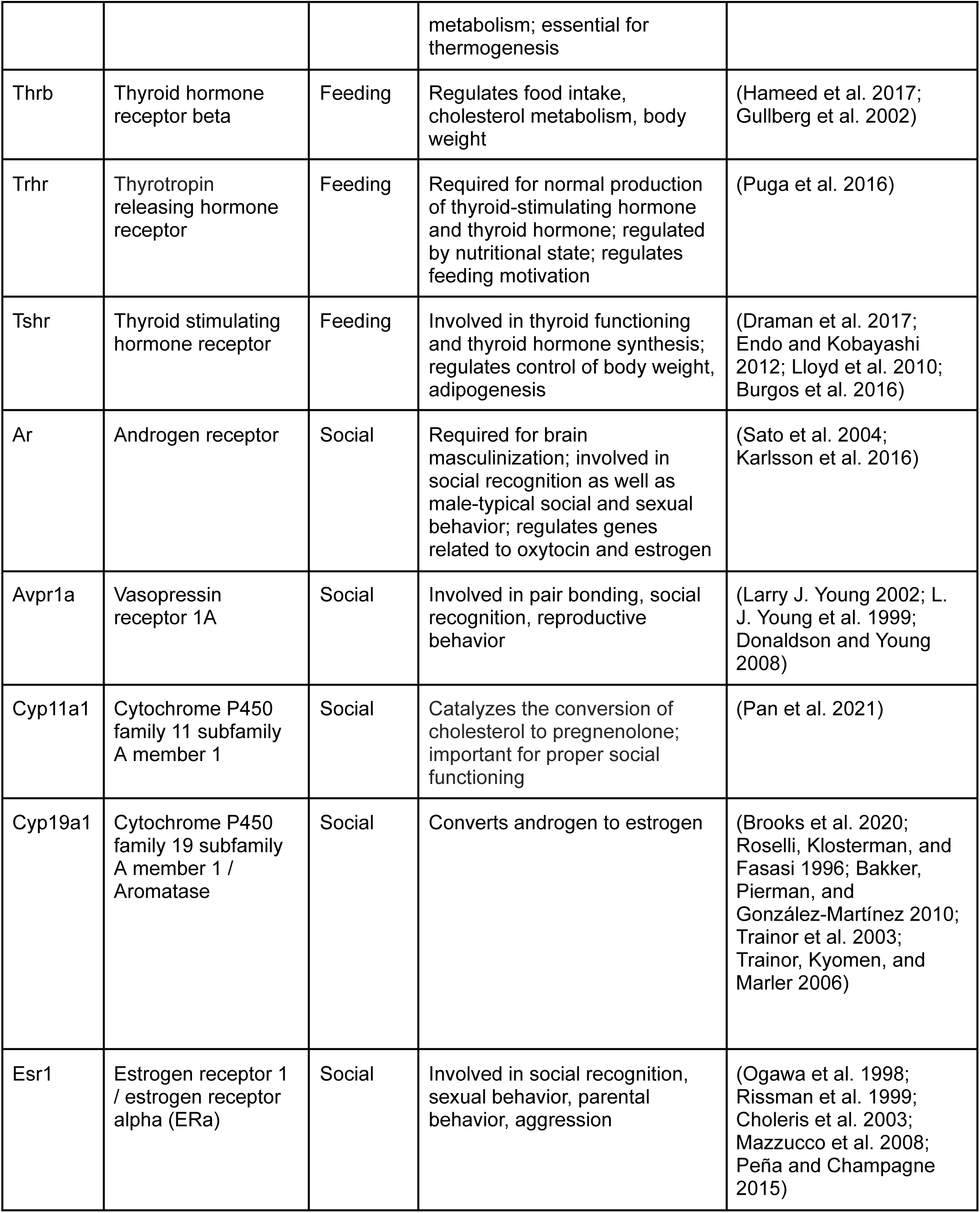
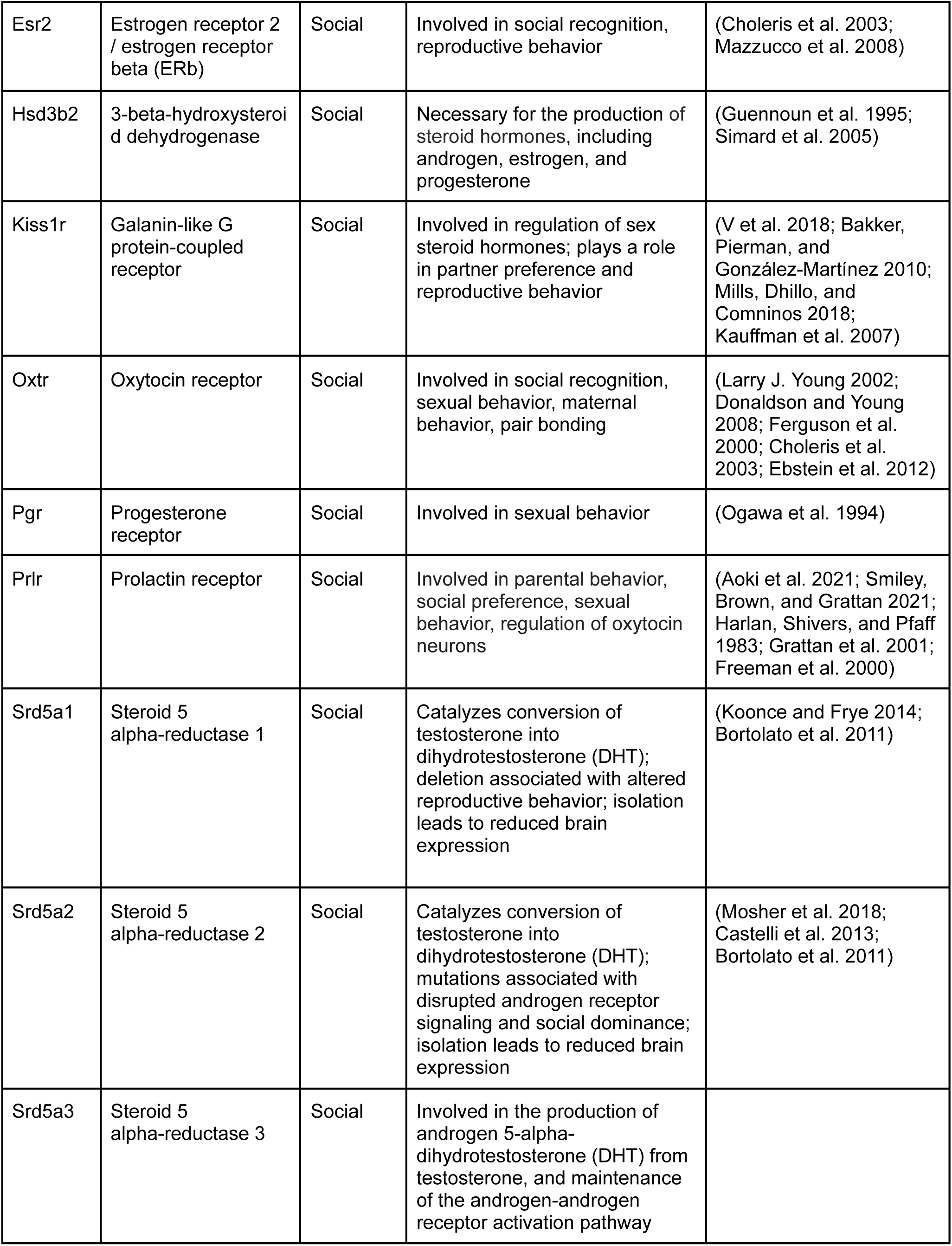
Gene list for Figure 4.

**Table S26:**
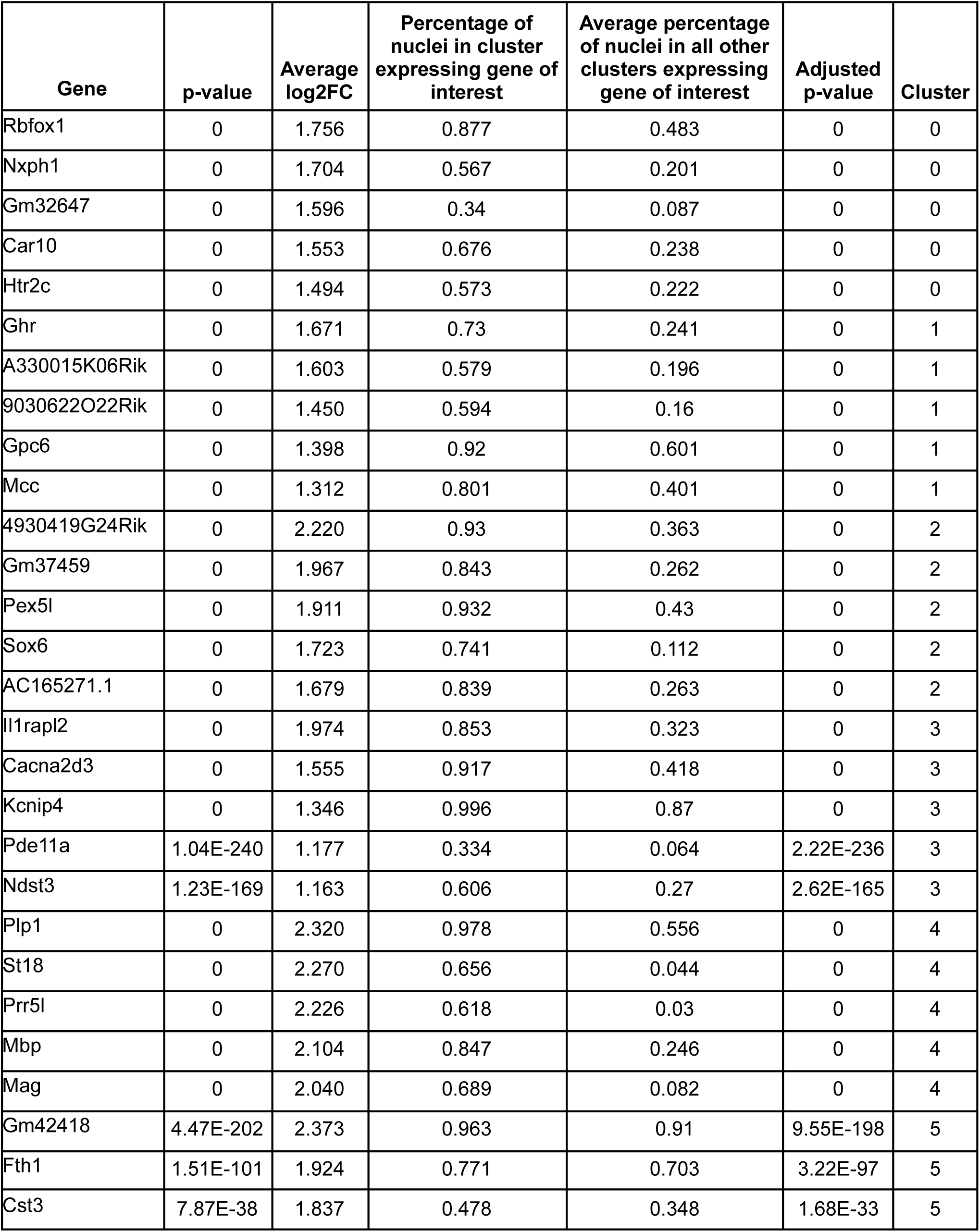
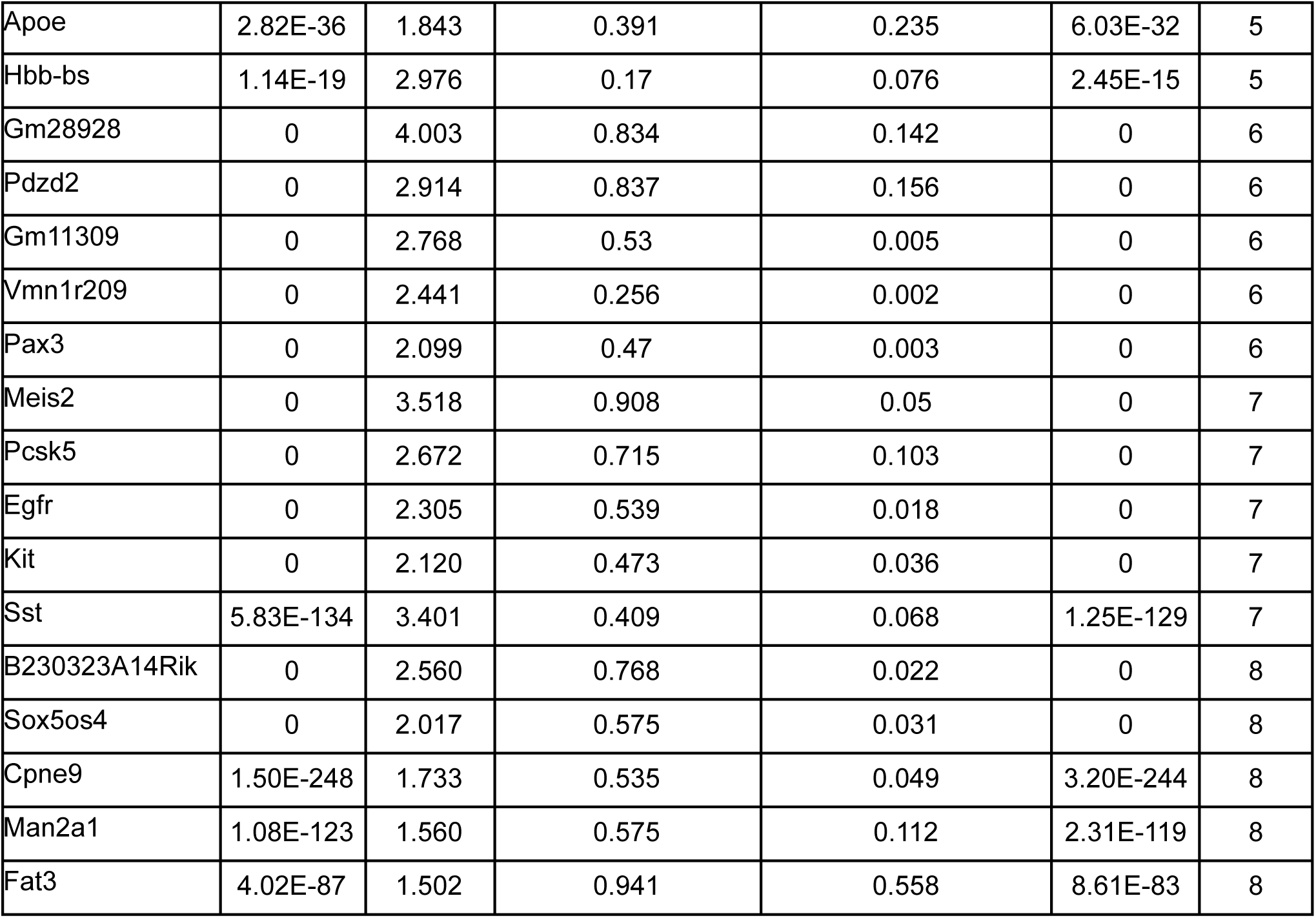
Marker genes for DAT+ subclusters shown in Figure 4E-F.

**Table S27:** Statistics for Figure 4.

| Figure | Description | Sample size | Statistical test | Statistical values | Test Statistic | p value | Significance |
| --- | --- | --- | --- | --- | --- | --- | --- |
| Fig. 4K | Percentage of DA nuclei expressing both genes (row and column) | 14,208 nuclei | Each point in the null distribution is constructed by shuffling whether a nucleus expresses each gene in a given pair, then calculating the percentage of nuclei expressing both genes from the population. | P-value is the fraction of simulated values from the null distribution greater than the true percentage of nuclei expressing both genes | NA | See table S28 | See table S28 |
| Fig. 4L | Proportion of feeding/social gene pairs coexpressed more than chance (DA vs GABA) | 264 gene pairs, 264 gene pairs | 2-sided, 2-sample test for equality of proportions with continuity correction | Difference in proportions from 0 | X-squared =1.351 | p=0.245 | ns |
| Fig. 4L | Proportion of feeding/social gene pairs coexpressed more than chance (DA vs glutamate) | 264 gene pairs, 264 gene pairs | 2-sided, 2-sample test for equality of proportions with continuity correction | Difference in proportions from 0 | X-squared =5.954 | p=0.015 | * |
| Fig. 4L | Proportion of feeding/social gene pairs coexpressed more than chance (GABA vs glutamate) | 264 gene pairs, 264 gene pairs | 2-sided, 2-sample test for equality of proportions with continuity correction | Difference in proportions from 0 | X-squared =1.367 | p=0.242 | ns |

**Table S28:**
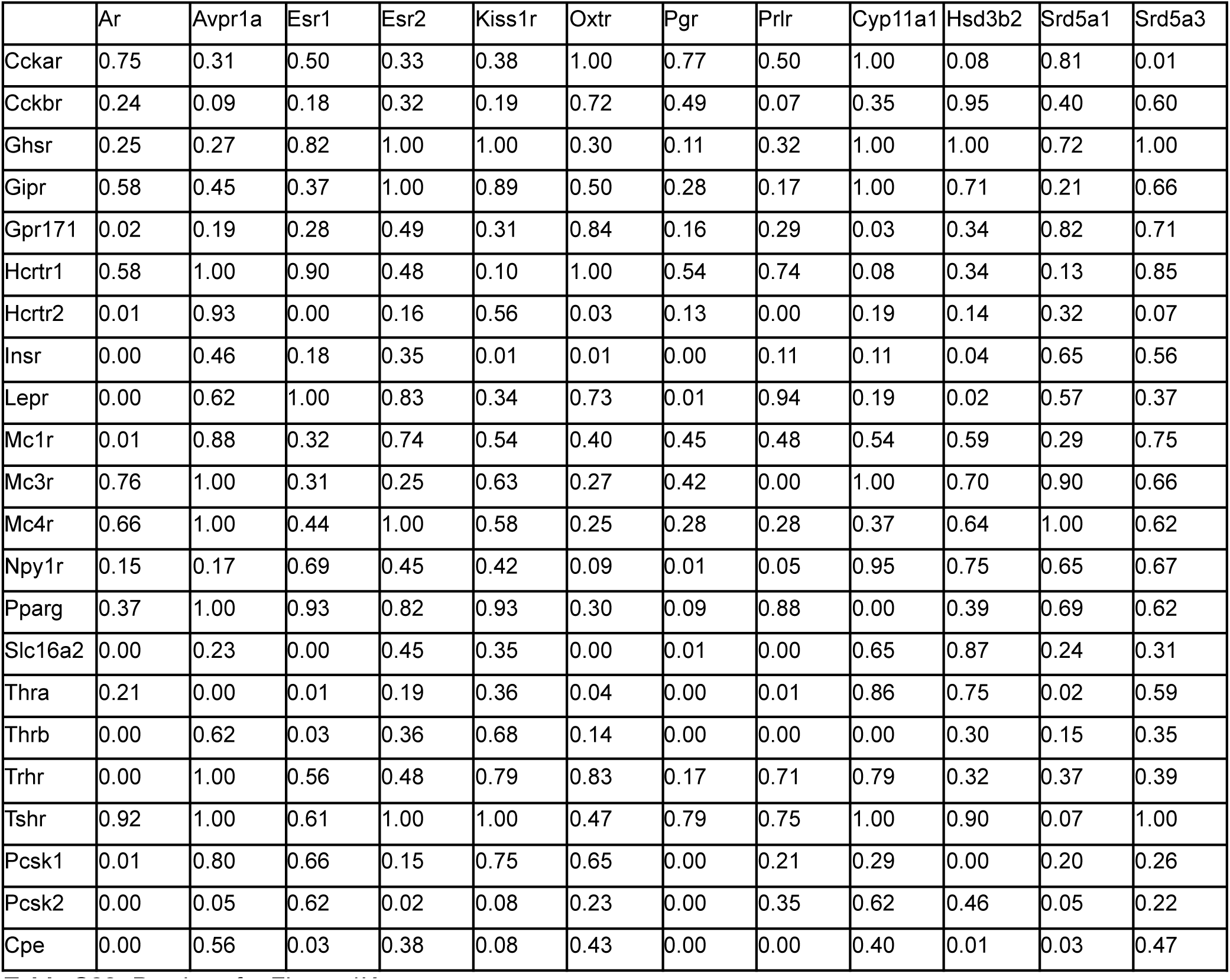
P-values for Figure 4K.

**Table S29:**
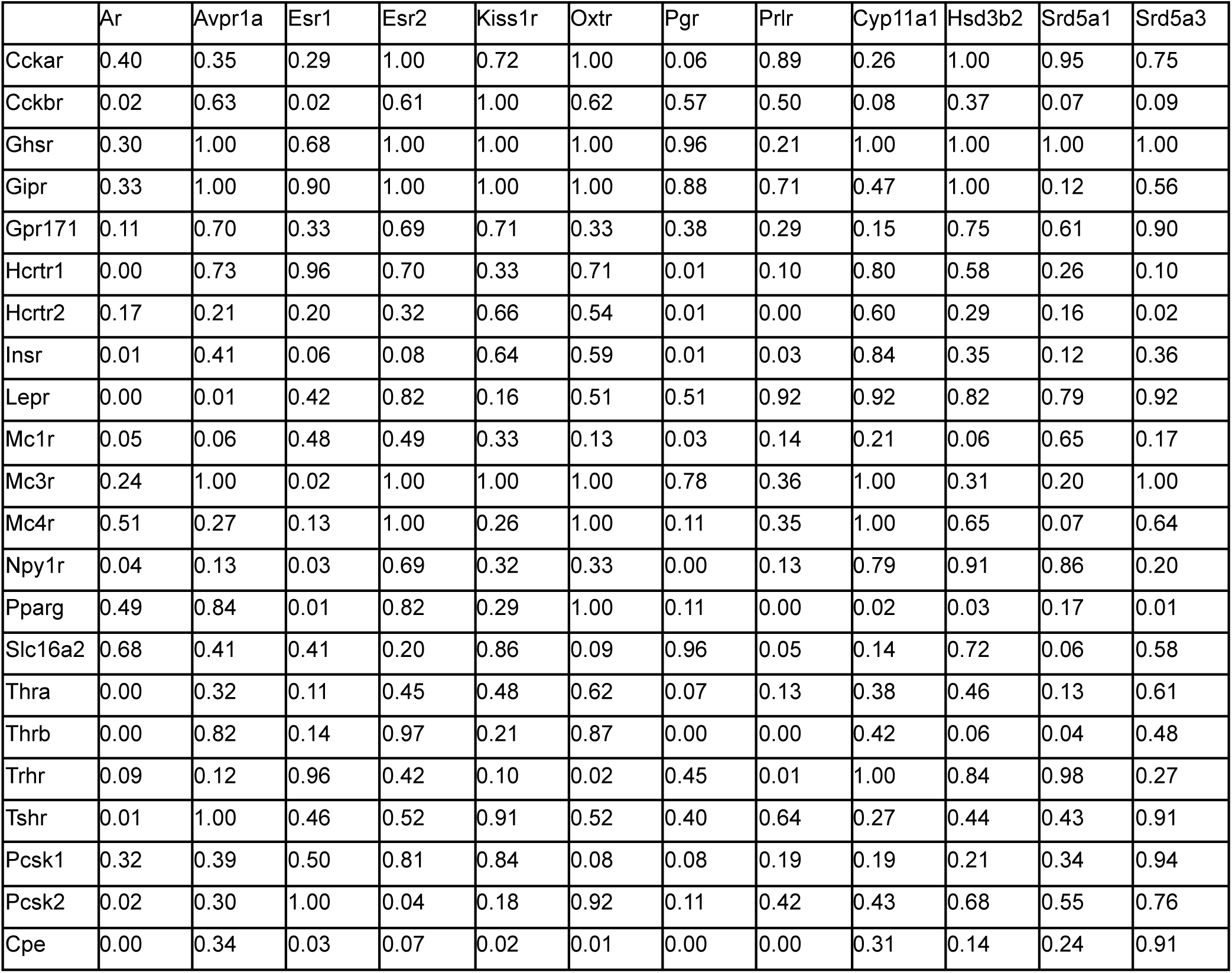
P-values for Figure S16B.

**Table S30:** P-values for Figure S16C.

|  | Ar | Avpr1a | Esr1 | Esr2 | Kiss1r | Oxtr | Pgr | Prlr | Cyp11a1 | Hsd3b2 | Srd5a1 | Srd5a3 |
| --- | --- | --- | --- | --- | --- | --- | --- | --- | --- | --- | --- | --- |
| Cckar | 1.00 | 1.00 | 0.64 | 1.00 | 1.00 | 1.00 | 0.88 | 0.61 | 1.00 | 1.00 | 0.56 | 1.00 |
| Cckbr | 0.37 | 1.00 | 0.84 | 1.00 | 0.15 | 1.00 | 0.55 | 0.92 | 1.00 | 0.58 | 0.32 | 0.90 |
| Ghsr | 0.40 | 1.00 | 1.00 | 1.00 | 1.00 | 1.00 | 0.40 | 1.00 | 0.12 | 1.00 | 0.42 | 1.00 |
| Gipr | 0.17 | 1.00 | 0.86 | 1.00 | 0.42 | 1.00 | 0.98 | 0.18 | 1.00 | 1.00 | 0.43 | 1.00 |
| Gpr171 | 0.90 | 1.00 | 0.98 | 1.00 | 0.22 | 1.00 | 0.05 | 0.23 | 1.00 | 0.45 | 0.15 | 0.84 |
| Hcrtr1 | 0.47 | 1.00 | 0.37 | 1.00 | 0.01 | 0.12 | 0.84 | 0.60 | 0.55 | 0.30 | 0.41 | 0.14 |
| Hcrtr2 | 0.39 | 0.65 | 0.03 | 0.46 | 0.18 | 0.38 | 0.38 | 0.00 | 0.82 | 0.99 | 0.57 | 0.81 |
| Insr | 0.02 | 0.15 | 0.07 | 0.02 | 0.01 | 0.41 | 0.01 | 0.01 | 0.14 | 0.58 | 0.02 | 0.07 |
| Lepr | 0.24 | 0.32 | 0.85 | 0.15 | 0.96 | 0.47 | 0.14 | 0.45 | 0.64 | 0.58 | 0.36 | 0.69 |
| Mc1r | 0.42 | 0.37 | 0.05 | 0.48 | 0.17 | 0.57 | 0.69 | 0.02 | 0.48 | 0.29 | 0.01 | 0.39 |
| Mc3r | 0.25 | 1.00 | 0.38 | 0.05 | 1.00 | 1.00 | 1.00 | 0.64 | 1.00 | 1.00 | 1.00 | 1.00 |
| Mc4r | 0.58 | 1.00 | 0.62 | 1.00 | 1.00 | 1.00 | 0.86 | 0.87 | 1.00 | 0.24 | 0.53 | 0.26 |
| Npy1r | 0.33 | 1.00 | 0.43 | 0.54 | 0.01 | 0.61 | 0.20 | 0.43 | 0.72 | 0.83 | 0.40 | 0.14 |
| Pparg | 0.83 | 0.23 | 0.10 | 0.60 | 0.63 | 0.28 | 0.05 | 0.30 | 0.16 | 0.60 | 0.22 | 0.00 |
| Slc16a2 | 0.03 | 1.00 | 0.61 | 0.69 | 0.44 | 0.01 | 0.23 | 0.05 | 0.66 | 0.23 | 0.92 | 0.90 |
| Thra | 0.06 | 0.43 | 0.63 | 0.17 | 0.41 | 0.08 | 0.01 | 0.09 | 0.14 | 0.61 | 0.39 | 0.10 |
| Thrb | 0.00 | 1.00 | 0.64 | 0.28 | 0.11 | 0.11 | 0.21 | 0.00 | 0.15 | 0.57 | 0.26 | 0.32 |
| Trhr | 0.70 | 1.00 | 0.63 | 0.09 | 0.01 | 0.00 | 0.27 | 0.39 | 0.82 | 0.91 | 0.92 | 0.29 |
| Tshr | 0.32 | 0.08 | 0.67 | 1.00 | 1.00 | 1.00 | 0.95 | 0.32 | 1.00 | 0.47 | 0.85 | 0.51 |
| Pcsk1 | 0.09 | 0.66 | 0.24 | 0.02 | 0.34 | 0.42 | 0.57 | 0.05 | 0.35 | 0.83 | 0.01 | 0.22 |
| Pcsk2 | 0.00 | 0.76 | 0.71 | 0.92 | 0.04 | 0.33 | 0.12 | 0.08 | 0.09 | 0.22 | 0.70 | 0.68 |
| Cpe | 0.02 | 0.67 | 0.02 | 0.01 | 0.15 | 0.81 | 0.29 | 0.00 | 0.27 | 0.65 | 0.13 | 0.75 |

## References

Abizaid, Alfonso, Zhong-Wu Liu, Zane B. Andrews, Marya Shanabrough, Erzsebet Borok, John D. Elsworth, Robert H. Roth, et al. 2006. “Ghrelin Modulates the Activity and Synaptic Input Organization of Midbrain Dopamine Neurons While Promoting Appetite.” The Journal of Clinical Investigation 116 (12): 3229–39.

Adkins-Regan, Elizabeth. 2009. “Neuroendocrinology of Social Behavior.” ILAR Journal / National Research Council, Institute of Laboratory Animal Resources 50 (1): 5–14.

Adriaenssens, Alice E., Emma K. Biggs, Tamana Darwish, John Tadross, Tanmay Sukthankar, Milind Girish, Joseph Polex-Wolf, et al. 2019. “Glucose-Dependent Insulinotropic Polypeptide Receptor-Expressing Cells in the Hypothalamus Regulate Food Intake.” Cell Metabolism. https://doi.org/10.1016/j.cmet.2019.07.013.

Ahima, R. S., D. Prabakaran, C. Mantzoros, D. Qu, B. Lowell, E. Maratos-Flier, and J. S. Flier. 1996. “Role of Leptin in the Neuroendocrine Response to Fasting.” Nature 382 (6588): 250–52.

Anderegg, Angela, Jean-Francois Poulin, and Rajeshwar Awatramani. 2015. “Molecular Heterogeneity of Midbrain Dopaminergic Neurons--Moving toward Single Cell Resolution.” FEBS Letters 589 (24 Pt A): 3714–26.

Andermann, Mark L., and Bradford B. Lowell. 2017. “Toward a Wiring Diagram Understanding of Appetite Control.” Neuron 95 (4): 757–78.

Aoki, Mari, Igor Gamayun, Amanda Wyatt, Ramona Grünewald, Martin Simon-Thomas, Stephan E. Philipp, Oliver Hummel, et al. 2021. “Prolactin-Sensitive Olfactory Sensory Neurons Regulate Male Preference in Female Mice by Modulating Responses to Chemosensory Cues.” Science Advances 7 (41): eabg4074.

Atasoy, Deniz, J. Nicholas Betley, Helen H. Su, and Scott M. Sternson. 2012. “Deconstruction of a Neural Circuit for Hunger.” Nature. https://doi.org/10.1038/nature11270.

Bakker, Julie, Sylvie Pierman, and David González-Martínez. 2010. “Effects of Aromatase Mutation (ArKO) on the Sexual Differentiation of Kisspeptin Neuronal Numbers and Their Activation by Same versus Opposite Sex Urinary Pheromones.” Hormones and Behavior. https://doi.org/10.1016/j.yhbeh.2009.11.005.

Bariselli, Sebastiano, Hanna Hörnberg, Clément Prévost-Solié, Stefano Musardo, Laetitia Hatstatt-Burklé, Peter Scheiffele, and Camilla Bellone. 2018. “Role of VTA Dopamine Neurons and Neuroligin 3 in Sociability Traits Related to Nonfamiliar Conspecific Interaction.” Nature Communications 9 (1): 3173.

Barter, Joseph W., Suellen Li, Dongye Lu, Ryan A. Bartholomew, Mark A. Rossi, Charles T. Shoemaker, Daniel Salas-Meza, Erin Gaidis, and Henry H. Yin. 2015. “Beyond Reward Prediction Errors: The Role of Dopamine in Movement Kinematics.” Frontiers in Integrative Neuroscience 9 (May): 39.

Beglinger, C., L. Degen, D. Matzinger, M. D’Amato, and J. Drewe. 2001. “Loxiglumide, a CCK-A Receptor Antagonist, Stimulates Calorie Intake and Hunger Feelings in Humans.” American Journal of Physiology. Regulatory, Integrative and Comparative Physiology 280 (4): R1149–54.

Beier, Kevin T., Elizabeth E. Steinberg, Katherine E. DeLoach, Stanley Xie, Kazunari Miyamichi, Lindsay Schwarz, Xiaojing J. Gao, Eric J. Kremer, Robert C. Malenka, and Liqun Luo. 2015. “Circuit Architecture of VTA Dopamine Neurons Revealed by Systematic Input-Output Mapping.” Cell 162 (3): 622–34.

Bortolato, Marco, Paola Devoto, Paola Roncada, Roberto Frau, Giovanna Flore, Pierluigi Saba, Giuseppa Pistritto, et al. 2011. “Isolation Rearing-Induced Reduction of Brain 5α-Reductase Expression: Relevance to Dopaminergic Impairments.” Neuropharmacology 60 (7-8): 1301–8.

Branch, Sarah Y., R. Brandon Goertz, Amanda L. Sharpe, Janie Pierce, Sudip Roy, Daijin Ko, Carlos A. Paladini, and Michael J. Beckstead. 2013. “Food Restriction Increases Glutamate Receptor-Mediated Burst Firing of Dopamine Neurons.” The Journal of Neuroscience: The Official Journal of the Society for Neuroscience 33 (34): 13861–72.

Brooks, David C., John S. Coon V, Cihangir M. Ercan, Xia Xu, Hongxin Dong, Jon E. Levine, Serdar E. Bulun, and Hong Zhao. 2020. “Brain Aromatase and the Regulation of Sexual Activity in Male Mice.” Endocrinology 161 (10). https://doi.org/10.1210/endocr/bqaa137.

Brüning, J. C., D. Gautam, D. J. Burks, J. Gillette, M. Schubert, P. C. Orban, R. Klein, W. Krone, D. Müller-Wieland, and C. R. Kahn. 2000. “Role of Brain Insulin Receptor in Control of Body Weight and Reproduction.” Science 289 (5487): 2122–25.

Burgos, Jonathan R., Britt-Marie Iresjö, Sara Wärnåker, and Ulrika Smedh. 2016. “Presence of TSH Receptors in Discrete Areas of the Hypothalamus and Caudal Brainstem with Relevance for Feeding controls—Support for Functional Significance.” Brain Research. https://doi.org/10.1016/j.brainres.2016.04.007.

Butler, Andrew, Paul Hoffman, Peter Smibert, Efthymia Papalexi, and Rahul Satija. 2018. “Integrating Single-Cell Transcriptomic Data across Different Conditions, Technologies, and Species.” Nature Biotechnology 36 (5): 411–20.

Byers, Shannon L., Michael V. Wiles, Sadie L. Dunn, and Robert A. Taft. 2012. “Mouse Estrous Cycle Identification Tool and Images.” PloS One 7 (4): e35538.

Cai, Lili X., Katherine Pizano, Gregory W. Gundersen, Cameron L. Hayes, Weston T. Fleming, Sebastian Holt, Julia M. Cox, and Ilana B. Witten. 2020. “Distinct Signals in Medial and Lateral VTA Dopamine Neurons Modulate Fear Extinction at Different Times.” eLife 9 (June). https://doi.org/10.7554/eLife.54936.

Campbell, Jonathan E. 2021. “Targeting the GIPR for Obesity: To Agonize or Antagonize? Potential Mechanisms.” Molecular Metabolism 46 (April): 101139.

Cano, V., B. Merino, L. Ezquerra, B. Somoza, and M. Ruiz-Gayo. 2008. “A Cholecystokinin-1 Receptor Agonist (CCK-8) Mediates Increased Permeability of Brain Barriers to Leptin.” British Journal of Pharmacology 154 (5): 1009–15.

Castelli, M. Paola, >Alberto Casti, Angelo Casu, Roberto Frau, Marco Bortolato, Saturnino Spiga, and Maria Grazia Ennas. 2013. “Regional Distribution of 5α-Reductase Type 2 in the Adult Rat Brain: An Immunohistochemical Analysis.” Psychoneuroendocrinology 38 (2): 281–93.

Cawley, Niamh X., Jiechun Zhou, Joanna M. Hill, Daniel Abebe, Sylvie Romboz, Tulin Yanik, Ramona M. Rodriguiz, William C. Wetsel, and Y. Peng Loh. 2004. “The Carboxypeptidase E Knockout Mouse Exhibits Endocrinological and Behavioral Deficits.” Endocrinology 145 (12): 5807–19.

Chabout, Jonathan, Abhra Sarkar, David B. Dunson, and Erich D. Jarvis. 2015. “Male Mice Song Syntax Depends on Social Contexts and Influences Female Preferences.” Frontiers in Behavioral Neuroscience 9 (April): 76.

Chen, A. S., J. M. Metzger, M. E. Trumbauer, X. M. Guan, H. Yu, E. G. Frazier, D. J. Marsh, et al. 2000. “Role of the Melanocortin-4 Receptor in Metabolic Rate and Food Intake in Mice.” Transgenic Research 9 (2): 145–54.

Chen, Yanyun, Changzhi Hu, Chiun-Kang Hsu, Qing Zhang, Chen Bi, Mark Asnicar, Hansen M. Hsiung, et al. 2002. “Targeted Disruption of the Melanin-Concentrating Hormone Receptor-1 Results in Hyperphagia and Resistance to Diet-Induced Obesity.” Endocrinology 143 (7): 2469–77.

Choi, Jung Yoon, Hee Jae Jang, Sharon Ornelas, Weston T. Fleming, Daniel Fürth, Jennifer Au, Akhil Bandi, Esteban A. Engel, and Ilana B. Witten. 2020. “A Comparison of Dopaminergic and Cholinergic Populations Reveals Unique Contributions of VTA Dopamine Neurons to Short-Term Memory.” Cell Reports. https://doi.org/10.1016/j.celrep.2020.108492.

Choleris, Elena, Jan-Ake Gustafsson, Kenneth S. Korach, Louis J. Muglia, Donald W. Pfaff, and Sonoko Ogawa. 2003. “An Estrogen-Dependent Four-Gene Micronet Regulating Social Recognition: A Study with Oxytocin and Estrogen Receptor-Alpha and -Beta Knockout Mice.” Proceedings of the National Academy of Sciences of the United States of America 100 (10): 6192–97.

Cohen, Jeremiah Y., Sebastian Haesler, Linh Vong, Bradford B. Lowell, and Naoshige Uchida. 2012. “Neuron-Type-Specific Signals for Reward and Punishment in the Ventral Tegmental Area.” Nature 482 (7383): 85–88.

Coll, Anthony P., Giles S. H. Yeo, I. Sadaf Farooqi, and Stephen O’Rahilly. 2008. “SnapShot: The Hormonal Control of Food Intake.” Cell 135 (3): 572.e1–2.

Collins, Anne L., and Benjamin T. Saunders. 2020. “Heterogeneity in Striatal Dopamine Circuits: Form and Function in Dynamic Reward Seeking.” Journal of Neuroscience Research 98 (6): 1046–69.

Crawley, J. N., and M. C. Beinfeld. 1983. “Rapid Development of Tolerance to the Behavioural Actions of Cholecystokinin.” Nature 302 (5910): 703–6.

Dabney, Will, Zeb Kurth-Nelson, Naoshige Uchida, Clara Kwon Starkweather, Demis Hassabis, Rémi Munos, and Matthew Botvinick. 2020. “A Distributional Code for Value in Dopamine-Based Reinforcement Learning.” Nature 577 (7792): 671–75.

Dai, Bing, Fangmiao Sun, Xiaoyu Tong, Yizhuo Ding, Amy Kuang, Takuya Osakada, Yulong Li, and Dayu Lin. 2022. “Responses and Functions of Dopamine in Nucleus Accumbens Core during Social Behaviors.” Cell Reports 40 (8). https://doi.org/10.1016/j.celrep.2022.111246.

Daigle, Tanya L., Linda Madisen, Travis A. Hage, Matthew T. Valley, Ulf Knoblich, Rylan S. Larsen, Marc M. Takeno, et al. 2018. “A Suite of Transgenic Driver and Reporter Mouse Lines with Enhanced Brain-Cell-Type Targeting and Functionality.” Cell 174 (2): 465–80.e22.

Damsma, G., J. G. Pfaus, D. Wenkstern, A. G. Phillips, and H. C. Fibiger. 1992. “Sexual Behavior Increases Dopamine Transmission in the Nucleus Accumbens and Striatum of Male Rats: Comparison with Novelty and Locomotion.” Behavioral Neuroscience 106 (1): 181–91.

Dey, Sandeepa, Pablo Chamero, James K. Pru, Ming-Shan Chien, Ximena Ibarra-Soria, Kathryn R. Spencer, Darren W. Logan, Hiroaki Matsunami, John J. Peluso, and Lisa Stowers. 2015. “Cyclic Regulation of Sensory Perception by a Female Hormone Alters Behavior.” Cell 161 (6): 1334–44.

Di Cosmo, Caterina, Xiao-Hui Liao, Alexandra M. Dumitrescu, Nancy J. Philp, Roy E. Weiss, and Samuel Refetoff. 2010. “Mice Deficient in MCT8 Reveal a Mechanism Regulating Thyroid Hormone Secretion.” The Journal of Clinical Investigation 120 (9): 3377–88.

Di Cosmo, Caterina, Xiao-Hui Liao, Honggang Ye, Alfonso Massimiliano Ferrara, Roy E. Weiss, Samuel Refetoff, and Alexandra M. Dumitrescu. 2013. “Mct8-Deficient Mice Have Increased Energy Expenditure and Reduced Fat Mass That Is Abrogated by Normalization of Serum T3 Levels.” Endocrinology 154 (12): 4885–95.

Dölen, Gül, Ayeh Darvishzadeh, Kee Wui Huang, and Robert C. Malenka. 2013. “Social Reward Requires Coordinated Activity of Nucleus Accumbens Oxytocin and Serotonin.” Nature 501 (7466): 179–84.

Domingos, Ana I., Jake Vaynshteyn, Henning U. Voss, Xueying Ren, Viviana Gradinaru, Feng Zang, Karl Deisseroth, Ivan E. de Araujo, and Jeffrey Friedman. 2011. “Leptin Regulates the Reward Value of Nutrient.” Nature Neuroscience 14 (12): 1562–68.

Donaldson, Zoe R., and Larry J. Young. 2008. “Oxytocin, Vasopressin, and the Neurogenetics of Sociality.” Science. https://doi.org/10.1126/science.1158668.

Dowsett, Georgina K. C., Brian Y. H. Lam, John A. Tadross, Irene Cimino, Debra Rimmington, Anthony P. Coll, Joseph Polex-Wolf, Lotte Bjerre Knudsen, Charles Pyke, and Giles S. H. Yeo. 2021. “A Survey of the Mouse Hindbrain in the Fed and Fasted States Using Single-Nucleus RNA Sequencing.” Molecular Metabolism 53 (November): 101240.

Draman, Mohd Shazli, Michael Stechman, David Scott-Coombes, Colin M. Dayan, Dafydd Aled Rees, Marian Ludgate, and Lei Zhang. 2017. “The Role of Thyrotropin Receptor Activation in Adipogenesis and Modulation of Fat Phenotype.” Frontiers in Endocrinology. https://doi.org/10.3389/fendo.2017.00083.

Dunigan, Anna I., David P. Olson, and Aaron G. Roseberry. 2021. “VTA MC3R Neurons Control Feeding in an Activity- and Sex-Dependent Manner in Mice.” Neuropharmacology 197 (October): 108746.

Dunigan, Anna I., and Aaron G. Roseberry. 2022. “Actions of Feeding-Related Peptides on the Mesolimbic Dopamine System in Regulation of Natural and Drug Rewards.” Addiction Neuroscience. https://doi.org/10.1016/j.addicn.2022.100011.

Ebstein, Richard P., Ariel Knafo, David Mankuta, Soo Hong Chew, and Poh San Lai. 2012. “The Contributions of Oxytocin and Vasopressin Pathway Genes to Human Behavior.” Hormones and Behavior 61 (3): 359–79.

Egecioglu, Emil, Elisabet Jerlhag, Nicolas Salomé, Karolina P. Skibicka, David Haage, Mohammad Bohlooly-Y, Daniel Andersson, et al. 2010. “Ghrelin Increases Intake of Rewarding Food in Rodents.” Addiction Biology. https://doi.org/10.1111/j.1369-1600.2010.00216.x.

Endo, Toyoshi, and Tetsuro Kobayashi. 2012. “Expression of Functional TSH Receptor in White Adipose Tissues of Hyt/hyt Mice Induces Lipolysis in Vivo.” American Journal of Physiology-Endocrinology and Metabolism. https://doi.org/10.1152/ajpendo.00572.2011.

Engelhard, Ben, Joel Finkelstein, Julia Cox, Weston Fleming, Hee Jae Jang, Sharon Ornelas, Sue Ann Koay, et al. 2019. “Specialized Coding of Sensory, Motor and Cognitive Variables in VTA Dopamine Neurons.” Nature 570 (7762): 509–13.

Eshel, Neir, Ju Tian, Michael Bukwich, and Naoshige Uchida. 2016. “Dopamine Neurons Share Common Response Function for Reward Prediction Error.” Nature Neuroscience 19 (3): 479–86.

Farooqi, I. Sadaf, Edward Bullmore, Julia Keogh, Jonathan Gillard, Stephen O’Rahilly, and Paul C. Fletcher. 2007. “Leptin Regulates Striatal Regions and Human Eating Behavior.” Science 317 (5843): 1355.

Ferguson, J. N., L. J. Young, E. F. Hearn, M. M. Matzuk, T. R. Insel, and J. T. Winslow. 2000. “Social Amnesia in Mice Lacking the Oxytocin Gene.” Nature Genetics 25 (3): 284–88.

Fernandes, Ana B., Joaquim Alves da Silva, Joana Almeida, Guohong Cui, Charles R. Gerfen, Rui M. Costa, and Albino J. Oliveira-Maia. 2020. “Postingestive Modulation of Food Seeking Depends on Vagus-Mediated Dopamine Neuron Activity.” Neuron 106 (5): 778–88.e6.

Figlewicz, D. P., M. S. Higgins, S. B. Ng-Evans, and P. J. Havel. 2001. “Leptin Reverses Sucrose-Conditioned Place Preference in Food-Restricted Rats.” Physiology & Behavior 73 (1-2): 229–34.

Fontes, Klaus N., Adriana Cabanelas, Flavia F. Bloise, Cherley Borba Vieira de Andrade, Luana L. Souza, Marianna Wilieman, Isis H. Trevenzoli, et al. 2017. “Differential Regulation of Thyroid Hormone Metabolism Target Genes during Non-Thyroidal [corrected] Illness Syndrome Triggered by Fasting or Sepsis in Adult Mice.” Frontiers in Physiology 8 (October): 828.

Freeman, M. E., B. Kanyicska, A. Lerant, and G. Nagy. 2000. “Prolactin: Structure, Function, and Regulation of Secretion.” Physiological Reviews 80 (4): 1523–1631.

Fricker, L. D., Y. L. Berman, E. H. Leiter, and L. A. Devi. 1996. “Carboxypeptidase E Activity Is Deficient in Mice with the Fat Mutation. Effect on Peptide Processing.” The Journal of Biological Chemistry 271 (48): 30619–24.

Fulton, Stephanie, Pavlos Pissios, Ramon Pinol Manchon, Linsey Stiles, Lauren Frank, Emmanuel N. Pothos, Eleftheria Maratos-Flier, and Jeffrey S. Flier. 2006. “Leptin Regulation of the Mesoaccumbens Dopamine Pathway.” Neuron 51 (6): 811–22.

Fürth, Daniel, Thomas Vaissière, Ourania Tzortzi, Yang Xuan, Antje Märtin, Iakovos Lazaridis, Giada Spigolon, et al. 2018. “An Interactive Framework for Whole-Brain Maps at Cellular Resolution.” Nature Neuroscience 21 (1): 139–49.

Garfield, Alastair S., Chia Li, Joseph C. Madara, Bhavik P. Shah, Emily Webber, Jennifer S. Steger, John N. Campbell, et al. 2015. “A Neural Basis for Melanocortin-4 Receptor–regulated Appetite.” Nature Neuroscience 18 (6): 863–71.

Georgescu, Dan, Robert M. Sears, Jonathan D. Hommel, Michel Barrot, Carlos A. Bolaños, Donald J. Marsh, Maria A. Bednarek, et al. 2005. “The Hypothalamic Neuropeptide Melanin-Concentrating Hormone Acts in the Nucleus Accumbens to Modulate Feeding Behavior and Forced-Swim Performance.” The Journal of Neuroscience: The Official Journal of the Society for Neuroscience 25 (11): 2933–40.

Gomes, Ivone, Dipendra K. Aryal, Jonathan H. Wardman, Achla Gupta, Khatuna Gagnidze, Ramona M. Rodriguiz, Sanjai Kumar, et al. 2013. “GPR171 Is a Hypothalamic G Protein-Coupled Receptor for BigLEN, a Neuropeptide Involved in Feeding.” Proceedings of the National Academy of Sciences of the United States of America 110 (40): 16211–16.

Grattan, D. R., X. J. Pi, Z. B. Andrews, R. A. Augustine, I. C. Kokay, M. R. Summerfield, B. Todd, and S. J. Bunn. 2001. “Prolactin Receptors in the Brain during Pregnancy and Lactation: Implications for Behavior.” Hormones and Behavior 40 (2): 115–24.

Groppe, Sarah E., Anna Gossen, Lena Rademacher, Alexa Hahn, Luzie Westphal, Gerhard Gründer, and Katja N. Spreckelmeyer. 2013. “Oxytocin Influences Processing of Socially Relevant Cues in the Ventral Tegmental Area of the Human Brain.” Biological Psychiatry 74 (3): 172–79.

Grove, James C. R., Lindsay A. Gray, Naymalis La Santa Medina, Nilla Sivakumar, Jamie S. Ahn, Timothy V. Corpuz, Joshua D. Berke, Anatol C. Kreitzer, and Zachary A. Knight. 2022. “Dopamine Subsystems That Track Internal States.” *Nature*, July. https://doi.org/10.1038/s41586-022-04954-0.

Guan, X. M., H. Yu, O. C. Palyha, K. K. McKee, S. D. Feighner, D. J. Sirinathsinghji, R. G. Smith, L. H. Van der Ploeg, and A. D. Howard. 1997. “Distribution of mRNA Encoding the Growth Hormone Secretagogue Receptor in Brain and Peripheral Tissues.” Brain Research. Molecular Brain Research 48 (1): 23–29.

Guennoun, R., R. J. Fiddes, M. Gouézou, M. Lombès, and E-E Baulieu. 1995. “A Key Enzyme in the Biosynthesis of Neurosteroids, 3β-Hydroxysteroid dehydrogenase/Δ5-Δ4-Isomerase (3β-HSD), Is Expressed in Rat Brain.” Molecular Brain Research. https://doi.org/10.1016/0169-328x(95)00016-l.

Gullberg, Hjalmar, Mats Rudling, Carmen Saltó, Douglas Forrest, Bo Angelin, and Björn Vennström. 2002. “Requirement for Thyroid Hormone Receptor Beta in T3 Regulation of Cholesterol Metabolism in Mice.” Molecular Endocrinology 16 (8): 1767–77.

Gunaydin, Lisa A., Logan Grosenick, Joel C. Finkelstein, Isaac V. Kauvar, Lief E. Fenno, Avishek Adhikari, Stephan Lammel, et al. 2014. “Natural Neural Projection Dynamics Underlying Social Behavior.” Cell 157 (7): 1535–51.

Hameed, Saira, Michael Patterson, Waljit S. Dhillo, Sofia A. Rahman, Yue Ma, Christopher Holton, Apostolos Gogakos, et al. 2017. “Thyroid Hormone Receptor Beta in the Ventromedial Hypothalamus Is Essential for the Physiological Regulation of Food Intake and Body Weight.” Cell Reports 19 (11): 2202–9.

Hamid, Arif A., Michael J. Frank, and Christopher I. Moore. 2021. “Wave-like Dopamine Dynamics as a Mechanism for Spatiotemporal Credit Assignment.” Cell 184 (10): 2733–49.e16.

Harlan, R. E., B. D. Shivers, and D. W. Pfaff. 1983. “Midbrain Microinfusions of Prolactin Increase the Estrogen-Dependent Behavior, Lordosis.” Science 219 (4591): 1451–53.

Harrison, Robert W., John von Neumann, and Oskar Morgenstern. 1945. “The Theory of Games and Economic Behavior.” Journal of Farm Economics 27 (3): 725.

Hassan, Anhar, and Eduardo E. Benarroch. 2015. “Heterogeneity of the Midbrain Dopamine System.” Neurology. https://doi.org/10.1212/wnl.0000000000002137.

Herwig, Annika, Dana Wilson, Tracy J. Logie, Anita Boelen, Peter J. Morgan, Julian G. Mercer, and Perry Barrett. 2009. “Photoperiod and Acute Energy Deficits Interact on Components of the Thyroid Hormone System in Hypothalamic Tanycytes of the Siberian Hamster.” American Journal of Physiology. Regulatory, Integrative and Comparative Physiology 296 (5): R1307–15.

Heuvel, José K. van den, Kara Furman, Myrtille C. R. Gumbs, Leslie Eggels, Darren M. Opland, Benjamin B. Land, Sharon M. Kolk, et al. 2015. “Neuropeptide Y Activity in the Nucleus Accumbens Modulates Feeding Behavior and Neuronal Activity.” Biological Psychiatry 77 (7): 633–41.

Hommel, Jonathan D., Richard Trinko, Robert M. Sears, Dan Georgescu, Zong-Wu Liu, Xiao-Bing Gao, Jeremy J. Thurmon, Michela Marinelli, and Ralph J. DiLeone. 2006. “Leptin Receptor Signaling in Midbrain Dopamine Neurons Regulates Feeding.” Neuron 51 (6): 801–10.

Houk, James C., Joel L. Davis, and David G. Beiser. 1995. Models of Information Processing in the Basal Ganglia. MIT Press.

Howe, M. W., and D. A. Dombeck. 2016. “Rapid Signalling in Distinct Dopaminergic Axons during Locomotion and Reward.” Nature 535 (7613): 505–10.

Hrvatin, Sinisa, Senmiao Sun, Oren F. Wilcox, Hanqi Yao, Aurora J. Lavin-Peter, Marcelo Cicconet, Elena G. Assad, et al. 2020. “Neurons That Regulate Mouse Torpor.” Nature 583 (7814): 115–21.

Hung, Lin W., Sophie Neuner, Jai S. Polepalli, Kevin T. Beier, Matthew Wright, Jessica J. Walsh, Eastman M. Lewis, et al. 2017. “Gating of Social Reward by Oxytocin in the Ventral Tegmental Area.” Science 357 (6358): 1406–11.

Huszar, D., C. A. Lynch, V. Fairchild-Huntress, J. H. Dunmore, Q. Fang, L. R. Berkemeier, W. Gu, et al. 1997. “Targeted Disruption of the Melanocortin-4 Receptor Results in Obesity in Mice.” Cell 88 (1): 131–41.

Isaac, Jennifer, Sonia Karkare, Hymavathy Balasubramanian, Nicholas Schappaugh, Jarildy Javier, Maha Rashid, and Malavika Murugan. 2023. “Sex Differences in Neural Representations of Social and Nonsocial Reward in the Medial Prefrontal Cortex.” bioRxiv. https://doi.org/10.1101/2023.03.09.531947.

Jennings, Joshua H., Christina K. Kim, James H. Marshel, Misha Raffiee, Li Ye, Sean Quirin, Sally Pak, Charu Ramakrishnan, and Karl Deisseroth. 2019. “Interacting Neural Ensembles in Orbitofrontal Cortex for Social and Feeding Behaviour.” Nature. https://doi.org/10.1038/s41586-018-0866-8.

Jerlhag, Elisabet, Emil Egecioglu, Suzanne L. Dickson, Annika Douhan, Lennart Svensson, and Jörgen A. Engel. 2007. “Ghrelin Administration into Tegmental Areas Stimulates Locomotor Activity and Increases Extracellular Concentration of Dopamine in the Nucleus Accumbens.” Addiction Biology 12 (1): 6–16.

Karlsson, Sara A., Erik Studer, Petronella Kettunen, and Lars Westberg. 2016. “Neural Androgen Receptors Modulate Gene Expression and Social Recognition But Not Social Investigation.” Frontiers in Behavioral Neuroscience 10 (March): 41.

Kauffman, Alexander S., Jin Ho Park, Anika A. McPhie-Lalmansingh, Michelle L. Gottsch, Cristian Bodo, John G. Hohmann, Maria N. Pavlova, et al. 2007. “The Kisspeptin Receptor GPR54 Is Required for Sexual Differentiation of the Brain and Behavior.” The Journal of Neuroscience: The Official Journal of the Society for Neuroscience 27 (33): 8826–35.

Kendrick, K. M., and R. F. Drewett. 1979. “Testosterone Reduces Refractory Period of Stria Terminalis Neurons in the Rat Brain.” Science 204 (4395): 877–79.

Koonce, Carolyn J., and Cheryl A. Frye. 2014. “Female Mice with Deletion of Type One 5α-Reductase Have Reduced Reproductive Responding during Proestrus and after Hormone-Priming.” Pharmacology, Biochemistry, and Behavior 122 (July): 20–29.

Krashes, Michael J., Bhavik P. Shah, Shuichi Koda, and Bradford B. Lowell. 2013. “Rapid versus Delayed Stimulation of Feeding by the Endogenously Released AgRP Neuron Mediators GABA, NPY, and AgRP.” Cell Metabolism 18 (4): 588–95.

Kremer, Yves, Jérôme Flakowski, Clément Rohner, and Christian Lüscher. 2020. “Context-Dependent Multiplexing by Individual VTA Dopamine Neurons.” The Journal of Neuroscience: The Official Journal of the Society for Neuroscience, August. https://doi.org/10.1523/JNEUROSCI.0502-20.2020.

Labouèbe, Gwenaël, Shuai Liu, Carine Dias, Haiyan Zou, Jovi C. Y. Wong, Subashini Karunakaran, Susanne M. Clee, Anthony G. Phillips, Benjamin Boutrel, and Stephanie L. Borgland. 2013. “Insulin Induces Long-Term Depression of Ventral Tegmental Area Dopamine Neurons via Endocannabinoids.” Nature Neuroscience 16 (3): 300–308.

La Manno, Gioele, Daniel Gyllborg, Simone Codeluppi, Kaneyasu Nishimura, Carmen Salto, Amit Zeisel, Lars E. Borm, et al. 2016. “Molecular Diversity of Midbrain Development in Mouse, Human, and Stem Cells.” Cell 167 (2): 566–80.e19.

Lam, Brian Y. H., Irene Cimino, Joseph Polex-Wolf, Sara Nicole Kohnke, Debra Rimmington, Valentine Iyemere, Nicholas Heeley, et al. 2017. “Heterogeneity of Hypothalamic pro-Opiomelanocortin-Expressing Neurons Revealed by Single-Cell RNA Sequencing.” Molecular Metabolism 6 (5): 383–92.

Lammel, Stephan, Daniela I. Ion, Jochen Roeper, and Robert C. Malenka. 2011. “Projection-Specific Modulation of Dopamine Neuron Synapses by Aversive and Rewarding Stimuli.” Neuron 70 (5): 855–62.

Lammel, Stephan, Byung Kook Lim, Chen Ran, Kee Wui Huang, Michael J. Betley, Kay M. Tye, Karl Deisseroth, and Robert C. Malenka. 2012. “Input-Specific Control of Reward and Aversion in the Ventral Tegmental Area.” Nature 491 (7423): 212–17.

Lammel, Stephan, Elizabeth E. Steinberg, Csaba Földy, Nicholas R. Wall, Kevin Beier, Liqun Luo, and Robert C. Malenka. 2015. “Diversity of Transgenic Mouse Models for Selective Targeting of Midbrain Dopamine Neurons.” Neuron 85 (2): 429–38.

Landreth, Anthony, and John Bickle. 2008. “NEUROECONOMICS, NEUROPHYSIOLOGY AND THE COMMON CURRENCY HYPOTHESIS.” Economics & Philosophy 24 (3): 419–29.

Lee, Christopher R., Alon Chen, and Kay M. Tye. 2021. “The Neural Circuitry of Social Homeostasis: Consequences of Acute versus Chronic Social Isolation.” Cell 184 (6): 1500–1516.

Lee, G. H., R. Proenca, J. M. Montez, K. M. Carroll, J. G. Darvishzadeh, J. I. Lee, and J. M. Friedman. 1996. “Abnormal Splicing of the Leptin Receptor in Diabetic Mice.” Nature 379 (6566): 632–35.

Lee, Rachel S., Ben Engelhard, Ilana B. Witten, and Nathaniel D. Daw. 2022. “A Vector Reward Prediction Error Model Explains Dopaminergic Heterogeneity.” bioRxiv. https://doi.org/10.1101/2022.02.28.482379.

Lee, Rachel S., Marcelo G. Mattar, Nathan F. Parker, Ilana B. Witten, and Nathaniel D. Daw. 2019. “Reward Prediction Error Does Not Explain Movement Selectivity in DMS-Projecting Dopamine Neurons.” eLife 8 (April). https://doi.org/10.7554/eLife.42992.

Lerner, Talia N., Carrie Shilyansky, Thomas J. Davidson, Kathryn E. Evans, Kevin T. Beier, Kelly A. Zalocusky, Ailey K. Crow, et al. 2015. “Intact-Brain Analyses Reveal Distinct Information Carried by SNc Dopamine Subcircuits.” Cell 162 (3): 635–47.

Levy, Dino J., and Paul W. Glimcher. 2012. “The Root of All Value: A Neural Common Currency for Choice.” Current Opinion in Neurobiology 22 (6): 1027–38.

Lippert, Rachel N., Kate L. J. Ellacott, and Roger D. Cone. 2014. “Gender-Specific Roles for the Melanocortin-3 Receptor in the Regulation of the Mesolimbic Dopamine System in Mice.” Endocrinology 155 (5): 1718–27.

Lloyd, Anna de, Anna de Lloyd, James Bursell, John W. Gregory, D. Aled Rees, and Marian Ludgate. 2010. “TSH Receptor Activation and Body Composition.” Journal of Endocrinology. https://doi.org/10.1677/joe-09-0262.

Louilot, A., J. L. Gonzalez-Mora, T. Guadalupe, and M. Mas. 1991. “Sex-Related Olfactory Stimuli Induce a Selective Increase in Dopamine Release in the Nucleus Accumbens of Male Rats. A Voltammetric Study.” Brain Research 553 (2): 313–17.

Love, Michael I., Wolfgang Huber, and Simon Anders. 2014. “Moderated Estimation of Fold Change and Dispersion for RNA-Seq Data with DESeq2.” Genome Biology 15 (12): 550.

Lutas, Andrew, Hakan Kucukdereli, Osama Alturkistani, Crista Carty, Arthur U. Sugden, Kayla Fernando, Veronica Diaz, Vanessa Flores-Maldonado, and Mark L. Andermann. 2019. “State-Specific Gating of Salient Cues by Midbrain Dopaminergic Input to Basal Amygdala.” Nature Neuroscience 22 (11): 1820–33.

Macosko, Evan Z., Anindita Basu, Rahul Satija, James Nemesh, Karthik Shekhar, Melissa Goldman, Itay Tirosh, et al. 2015. “Highly Parallel Genome-Wide Expression Profiling of Individual Cells Using Nanoliter Droplets.” Cell 161 (5): 1202–14.

Marie, Linda Ste, Grant I. Miura, Donald J. Marsh, Keith Yagaloff, and Richard D. Palmiter. 2000. “A Metabolic Defect Promotes Obesity in Mice Lacking Melanocortin-4 Receptors.” Proceedings of the National Academy of Sciences 97 (22): 12339–44.

Marinelli, M., and J. E. McCutcheon. 2014. “Heterogeneity of Dopamine Neuron Activity across Traits and States.” Neuroscience. https://doi.org/10.1016/j.neuroscience.2014.07.034.

Marlin, Bianca J., Mariela Mitre, James A. D’amour, Moses V. Chao, and Robert C. Froemke. 2015. “Oxytocin Enables Maternal Behaviour by Balancing Cortical Inhibition.” Nature 520 (7548): 499–504.

Marsh, Donald J., Gunther Hollopeter, Dennis Huszar, Ralph Laufer, Keith A. Yagaloff, Stewart L. Fisher, Paul Burn, and Richard D. Palmiter. 1999. “Response of melanocortin–4 Receptor–deficient Mice to Anorectic and Orexigenic Peptides.” Nature Genetics 21 (1): 119–22.

Martel, P., and M. Fantino. 1996. “Mesolimbic Dopaminergic System Activity as a Function of Food Reward: A Microdialysis Study.” Pharmacology, Biochemistry, and Behavior 53 (1): 221–26.

Matthews, Gillian A., Edward H. Nieh, Caitlin M. Vander Weele, Sarah A. Halbert, Roma V. Pradhan, Ariella S. Yosafat, Gordon F. Glober, et al. 2016. “Dorsal Raphe Dopamine Neurons Represent the Experience of Social Isolation.” Cell. https://doi.org/10.1016/j.cell.2015.12.040.

Mazzone, Christopher M., Jing Liang-Guallpa, Chia Li, Nora S. Wolcott, Montana H. Boone, Morgan Southern, Nicholas P. Kobzar, et al. 2020. “High-Fat Food Biases Hypothalamic and Mesolimbic Expression of Consummatory Drives.” Nature Neuroscience 23 (10): 1253–66.

Mazzucco, Christine A., Hope A. Walker, Jodi L. Pawluski, Stephanie E. Lieblich, and Liisa A. M. Galea. 2008. “ERalpha, but Not ERbeta, Mediates the Expression of Sexual Behavior in the Female Rat.” Behavioural Brain Research 191 (1): 111–17.

McHenry, Jenna A., James M. Otis, Mark A. Rossi, J. Elliott Robinson, Oksana Kosyk, Noah W. Miller, Zoe A. McElligott, Evgeny A. Budygin, David R. Rubinow, and Garret D. Stuber. 2017. “Hormonal Gain Control of a Medial Preoptic Area Social Reward Circuit.” Nature Neuroscience 20 (3): 449–58.

McNamara, John M., and Alasdair I. Houston. 1986. “The Common Currency for Behavioral Decisions.” The American Naturalist 127 (3): 358–78.

Mills, Edouard G. A., Waljit S. Dhillo, and Alexander N. Comninos. 2018. “Kisspeptin and the Control of Emotions, Mood and Reproductive Behaviour.” The Journal of Endocrinology 239 (1): R1–12.

Mitra, Sudha Warrier, Elena Hoskin, Joel Yudkovitz, Lisset Pear, Hilary A. Wilkinson, Shinji Hayashi, Donald W. Pfaff, et al. 2003. “Immunolocalization of Estrogen Receptor Beta in the Mouse Brain: Comparison with Estrogen Receptor Alpha.” Endocrinology 144 (5): 2055–67.

Mosher, Laura J., Sean C. Godar, Marc Morissette, Kenneth M. McFarlin, Simona Scheggi, Carla Gambarana, Stephen C. Fowler, Thérèse Di Paolo, and Marco Bortolato. 2018. “Steroid 5α-Reductase 2 Deficiency Leads to Reduced Dominance-Related and Impulse-Control Behaviors.” Psychoneuroendocrinology 91 (May): 95–104.

Mumtaz, Faiza, Muhammad Imran Khan, Muhammad Zubair, and Ahmad Reza Dehpour. 2018. “Neurobiology and Consequences of Social Isolation Stress in Animal Model-A Comprehensive Review.” Biomedicine & Pharmacotherapy = Biomedecine & Pharmacotherapie 105 (September): 1205–22.

Nakazato, M., N. Murakami, Y. Date, M. Kojima, H. Matsuo, K. Kangawa, and S. Matsukura. 2001. “A Role for Ghrelin in the Central Regulation of Feeding.” Nature 409 (6817): 194–98.

Neunuebel, Joshua P., Adam L. Taylor, Ben J. Arthur, and S. E. Roian Egnor. 2015. “Female Mice Ultrasonically Interact with Males during Courtship Displays.” eLife 4 (May). https://doi.org/10.7554/eLife.06203.

O’Connell, Lauren A., and Hans A. Hofmann. 2011. “Genes, Hormones, and Circuits: An Integrative Approach to Study the Evolution of Social Behavior.” Frontiers in Neuroendocrinology 32 (3): 320–35.

Ogawa, S., V. Eng, J. Taylor, D. B. Lubahn, K. S. Korach, and D. W. Pfaff. 1998. “Roles of Estrogen Receptor-Alpha Gene Expression in Reproduction-Related Behaviors in Female Mice.” Endocrinology 139 (12): 5070–81.

Ogawa, S., U. E. Olazabal, I. S. Parhar, and D. W. Pfaff. 1994. “Effects of Intrahypothalamic Administration of Antisense DNA for Progesterone Receptor mRNA on Reproductive Behavior and Progesterone Receptor Immunoreactivity in Female Rat.” The Journal of Neuroscience. https://doi.org/10.1523/jneurosci.14-03-01766.1994.

Pachitariu, Marius, Carsen Stringer, Mario Dipoppa, Sylvia Schröder, L. Federico Rossi, Henry Dalgleish, Matteo Carandini, and Kenneth D. Harris. n.d. “Suite2p: Beyond 10,000 Neurons with Standard Two-Photon Microscopy.” https://doi.org/10.1101/061507.

Pandit, Rahul, Mieneke C. M. Luijendijk, Louk J. M. Vanderschuren, Susanne E. la Fleur, and Roger A. H. Adan. 2014. “Limbic Substrates of the Effects of Neuropeptide Y on Intake of and Motivation for Palatable Food.” Obesity. https://doi.org/10.1002/oby.20718.

Pandit, Rahul, Azar Omrani, Mieneke C. M. Luijendijk, Véronne A. J. de Vrind, Andrea J. Van Rozen, Ralph J. A. Oude Ophuis, Keith Garner, et al. 2016. “Melanocortin 3 Receptor Signaling in Midbrain Dopamine Neurons Increases the Motivation for Food Reward.” Neuropsychopharmacology: Official Publication of the American College of Neuropsychopharmacology 41 (9): 2241–51.

Panksepp, J., and W. W. Beatty. 1980. “Social Deprivation and Play in Rats.” Behavioral and Neural Biology 30 (2): 197–206.

Pan, Tianying, Chuan Jiang, Juan Cheng, Jiang Xie, Xinghui Liu, Wenming Xu, and Guolin He. 2021. “Autism-Like Behavior in the Offspring of CYP11A1-Overexpressing Pregnant Rats.” Frontiers in Neuroscience 15 (December): 774439.

Parker, Nathan F., Courtney M. Cameron, Joshua P. Taliaferro, Junuk Lee, Jung Yoon Choi, Thomas J. Davidson, Nathaniel D. Daw, and Ilana B. Witten. 2016. “Reward and Choice Encoding in Terminals of Midbrain Dopamine Neurons Depends on Striatal Target.” Nature Neuroscience 19 (6): 845–54.

Peña, Catherine Jensen, and Frances A. Champagne. 2015. “Neonatal Overexpression of Estrogen Receptor-α Alters Midbrain Dopamine Neuron Development and Reverses the Effects of Low Maternal Care in Female Offspring.” Developmental Neurobiology 75 (10): 1114–24.

Phillips, Robert A., 3rd, Jennifer J. Tuscher, Samantha L. Black, Emma Andraka, N. Dalton Fitzgerald, Lara Ianov, and Jeremy J. Day. 2022. “An Atlas of Transcriptionally Defined Cell Populations in the Rat Ventral Tegmental Area.” Cell Reports 39 (1): 110616.

Pinto, Shirly, Aaron G. Roseberry, Hongyan Liu, Sabrina Diano, Marya Shanabrough, Xiaoli Cai, Jeffrey M. Friedman, and Tamas L. Horvath. 2004. “Rapid Rewiring of Arcuate Nucleus Feeding Circuits by Leptin.” Science 304 (5667): 110–15.

Plasse, G. van der, R. van Zessen, M. C. M. Luijendijk, H. Erkan, G. D. Stuber, G. M. J. Ramakers, and R. A. H. Adan. 2015. “Modulation of Cue-Induced Firing of Ventral Tegmental Area Dopamine Neurons by Leptin and Ghrelin.” International Journal of Obesity 39 (12): 1742–49.

Pologruto, Thomas A., Bernardo L. Sabatini, and Karel Svoboda. 2003. “ScanImage: Flexible Software for Operating Laser Scanning Microscopes.” Biomedical Engineering Online 2 (May): 13.

Poulin, Jean-Francois, Jian Zou, Janelle Drouin-Ouellet, Kwang-Youn A. Kim, Francesca Cicchetti, and Rajeshwar B. Awatramani. 2014. “Defining Midbrain Dopaminergic Neuron Diversity by Single-Cell Gene Expression Profiling.” Cell Reports 9 (3): 930–43.

Puga, L., V. Alcántara-Alonso, U. Coffeen, O. Jaimes, and P. de Gortari. 2016. “TRH Injected into the Nucleus Accumbens Shell Releases Dopamine and Reduces Feeding Motivation in Rats.” Behavioural Brain Research 306 (June): 128–36.

Remedios, Ryan, Ann Kennedy, Moriel Zelikowsky, Benjamin F. Grewe, Mark J. Schnitzer, and David J. Anderson. 2017. “Social Behaviour Shapes Hypothalamic Neural Ensemble Representations of Conspecific Sex.” Nature 550 (7676): 388–92.

Riediger, Thomas, Martin Traebert, Herbert A. Schmid, Caroline Scheel, Thomas A. Lutz, and Erwin Scharrer. 2003. “Site-Specific Effects of Ghrelin on the Neuronal Activity in the Hypothalamic Arcuate Nucleus.” Neuroscience Letters 341 (2): 151–55.

Rissman, Emilie F., Scott R. Wersinger, Heather N. Fugger, and Thomas C. Foster. 1999. “Sex with Knockout Models: Behavioral Studies of Estrogen Receptor α1Published on the World Wide Web on 26 April 1999.1.” Brain Research 835 (1): 80–90.

Robinson, Donita L., Michael L. A. V. Heien, and R. Mark Wightman. 2002. “Frequency of Dopamine Concentration Transients Increases in Dorsal and Ventral Striatum of Male Rats during Introduction of Conspecifics.” The Journal of Neuroscience: The Official Journal of the Society for Neuroscience 22 (23): 10477–86.

Roepke, Troy A., Anna Malyala, Martha A. Bosch, Martin J. Kelly, and Oline K. Rønnekleiv. 2007. “Estrogen Regulation of Genes Important for K+ Channel Signaling in the Arcuate Nucleus.” Endocrinology 148 (10): 4937–51.

Roselli, C. E., S. A. Klosterman, and T. A. Fasasi. 1996. “Sex Differences in Androgen Responsiveness in the Rat Brain: Regional Differences in the Induction of Aromatase Activity.” Neuroendocrinology 64 (2): 139–45.

Sakurai, T., A. Amemiya, M. Ishii, I. Matsuzaki, R. M. Chemelli, H. Tanaka, S. C. Williams, et al. 1998. “Orexins and Orexin Receptors: A Family of Hypothalamic Neuropeptides and G Protein-Coupled Receptors That Regulate Feeding Behavior.” *Cell*.

Samms, Ricardo J., Kyle W. Sloop, Fiona M. Gribble, Frank Reimann, and Alice E. Adriaenssens. 2021. “GIPR Function in the Central Nervous System: Implications and Novel Perspectives for GIP-Based Therapies in Treating Metabolic Disorders.” Diabetes 70 (9): 1938–44.

Sato, Takashi, Takahiro Matsumoto, Hirotaka Kawano, Tomoyuki Watanabe, Yoshikatsu Uematsu, Keisuke Sekine, Toru Fukuda, et al. 2004. “Brain Masculinization Requires Androgen Receptor Function.” Proceedings of the National Academy of Sciences of the United States of America 101 (6): 1673–78.

Saunders, Arpiar, Evan Z. Macosko, Alec Wysoker, Melissa Goldman, Fenna M. Krienen, Heather de Rivera, Elizabeth Bien, et al. 2018. “Molecular Diversity and Specializations among the Cells of the Adult Mouse Brain.” Cell 174 (4): 1015–30.e16.

Schultz, W., P. Dayan, and P. R. Montague. 1997. “A Neural Substrate of Prediction and Reward.” Science 275 (5306): 1593–99.

Shahrokh, Dara K., Tie-Yuan Zhang, Josie Diorio, Alain Gratton, and Michael J. Meaney. 2010. “Oxytocin-Dopamine Interactions Mediate Variations in Maternal Behavior in the Rat.” Endocrinology 151 (5): 2276–86.

Shen, M., C. Jiang, P. Liu, F. Wang, and L. Ma. 2016. “Mesolimbic Leptin Signaling Negatively Regulates Cocaine-Conditioned Reward.” Translational Psychiatry 6 (12): e972.

Simard, Jacques, Marie-Louise Ricketts, Sébastien Gingras, Penny Soucy, F. Alex Feltus, and Michael H. Melner. 2005. “Molecular Biology of the 3beta-Hydroxysteroid dehydrogenase/delta5-delta4 Isomerase Gene Family.” Endocrine Reviews 26 (4): 525–82.

Simerly, R. B., C. Chang, M. Muramatsu, and L. W. Swanson. 1990. “Distribution of Androgen and Estrogen Receptor mRNA-Containing Cells in the Rat Brain: An in Situ Hybridization Study.” The Journal of Comparative Neurology 294 (1): 76–95.

Sjögren, Maria, Anneke Alkemade, Jens Mittag, Kristina Nordström, Abram Katz, Björn Rozell, Håkan Westerblad, Anders Arner, and Björn Vennström. 2007. “Hypermetabolism in Mice Caused by the Central Action of an Unliganded Thyroid Hormone Receptor alpha1.” The EMBO Journal 26 (21): 4535–45.

Smiley, Kristina O., Rosemary S. E. Brown, and David R. Grattan. 2021. “Prolactin Action Is Necessary for Parental Behavior in Male Mice.” bioRxiv. https://doi.org/10.1101/2021.12.30.474525.

Solié, Clément, Benoit Girard, Beatrice Righetti, Malika Tapparel, and Camilla Bellone. 2021. “VTA Dopamine Neuron Activity Encodes Social Interaction and Promotes Reinforcement Learning through Social Prediction Error.” *Nature Neuroscience*, December. https://doi.org/10.1038/s41593-021-00972-9.

Song, Zhimin, Johnathan M. Borland, Tony E. Larkin, Maureen O’Malley, and H. Elliott Albers. 2016. “Activation of Oxytocin Receptors, but Not Arginine-Vasopressin V1a Receptors, in the Ventral Tegmental Area of Male Syrian Hamsters Is Essential for the Reward-like Properties of Social Interactions.” Psychoneuroendocrinology 74 (December): 164–72.

Steinberg, Elizabeth E., Ronald Keiflin, Josiah R. Boivin, Ilana B. Witten, Karl Deisseroth, and Patricia H. Janak. 2013. “A Causal Link between Prediction Errors, Dopamine Neurons and Learning.” Nature Neuroscience. https://doi.org/10.1038/nn.3413.

Straus, E., and R. S. Yalow. 1979. “Cholecystokinin in the Brains of Obese and Nonobese Mice.” Science 203 (4375): 68–69.

Stuart, Tim, Andrew Butler, Paul Hoffman, Christoph Hafemeister, Efthymia Papalexi, William M. Mauck 3rd, Yuhan Hao, Marlon Stoeckius, Peter Smibert, and Rahul Satija. 2019. “Comprehensive Integration of Single-Cell Data.” Cell 177 (7): 1888–1902.e21.

Tenk, Christine M., Hilary Wilson, Qi Zhang, Kyle K. Pitchers, and Lique M. Coolen. 2009. “Sexual Reward in Male Rats: Effects of Sexual Experience on Conditioned Place Preferences Associated with Ejaculation and Intromissions.” Hormones and Behavior 55 (1): 93–97.

Thompson, Jennifer L., and Stephanie L. Borgland. 2013. “Presynaptic Leptin Action Suppresses Excitatory Synaptic Transmission onto Ventral Tegmental Area Dopamine Neurons.” Biological Psychiatry 73 (9): 860–68.

Tiklová, Katarína, Åsa K. Björklund, Laura Lahti, Alessandro Fiorenzano, Sara Nolbrant, Linda Gillberg, Nikolaos Volakakis, et al. 2019. “Single-Cell RNA Sequencing Reveals Midbrain Dopamine Neuron Diversity Emerging during Mouse Brain Development.” Nature Communications 10 (1): 581.

Tomova, Livia, Kimberly L. Wang, Todd Thompson, Gillian A. Matthews, Atsushi Takahashi, Kay M. Tye, and Rebecca Saxe. 2020. “Acute Social Isolation Evokes Midbrain Craving Responses Similar to Hunger.” Nature Neuroscience 23 (12): 1597–1605.

Trainor, Brian C., Ian M. Bird, Noel A. Alday, Barney A. Schlinger, and Catherine A. Marler. 2003. “Variation in Aromatase Activity in the Medial Preoptic Area and Plasma Progesterone Is Associated with the Onset of Paternal Behavior.” Neuroendocrinology 78 (1): 36–44.

Trainor, Brian C., Helen H. Kyomen, and Catherine A. Marler. 2006. “Estrogenic Encounters: How Interactions between Aromatase and the Environment Modulate Aggression.” Frontiers in Neuroendocrinology 27 (2): 170–79.

Van Segbroeck, Maarten, Allison T. Knoll, Pat Levitt, and Shrikanth Narayanan. 2017. “MUPET-Mouse Ultrasonic Profile ExTraction: A Signal Processing Tool for Rapid and Unsupervised Analysis of Ultrasonic Vocalizations.” Neuron 94 (3): 465–85.e5.

Verharen, J. P. H., Yichen Zhu, and Stephan Lammel. 2020. “Aversion Hot Spots in the Dopamine System.” Current Opinion in Neurobiology 64 (October): 46–52.

V, Hellier, V. Hellier, O. Brock, M. Candlish, E. Desroziers, M. Aoki, C. Mayer, et al. 2018. “Female Sexual Behavior in Mice Is Controlled by Kisspeptin Neurons.” Yearbook of Paediatric Endocrinology. https://doi.org/10.1530/ey.15.1.1.

Walker, Angela K., Imikomobong E. Ibia, and Jeffrey M. Zigman. 2012. “Disruption of Cue-Potentiated Feeding in Mice with Blocked Ghrelin Signaling.” Physiology & Behavior 108 (December): 34–43.

Wardman, Jonathan H., Ivone Gomes, Erin N. Bobeck, Jennifer A. Stockert, Abhijeet Kapoor, Paola Bisignano, Achla Gupta, et al. 2016. “Identification of a Small-Molecule Ligand That Activates the Neuropeptide Receptor GPR171 and Increases Food Intake.” Science Signaling 9 (430): ra55.

Watabe-Uchida, Mitsuko, Lisa Zhu, Sachie K. Ogawa, Archana Vamanrao, and Naoshige Uchida. 2012. “Whole-Brain Mapping of Direct Inputs to Midbrain Dopamine Neurons.” Neuron 74 (5): 858–73.

Weiner, Benjamin, Stav Hertz, Nisim Perets, and Michael London. 2016. “Social Ultrasonic Vocalization in Awake Head-Restrained Mouse.” Frontiers in Behavioral Neuroscience 10 (December): 236.

Wei, Wei, Ali Mohebi, and Joshua D. Berke. 2021. “Striatal Dopamine Pulses Follow a Temporal Discounting Spectrum.” bioRxiv. https://doi.org/10.1101/2021.10.31.466705.

Whitney, G., M. Alpern, G. Dizinno, and G. Horowitz. 1974. “Female Odors Evoke Ultrasounds from Male Mice.” Animal Learning & Behavior 2 (1): 13–18.

Whitten, W. K. 1956. “Modification of the Oestrous Cycle of the Mouse by External Stimuli Associated with the Male.” The Journal of Endocrinology 13 (4): 399–404.

Willmore, Lindsay, Courtney Cameron, John Yang, Ilana Witten, and Annegret Falkner. 2022. “Behavioral and Dopaminergic Signatures of Resilience.” bioRxiv. https://doi.org/10.1101/2022.03.18.484885.

Woods, S. C., E. C. Lotter, L. D. McKay, and D. Porte Jr. 1979. “Chronic Intracerebroventricular Infusion of Insulin Reduces Food Intake and Body Weight of Baboons.” Nature 282 (5738): 503–5.

Xiao, Lei, Michael F. Priest, Jordan Nasenbeny, Ting Lu, and Yevgenia Kozorovitskiy. 2017. “Biased Oxytocinergic Modulation of Midbrain Dopamine Systems.” Neuron 95 (2): 368–84.e5.

Young, Larry J. 2002. “The Neurobiology of Social Recognition, Approach, and Avoidance.” Biological Psychiatry 51 (1): 18–26.

Young, L. J., R. Nilsen, K. G. Waymire, G. R. MacGregor, and T. R. Insel. 1999. “Increased Affiliative Response to Vasopressin in Mice Expressing the V1a Receptor from a Monogamous Vole.” Nature 400 (6746): 766–68.

Zhang, Qian, Challa Tenagne Delessa, Robert Augustin, Mostafa Bakhti, Gustav Colldén, Daniel J. Drucker, Annette Feuchtinger, et al. 2021. “The Glucose-Dependent Insulinotropic Polypeptide (GIP) Regulates Body Weight and Food Intake via CNS-GIPR Signaling.” Cell Metabolism 33 (4): 833–44.e5.

Zhang, Stephen X., Andrew Lutas, Shang Yang, Adriana Diaz, Hugo Fluhr, Georg Nagel, Shiqiang Gao, and Mark L. Andermann. 2021. “Hypothalamic Dopamine Neurons Motivate Mating through Persistent cAMP Signalling.” Nature 597 (7875): 245–49.

Zheng, Danielle, Soledad Cabeza de Vaca, and Kenneth D. Carr. 2012. “Food Restriction Increases Acquisition, Persistence and Drug Prime-Induced Expression of a Cocaine-Conditioned Place Preference in Rats.” Pharmacology, Biochemistry, and Behavior 100 (3): 538–44.

Zigman, Jeffrey M., Juli E. Jones, Charlotte E. Lee, Clifford B. Saper, and Joel K. Elmquist. 2006. “Expression of Ghrelin Receptor mRNA in the Rat and the Mouse Brain.” The Journal of Comparative Neurology 494 (3): 528–48.

Zolin, Aryeh, Raphael Cohn, Rich Pang, Andrew F. Siliciano, Adrienne L. Fairhall, and Vanessa Ruta. 2021. “Context-Dependent Representations of Movement in Drosophila Dopaminergic Reinforcement Pathways.” Nature Neuroscience 24 (11): 1555–66.

